# Perinatal selective serotonin reuptake inhibitor exposure and behavioral outcomes: a systematic review and meta-analyses of animal studies

**DOI:** 10.1101/868265

**Authors:** A.S. Ramsteijn, L. Van de Wijer, J. Rando, J. van Luijk, J.R. Homberg, J.D.A. Olivier

## Abstract

In the Western world, 2-5% of pregnant women use selective serotonin reuptake inhibitor (SSRI) antidepressants. There is no consensus on the potential long-term neurodevelopmental outcomes of early SSRI exposure. Our aim was to determine whether there is an overall effect of perinatal SSRI exposure in animals on a spectrum of behavioral domains. After a comprehensive database search in PubMed, PsycINFO, and Web of Science, we included 99 publications. We performed nine meta-analyses and two qualitative syntheses corresponding to different behavioral categories, aggregating data from thousands of animals. We found evidence for reduced activity and exploration behavior (standardized mean difference (SMD) −0.28 [-0.38, −0.18]), more passive stress coping (SMD −0.37 [-0.52, −0.23]), and less efficient sensory processing (SMD −0.37 [-0.69, −0.06]) in SSRI-versus vehicle-exposed animals. No differences were found for anxiety (p=0.06), social behavior, learning and memory, ingestive- and reward behavior, motoric behavior, or reflex and pain sensitivity. Exposure in the period equivalent to the human third trimester was associated with the strongest effects.

**Highlights:**

- Perinatal SSRI exposure in rodents alters outcomes in three behavioral domains.
- It leads to reduced activity, passive stress coping, and weaker sensory processing.
- Females are understudied but seem to be less vulnerable than males.
- Early postnatal exposure in rodents leads to the largest effects on behavior.
- This is equivalent to the third trimester of pregnancy in humans.

## 1. Introduction

Selective serotonin reuptake inhibitor (SSRI) antidepressant use during pregnancy has increased tremendously over the past decades^1–4^. Recent estimates of SSRI exposure in large population-based studies range from 2.5-3.3% of pregnancies in Europe ^5, 6^ to 2.7-5.4% in the US ^7, 8^. These numbers imply that every year, in these regions alone, hundreds of thousands of babies are born after exposure to SSRIs. Although major teratogenic effects are absent, in utero SSRI exposure has been associated with increased risk of neonatal complications such as preterm birth^9^. SSRIs reach the developing fetus by crossing the placental barrier^10^. During fetal development, the serotonin transporter (SERT), the target of SSRIs, is much more diffusely expressed in the brain than during adulthood^11^. In fact, the entire serotonergic neurotransmitter system functions differently in adulthood than during development. In adulthood, serotonin is involved in fundamental brain functions such as the regulation of mood, sleep and wake rhythms, aggression, appetite, learning and memory, and reward^12^, while during early development, serotonin serves as a neurotrophic factor mediating basic processes such as neurogenesis, cell migration, axon guidance, dendritogenesis and synaptogenesis^13^. Consequently, by reaching the brain and modulating serotonin regulation at crucial neurodevelopmental stages, SSRIs could interfere with brain circuit formation and lifelong mental health^14^.

This is the rationale for the “SSRI paradox”, which refers to the phenomenon in which adult SSRI exposure decreases symptoms of anxiety and depression, while in utero SSRI exposure increases the risk of developing anxiety and depression^15^. There is mixed evidence for this theory from human studies, which do not always identify long-lasting neurodevelopmental effects of perinatal SSRI exposure. On the one hand, studies have reported higher levels of anxiety^16^ and lower scores on motor-, social-emotional- and adaptive behavioral tests^17^ after prenatal SSRI exposure. On the other hand, other studies found no association between in utero SSRI exposure and intellectual disability^18^, executive functioning^19^, and emotional or social problems^20^. Most of the evidence is obtained from studies in infants and children, likely due to the practical challenges of examining the effects of in utero exposure to SSRIs on behavioral outcomes in adulthood^21^. Interestingly, some of the reported associations are modulated by behavioral outcome domain^22, 23^, timing of exposure^20^, and sex ^23, 24^. Summarizing the available evidence, a recent meta-analysis reported significant positive associations between SSRI exposure during pregnancy and the development of mental and behavioral disorders such as autism spectrum disorder (ASD), attention-deficit/hyperactivity disorder (ADHD), and mental disability^25^. As these results may be confounded by factors such as the severity of mental health problems, it remains difficult to draw conclusions on causality^25^. Indeed, it is known that maternal mental health issues during pregnancy are associated with long-term neurodevelopmental outcomes in children as well^26^.

Laboratory rodents mature much faster than humans, yet the sequence of brain developmental milestones is remarkably similar^27^. In contrast to human studies, experimental studies in laboratory animals allow for investigation of the causal relationship between perinatal SSRI exposure and long-term neurodevelopmental outcomes^28^. Animal experiments have several other advantages, such as the ability to study the developmental effects of SSRI treatment during a healthy pregnancy and a high degree of control over drug dosing and period of exposure. The last decade especially has witnessed a major surge in animal studies examining various neurobiological outcomes of perinatal SSRI exposure, which have been described in numerous narrative reviews^14, 29–34^. To maximize the translational value of animal studies, and in line with efforts to reduce the use of animals in research, it is imperative to comprehensively bundle all available preclinical evidence. Our aim is to systematically review and analyze preclinical studies in order to determine whether there is an overall effect of perinatal SSRI exposure on later-life behavior in animal models, and if so, under what conditions. We particularly focused on potential sex differences, interactions with stress exposure, and the timing of SSRI exposure. The results of this review and accompanying meta-analyses may assist in understanding the mixed results of perinatal SSRI exposure in human studies and help inform future study design.

## 2. Methods

The review protocol was registered at the SYRCLE website (www.syrcle.nl) in 2016. The reporting in this systematic review adheres to the Preferred Reporting Items for Systematic Reviews and Meta-Analyses (PRISMA) statement^35^.

### 2.1. Search strategy

Three databases were searched systematically from inception to February 27^th^ 2018: PubMed, PsycINFO, and Web of Science. The initial search was performed by JR on April 19^th^ 2016. An updated search was performed by AR on February 27^th^ 2018. We searched for the following concepts, using both controlled terms (i.e. MeSH) and free text words: (i) perinatal exposure; (ii) selective serotonin reuptake inhibitor ^36^ (SSRI); (iii) animal (Supplementary File 1). The SYRCLE animal filter was used for PubMed and adapted for PsycINFO and Web of Science. The bibliographic records retrieved were imported and de-duplicated in Mendeley.

### 2.2. Eligibility screening

Studies were eligible for inclusion if they compared behavioral outcomes of animals perinatally exposed to SSRIs to those of animals exposed to a vehicle treatment. Two reviewers independently screened all identified records for eligibility in two stages using EROS 3.0 (Early Review Organizing Software, Institute of Clinical Effectiveness and Health Policy, Buenos Aires, Argentina). JR and LW performed the screening for the articles identified in the initial search, and AR and LW for those identified in the updated search. Disagreements were resolved by discussion.

The first screening stage involved screening only the title and abstract of the articles. Articles were excluded for one or more of the following reasons: (i) not an original primary study (e.g. review, editorial, conference abstract without full data available) or correction to an original primary study; (ii) not an in vivo mammalian (non-human) study; (iii) no SSRI treatment.

In the second stage, the full text of all articles passing the first stage was consulted. Articles were excluded at this stage for one or more of the following reasons: (i) not an original primary study (e.g., review, editorial, conference abstract without full data available or data published in duplicate) or correction to an original primary study; (ii) not an in vivo mammalian (non-human) study; (iii) no SSRI treatment; (iv) no exposure on or before the developmental day equivalent to human birth in terms of neurogenesis, GABA cortex development, and axon extension, calculated using the Translating Time tool developed by Workman et al^37^: PND11 in mice and PND10 in rats; (v) no behavior analyses; (vi) no control population; (vii) animals subjected to other factors (e.g., genetic mutation, repeated exposure to additional drug), but studies in which animals or their mothers were exposed to stress were included because these studies are translationally relevant; (viii) no repeated exposure; (ix) no English full text or translation available.

### 2.3. Extraction of study characteristics and data

The following study characteristics were extracted: (i) study ID: authors, year, title; (ii) study design characteristics: no. of groups, no. of animals per group, no. of litters per group, litter size, repeated measures vs. comparison between groups; (iii) animal model characteristics: species, strain, sex, age at testing, presence/absence of stress exposure; (iv) intervention characteristics: type of control, type of SSRI, age and duration of exposure, administration method, dosage (concentration, volume of administration); (v) outcome measures: behavioral test used, test outcome; (vi) other: no. of animals excluded from statistical analysis, reason for excluding animals.

Then, the data from all behavioral outcomes were extracted: means, standard deviation (SD) or standard error of the mean (SEM) and number of animals (N). The methods for extraction were, in order of priority, (i) extract data from text or tables; (ii) extract data from figures using digital image analysis software (ImageJ v. 1.52a^38^); (iii) contact authors for missing data. When SDs/SEMs were not clearly distinguishable in a figure, we extracted the most conservative estimate. JR performed the data extraction for all eligible articles retrieved in the initial search, and AR for those in the updated search. LW checked the extraction process for all studies.

### 2.4. Data analysis

#### 2.4.1. Categorization of behavioral tests

After the data extraction, all behavioral tests found were categorized by AR in consultation with JH and JO and other members of the Behavioral Neuroscience group at the University of Groningen. Ten categories were defined – in order of number of comparisons: (i) activity & exploration; (ii) anxiety; (iii) stress coping; (iv) social behavior; (v) learning & memory; (vi) ingestive & reward; (vii) motoric; (viii) sensory processing; (ix) reflex & pain sensitivity; (x) sleep & circadian activity. Every category had a number of behavioral tests associated with it (Supplementary File 2). For every behavioral category we performed a meta-analysis. An exception was the category sleep & circadian activity, which was deemed too heterogeneous and more suitable for a qualitative synthesis. There was an eleventh category of behavioral tests, in which the animals were challenged with an acute injection of a drug or LPS right before the test. To ensure the analyses for the above-mentioned behavioral categories were not confounded by the effects of an acute injection, we decided not to include these results in any of the 10 categories, and to create a separate qualitative synthesis for them.

#### 2.4.2. Selection of comparisons

If a study reported separate comparisons for males and females, or animals exposed to different SSRIs, we analyzed these comparisons as if they were separate studies. Per meta-analysis, one unique animal can only be used once. If the same animal was exposed to different behavioral tests within the same category, we used the data from the test that was performed first (but when data was available from both during and after SSRI exposure, we used the data from the test performed after SSRI exposure). If the same animal was exposed to the same behavioral tests multiple times, we also used the data from the first time it was administered, unless the test contained an important learning or habituation component. For that reason, the data from the last time of test administration was used for the following behavioral tests: alcohol consumption, cocaine conditioning, forced swim test, Morris water maze, sexual behavior, sucrose preference test, and tube runway. In the prepulse inhibition test, usually a range of pulse intensities was tested, in which case we used the data from the middle intensity. For every behavioral test, we only used one outcome measure according to the priority outcome measures we defined (Supplemental File 2). We did not include non-treated or non-handled controls; only vehicle-treated controls.

#### 2.4.3. Meta-analyses

We performed the meta-analyses using Review Manager (RevMan v.5.3.,The Nordic Cochrane Centre, The Cochrane Collaboration, Copenhagen 2014). When a range was reported for N, instead of a specific number per treatment group (for instance N=11-13), we used the most conservative estimate of N. In practice, this meant we used the maximum value of N (in this case N_max_=13) to calculate the SD (SD=SEM*√N), and the minimum value of N (in this case N_min_=11) in the actual meta-analysis. We used random effects models using standardized mean differences (SMDs). The individual SMDs were pooled ^2^ to obtain an overall SMD and 95% confidence interval (CI). I was used as a measure of heterogeneity. A p-value lower than 0.05 was considered significant.

To examine potential sources of heterogeneity within the data, we performed subgroup analyses ^2^ using a Chi test for subgroup differences based on sex, presence/absence of stress exposure, and period of SSRI exposure for every meta-analysis. For the subgroup analysis for sex there were three subgroups (male, mixed-sex, and female), for presence/absence of stress exposure there were two (no stress and stress) and for period of SSRI exposure three (prenatal, pre- and postnatal, and postnatal). A subgroup analysis was only performed when there was at least one independent comparison. Although there were six subgroup analyses defined in the initial published protocol, we decided to only perform three in order to constrain the scope of this review. We decided not to perform subgroup analyses based on animal species, timing of behavioral test, type of SSRI, and specific behavioral test used. Of the three subgroup analyses we performed, two were included in the original protocol (sex and presence/absence of stress exposure) and one was added (period of SSRI exposure).

### 2.5. Risk of bias assessment

To assess the methodological quality of each included study, we used the SYRCLE risk of bias tool for animal studies^39^. We added three questions on reporting of randomization, blinding, and a power- or sample size calculation (question 1-3). For these questions, a “Yes” score indicates that it was reported, and a “No” score indicates that it was not reported. The other questions (question 4-14) addressed risk of bias, where “Yes” indicates low risk of bias, “?” indicates unclear risk of bias, and “No” indicates high risk of bias.

### 2.6. Publication bias assessment

To assess publication bias, funnel plots were produced for each of the nine meta-analyses using the ^40^ package “metafor” v2.1-0 in R v3.5. Each funnel plot displays all studies in one plot with SMD as the x-value and 1/√N as the y-value. We used this method because it was shown that plotting the SMD against the SE can lead to false-positive results, especially when the included studies have small sample sizes^41^. In the funnel plot, larger studies with high precision and power will be displayed towards the top of the graph, around the average SMD. In the absence of publication bias, smaller studies with lower precision and power will spread evenly on both sides of the average near the bottom of the graph. If the plot is asymmetrical, for example when smaller studies predominantly have SMDs larger than the average, this is an indication of small-study bias, potentially related to publication bias. To test and adjust for funnel ^42^ plot asymmetry, we used the trim and fill method in the “metafor” package.

## 3. Results

### 3.1. Search results

Through database searching, 5951 records were retrieved, leaving 3930 records after removal of duplicates (Figure 1). After screening by title and abstract, 1460 full-text articles were assessed for eligibility, from which 103 were deemed eligible. After adding one extra article identified by scanning of the reference lists of the included articles, and excluding five publications because they did not contain usable data, we finally included 99 publications in this synthesis of evidence (Figure 1).

**Figure 1:**
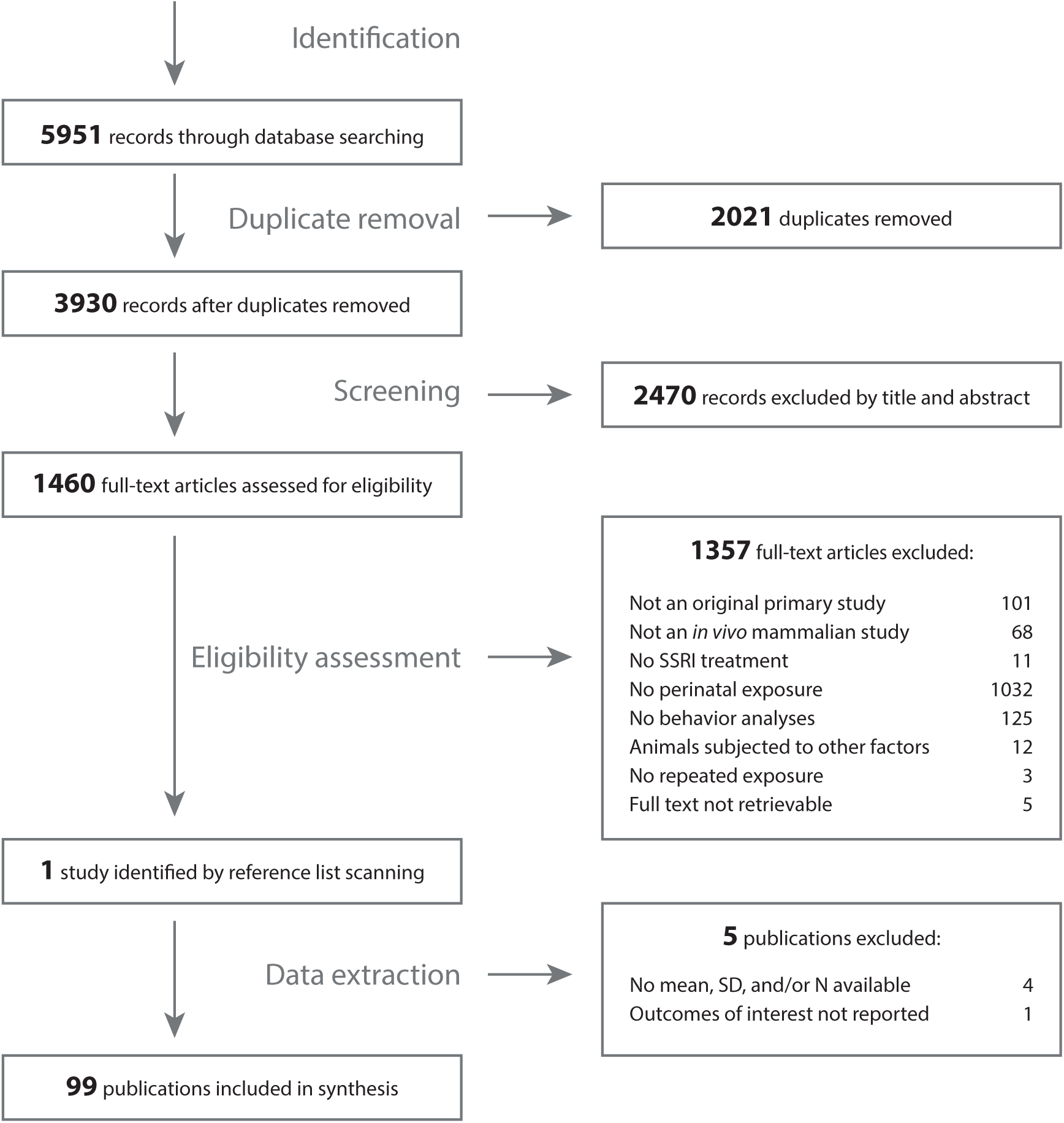
Study flowchart.

### 3.2. Study characteristics

From the 99 included publications, 63 studied rats, 35 mice and one guinea pigs (Table 1). The majority of studies treated animals with fluoxetine (67 studies), followed by citalopram (15 studies), zimelidine (eight studies), escitalopram (five studies), sertraline (four studies), fluvoxamine (three studies), paroxetine (three studies), and LU 10-134-C (one study) (Supplementary Figure 1A). SSRI exposure was prenatal in 18 studies, both prenatal and postnatal in 23 studies, and postnatal in 59 studies. From the studies where SSRIs were administered postnatally (either exclusively, or also prenatally), 54 reported injecting the drug directly into the pups, and 28 reported exposure through the mother. The method of SSRI administration was subcutaneous in 43 studies, oral in 31 studies, and intraperitoneal in 25 studies. Forty-seven studies tested male rats, seven studies female, and 45 studies examined both sexes (Supplementary Figure 1B). Please note that study numbers might add up to more than 99, because the same study could use multiple SSRIs or exposure periods (Table 1).

**Table 1:**
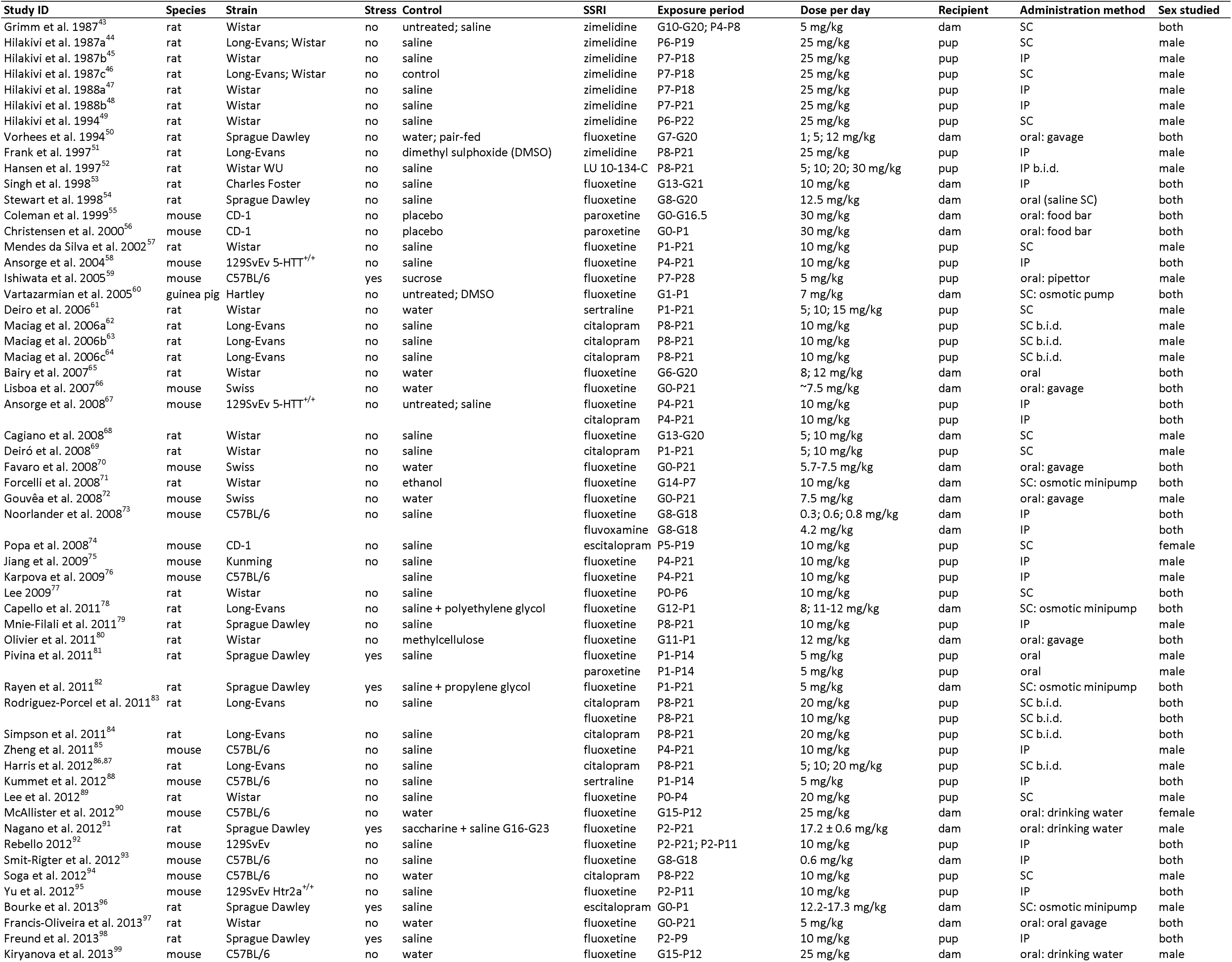
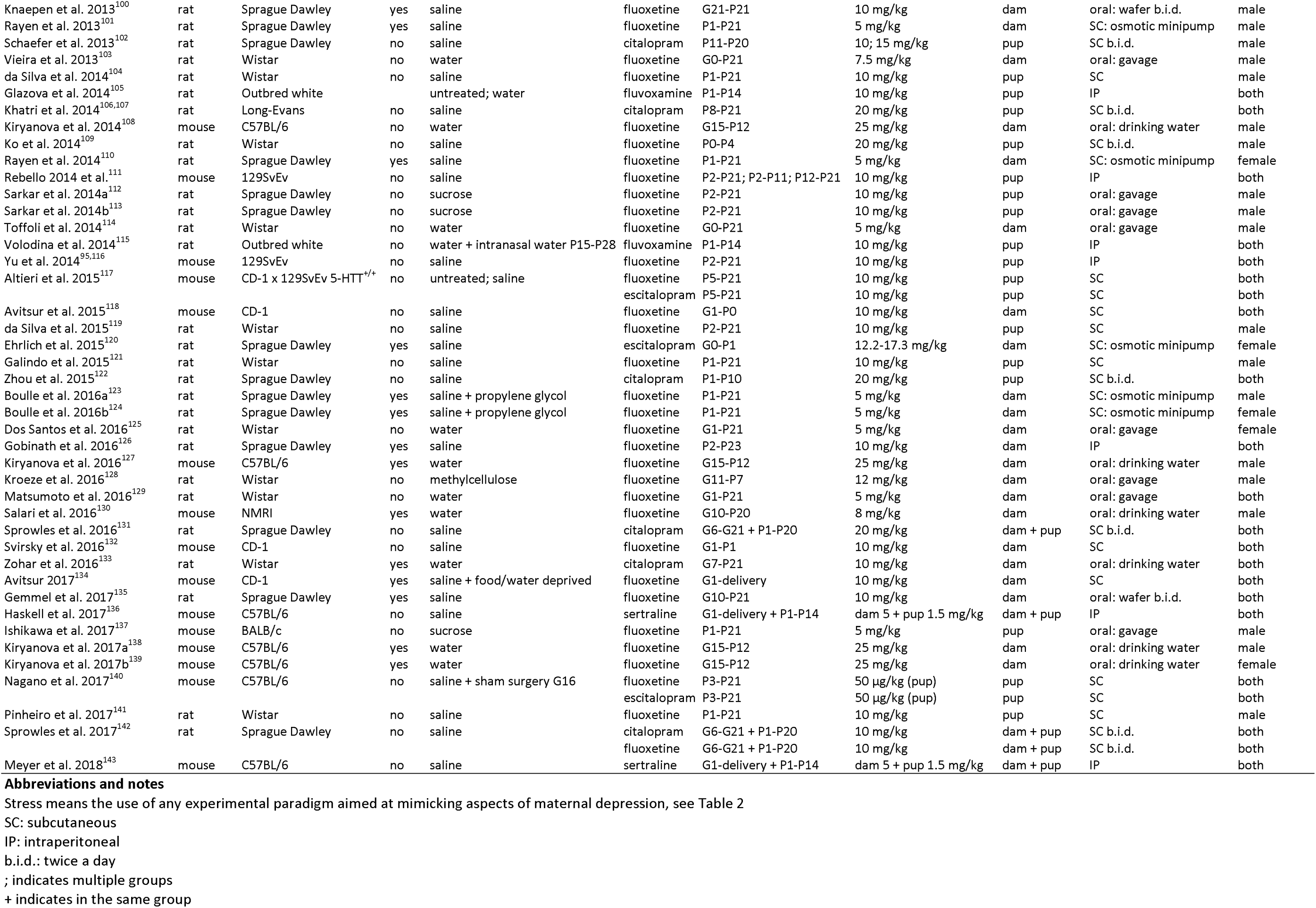
study characteristics

Twenty studies used ways to mimic symptoms associated with maternal depression in laboratory animals (Table 2). In 19 studies, the dam was exposed to some form of stress, and in one study the pups were stressed by means of maternal separation. The most common way to apply stress to the mother was using repeated restraint stress (10 studies), followed by chronic unpredictable mild stress (seven studies), and injections of corticosterone or dexamethasone (one study each).

**Table 2:**
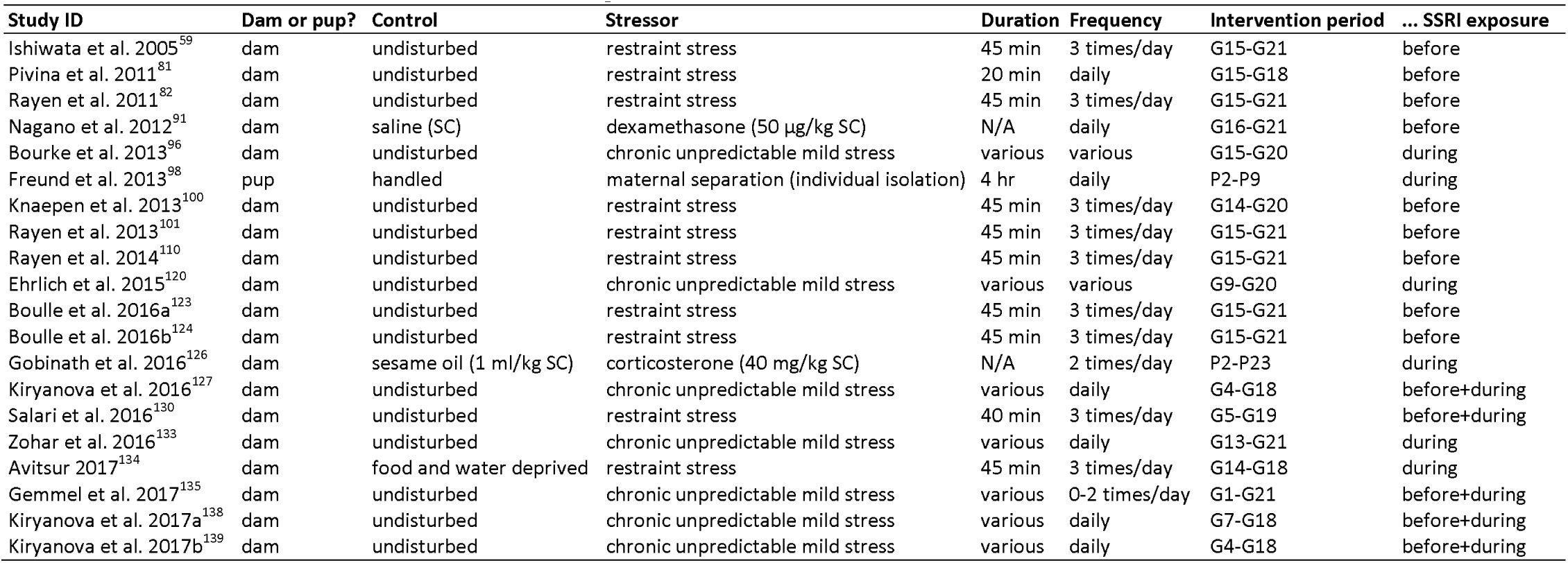
characteristics of studies combining (maternal) stress with SSRI treatment

### 3.3. Study quality

Forty-eight studies mentioned the experiment was randomized at some level, 31 reported blinding, and three included a power or sample size calculation (Table 3). Overall risk of bias was unclear. Only 68 studies reported all outcome measures that were described in the methods section.

**Table 3:**
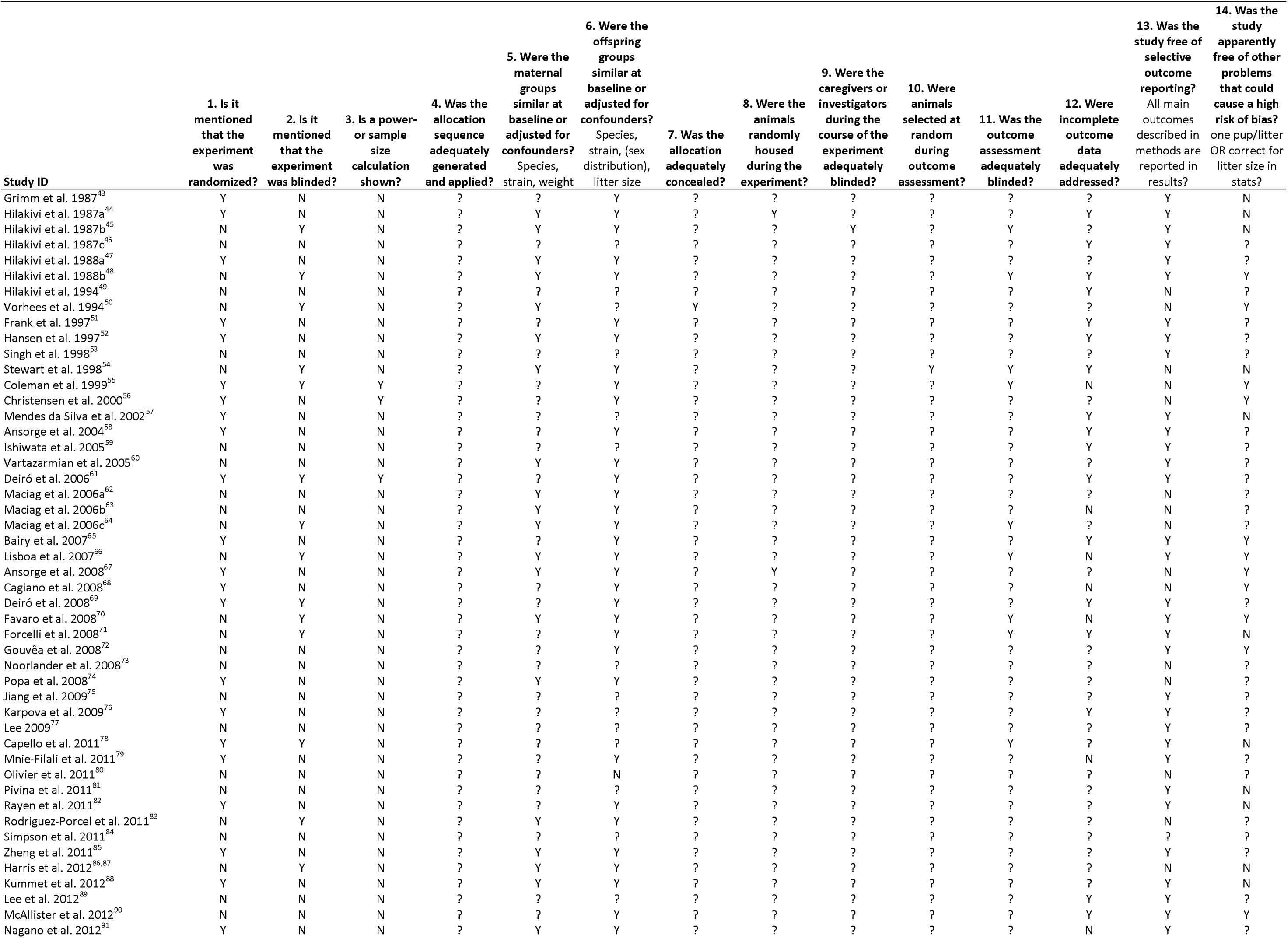
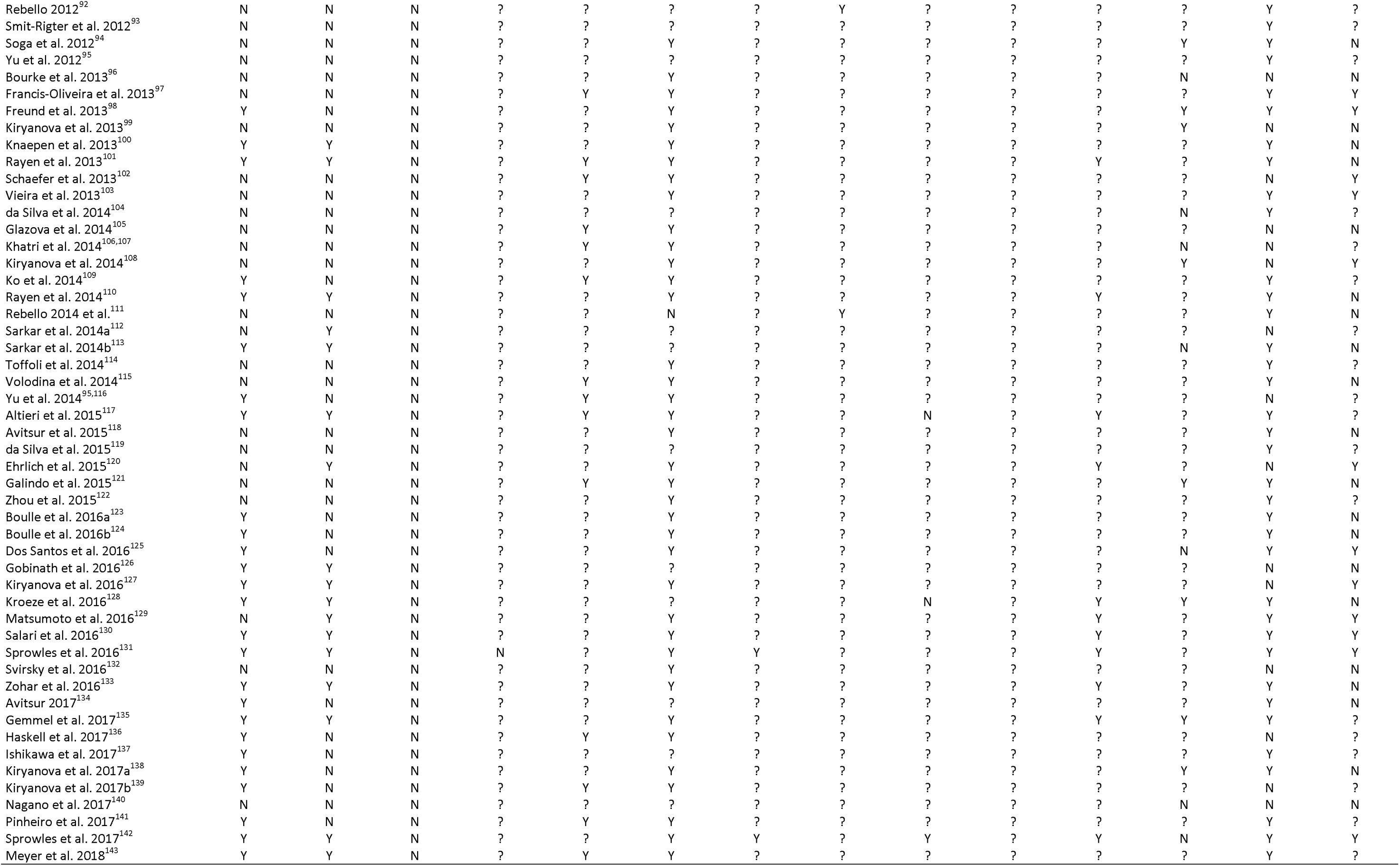
Risk of bias results

### 3.4. Activity and exploration

The meta-analysis for activity and exploration comprised 52 studies and 134 comparisons. The most used behavioral test in this category was the open field test with outcome measures such as total distance moved (121 comparisons), followed by the novel object exploration test (six comparisons), running wheel activity (three comparisons), elevated plus maze (two comparisons), home cage activity (one comparison), and object-directed behavior/novel object recognition test (one comparison). In total, 2646 SSRI-exposed animals and 1627 vehicle-treated animals were included in this analysis.

Overall pooled analysis revealed significantly lower activity scores in animals that were developmentally exposed to SSRIs than in those exposed to vehicle (Figure 2A; Supplementary Figure 2A, SMD −0.28 [-0.38, −0.18], p<0.00001). Subgroup analysis showed that the effect was different depending on sex (Figure 2A; Supplementary Figure 2B, Chi²=13.89, p<0.01). More specifically, while activity scores were significantly lower for males (SMD −0.28 [-0.41, −0.15], p<0.0001) and mixed-sex groups (SMD −0.62 [-0.82, −0.42], p<0.00001) developmentally exposed to SSRIs versus those exposed to vehicle, they were not for females (SMD −0.12 [-0.29, 0.04], p=0.14) (Figure 2A; Supplementary Figure 2B). Subgroup analysis based on stress exposure did not reveal significantly different effects of developmental SSRI exposure depending on stress exposure (Figure 2A; Supplementary Figure 2C, Chi²=1.76, p=0.18). Subgroup analysis based on the period of SSRI exposure showed that the effect of developmental SSRI exposure on later-life activity and exploration was different depending on exposure timing (Figure 2A; Supplementary Figure 2D, Chi²=11.60, p<0.01). More specifically, while activity scores were not different for those exposed only prenatally (SMD −0.01 [-0.21, 0.19], p=0.93), they were significantly lower for animals exposed pre- and postnatally (SMD −0.40 [-0.59, −0.22], p<0.0001), and postnatally (SMD −0.39 [-0.51, - 0.27], p<0.00001) versus those exposed to vehicle (Figure 2A; Supplementary Figure 2D).

**Figure 2:**
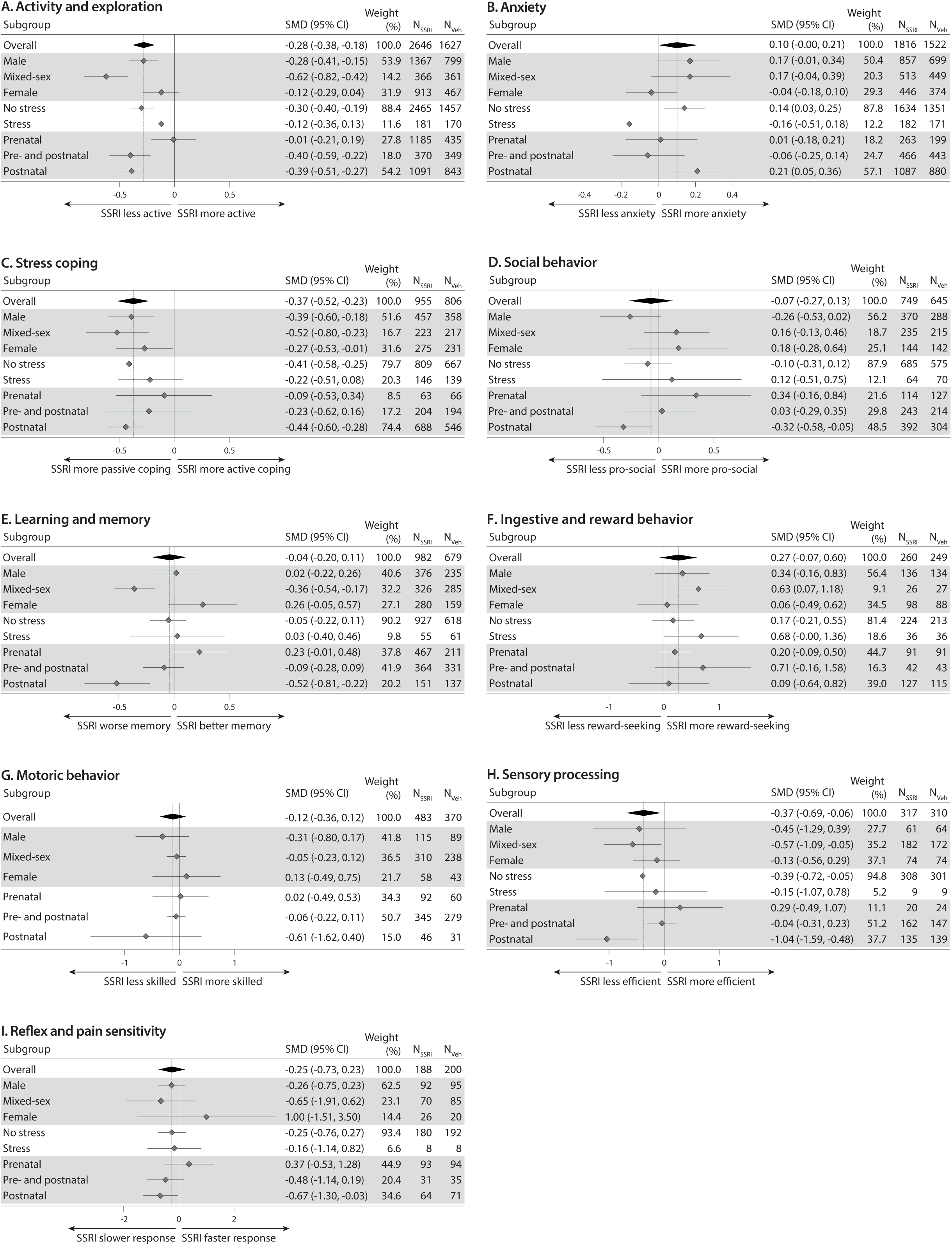
Summary forest plots from all meta-analyses comparing animals perinatally exposed to SSRIs to those exposed to vehicle. (A) Activity and exploration. (B) Anxiety. (C) Stress coping. (D) Social behavior. (E) Learning and memory. (F) Ingestive and reward. (G) Motoric behavior. (H) Sensory processing. (I) Reflex and pain sensitivity.

The heterogeneity (I^2^) of the overall analysis was 49%. Subgroup analyses based on sex decreased the heterogeneity to 44% for males, 39% for mixed-sex, and 46% for females. The subgroups based on stress exposure and SSRI exposure timing did not lower the heterogeneity.

### 3.5. Anxiety

The meta-analysis for anxiety comprised 55 studies and 133 comparisons. The most used behavioral test in this category was the open field test with outcome measures such as time spent in center (55 comparisons), followed by the elevated plus maze (46 comparisons), the novelty-suppressed feeding test (11 comparisons), fear during tone (nine comparisons), the defensive withdrawal test (six comparisons), the elevated zero maze (four comparisons), and the light-dark test (two comparisons). In total, 1816 SSRI-exposed animals and 1522 vehicle-treated animals were included in this analysis.

Overall pooled analysis did not show significantly different anxiety scores in animals that were developmentally exposed to SSRIs than in those exposed to vehicle (Figure 2B; Supplementary Figure 3A, SMD 0.10 [-0.00, 0.21], p=0.06). Subgroup analyses did not reveal significantly different effects of developmental SSRI exposure depending on sex (Figure 2B; Supplementary Figure 3B, Chi² = 4.44, p=0.11), stress exposure (Figure 2B; Supplementary Figure 3C, Chi² = 2.73, p=0.10), or period of SSRI exposure (Figure 2B; Supplementary Figure 3D, Chi² = 4.95, p=0.08).

The heterogeneity (I^2^) of the overall analysis was 51%. The subgroups based on sex, stress exposure and SSRI exposure timing did not lower the heterogeneity.

### 3.6. Stress coping

The meta-analysis for stress coping comprised 30 studies and 90 comparisons. The most used behavioral test in this category was the forced swim test (55 comparisons), followed by shock avoidance (30 comparisons), the open field test after stress and the tail suspension test (two comparisons each), and the elevated plus maze after stress (one comparison). In total, 955 SSRI-exposed animals and 806 vehicle-treated animals were included in this analysis.

Overall pooled analysis showed a significantly more passive coping style in animals that were developmentally exposed to SSRIs than in those exposed to vehicle (Figure 2C; Supplementary Figure 4A, SMD −0.37 [-0.52, −0.23], p<0.00001). Subgroup analyses did not reveal significantly different effects of developmental SSRI exposure depending on sex (Figure 2C; Supplementary Figure 4B, Chi² = 1.61, p=0.45), stress exposure (Figure 2C; Supplementary Figure 4C, Chi² = 1.32, p=0.25), or period of SSRI exposure (Figure 2C; Supplementary Figure 4D, Chi² = 2.72, p=0.26).

The heterogeneity (I^2^) of the overall analysis was 48%. The subgroups based on sex, stress exposure and SSRI exposure timing did not lower the heterogeneity.

### 3.7. Social behavior

The meta-analysis for social behavior comprised 30 studies with 53 comparisons. The most used behavioral tests in this category were sexual behavior and social play behavior (14 comparisons each), followed by the social interaction test (10 comparisons), the social preference test (five comparisons), the resident-intruder test (four comparisons), ultrasonic vocalizations (three comparisons), aggressive behavior (two comparisons) and maternal behavior (one comparison). In total, 749 SSRI-exposed animals and 645 vehicle-treated animals were included in this analysis.

Overall pooled analysis did not show significantly different social behavior in animals that were developmentally exposed to SSRIs than in those exposed to vehicle (Figure 2D; Supplementary Figure 5A, SMD −0.07 [-0.27, 0.13], p=0.47). Whereas subgroup analyses did not show significantly different effects of developmental SSRI exposure depending on sex (Figure 2D; Supplementary Figure 5B, Chi² = 5.12, p=0.08) and stress exposure (Figure 2D; Supplementary Figure 5C, Chi² = 0.41, p=0.52), the effect was different depending on period of SSRI exposure (Figure 2D; Supplementary Figure 5D, Chi² = 6.20, p<0.05). More specifically, while SSRI-exposed offspring did not differ in social behavior in those exposed prenatally (SMD 0.34 [-0.16, 0.84], p=0.18) and pre- and postnatally (SMD 0.03 [-0.29, 0.35], p=0.85), animals exposed to SSRIs postnatally were significantly less pro-social than those exposed to vehicle (SMD −0.32 [-0.58, −0.05], p<0.05) (Figure 2D; Supplementary Figure 5D).

The heterogeneity (I^2^) of the overall analysis was 65%. The subgroups based on sex, stress exposure and SSRI exposure timing did not lower the heterogeneity.

### 3.8. Learning and memory

The meta-analysis for learning and memory comprised 23 studies with 47 comparisons. The most used behavioral test in this category was the Morris water maze (18 comparisons), followed by the passive avoidance test (eight comparisons), novel object recognition (seven comparisons), the Cincinnati water maze (five comparisons), contextual fear conditioning (three comparisons), the radial water maze (two comparisons) and the Barnes maze, complex maze, cued fear conditioning and novel scent recognition (one comparison each). In total, 982 SSRI-exposed animals and 679 vehicle-treated animals were included in this analysis.

Overall pooled analysis did not show significantly different learning and memory in animals that were developmentally exposed to SSRIs than in those exposed to vehicle (Figure 2E; Supplementary Figure 6A, SMD −0.04 [-0.20, 0.11], p=0.57). Subgroup analyses revealed significantly different effects of developmental SSRI exposure depending on sex (Figure 2E; Supplementary Figure 6B, Chi² = 13.54, p<0.01). More specifically, the mixed-sex subgroup showed a significantly lower score on learning and memory tests (SMD −0.36 [-0.54, −0.17], p<0.001), but this was not the case for the groups consisting of only males (SMD 0.02 [-0.22, 0.26], p=0.86) or females (SMD 0.26 [-0.05, 0.57], p=0.10) (Figure 2E; Supplementary Figure 6B). There was no different effect of developmental SSRI exposure on learning and memory outcomes depending on stress exposure (Figure 2E; Supplementary Figure 6C, Chi² = 0.13, p=0.72). In contrast, the effect was different depending on period of SSRI exposure (Figure 2E; Supplementary Figure 6D, Chi² = 14.79, p<0.001). More specifically, while SSRI-exposed offspring did not differ significantly in learning and memory outcomes in the groups exposed prenatally (SMD 0.23 [-0.01, 0.48], p=0.06) and pre- and postnatally (SMD −0.09 [-0.28, 0.09], p=0.33), animals exposed to SSRIs postnatally scored significantly lower on learning and memory tests than those exposed to vehicle (SMD −0.52 [-0.81, −0.22], p<0.001) (Figure 2E; Supplementary Figure 6D).

The heterogeneity (I^2^) of the overall analysis was 49%. Subgroup analyses based on sex lowered the heterogeneity to 43% for males, 15% for mixed-sex, and 48% for females. The subgroups based on stress exposure did not lower the heterogeneity. Subgroup analyses based on SSRI exposure timing lowered the heterogeneity to 42% for those exposed prenatally, 27% for those exposed pre- and postnatally, and 28% for those exposed postnatally.

### 3.9. Ingestive- and reward behavior

The meta-analysis for ingestive- and reward behavior comprised 14 studies with 24 comparisons. The most used behavioral test in this category was food consumption (13 comparisons), followed by the sucrose preference test (four comparisons), alcohol consumption, cocaine place preference, and the tube runway (two comparisons each), and cocaine self-administration (one comparison). In total, SSRI-exposed animals and vehicle-treated animals were included in this analysis.

Overall pooled analysis did not show significantly different ingestive- and reward behavior in animals that were developmentally exposed to SSRIs than in those exposed to vehicle (Figure 2F; Supplementary Figure 7A, SMD 0.27 [-0.07, 0.60], p=0.12). Subgroup analyses did not show significantly different effects of developmental SSRI exposure depending on sex (Figure 2F; Supplementary Figure 7B, Chi² = 1.98, p=0.37), stress exposure (Figure 2F; Supplementary Figure 7C, Chi² = 1.65, p=0.20), or period of SSRI exposure (Figure 2F; Supplementary Figure 7D, Chi² = 1.33, p=0.52).

The heterogeneity (I^2^) of the overall analysis was 69%. The subgroups based on sex, stress exposure and SSRI exposure timing did not lower the heterogeneity.

### 3.10. Motoric behavior

The meta-analysis for motoric behavior comprised 11 studies with 20 comparisons. The most used behavioral test in this category was swimming (seven comparisons), followed by beam traversing and the rotarod test (five comparisons each), the horizontal ladder test (two comparisons), and walking (one comparison). In total, 483 SSRI-exposed animals and 370 vehicle-treated animals were included in this analysis.

Overall pooled analysis did not show significantly different motoric behavior in animals that were developmentally exposed to SSRIs than in those exposed to vehicle (Figure 2G; Supplementary Figure 8A, SMD −0.12 [-0.36, 0.12], p=0.50). Subgroup analyses did not show significantly different effects of developmental SSRI exposure depending on sex (Figure 2G; Supplementary Figure 8B, Chi² = 1.40, p=0.50) or period of SSRI exposure (Figure 2G; Supplementary Figure 8C, Chi² = 1.24, p=0.54). Subgroup analysis based on stress exposure could not be done because there were no studies with stress exposure in this category.

The heterogeneity (I^2^) of the overall analysis was 49%. The subgroups based on sex and SSRI exposure timing did not lower the heterogeneity.

### 3.11. Sensory processing

The meta-analysis for sensory processing comprised 12 studies with 17 comparisons. The most used behavioral test in this category was prepulse inhibition (13 comparisons), followed by auditory temporal rate discrimination (two comparisons), and gap crossing and olfactory investigation (one comparison each). In total, 317 SSRI-exposed animals and 310 vehicle-treated animals were included in this analysis.

Overall pooled analysis showed significantly less efficient sensory processing in animals that were developmentally exposed to SSRIs than in those exposed to vehicle (Figure 2H; Supplementary Figure 9A, SMD −0.37 [-0.69, −0.06], p<0.05). Whereas subgroup analyses did not show significantly different effects of developmental SSRI exposure depending on sex (Figure 2H; Supplementary Figure 9B, Chi² = 1.71, p=0.42) and stress exposure (Figure 2H; Supplementary Figure 9C, Chi² = 0.23, p=0.63), the effect was different depending on period of SSRI exposure (Figure 2H; Supplementary Figure 9D, Chi² = 11.67, p<0.01). More specifically, while SSRI-exposed offspring did not differ in sensory processing in those exposed prenatally (SMD 0.29 [-0.49, 1.07], p=0.47) and pre- and postnatally (SMD −0.04 [-0.31, 0.23], p=0.77), animals exposed to SSRIs postnatally showed significantly less efficient sensory processing than those exposed to vehicle (SMD −1.04 [−1.59, −0.48], p<0.001) (Figure 2H; Supplementary Figure 9D).

The heterogeneity (I^2^) of the overall analysis was 68%. The subgroups based on sex and stress exposure did not lower the heterogeneity. Subgroup analyses based on SSRI exposure timing lowered the heterogeneity to 40% for those exposed prenatally, 21% for those exposed pre- and postnatally, and 68% for those exposed postnatally.

### 3.12. Reflex and pain sensitivity

The meta-analysis for reflex and pain sensitivity comprised 11 studies with 16 comparisons. The most used behavioral tests in this category were the hot plate test and negative geotaxis (six comparisons each), followed by mechanical sensitivity and righting reflex (two comparisons each). In total, 188 SSRI-exposed animals and 200 vehicle-treated animals were included in this analysis.

Overall pooled analysis did not show significantly different reflex and pain sensitivity in animals that were developmentally exposed to SSRIs than in those exposed to vehicle (Figure 2I; Supplementary Figure 10A, SMD −0.25 [-0.73, 0.23], p=0.31). Subgroup analyses did not show significantly different effects of developmental SSRI exposure depending on sex (Figure 2I; Supplementary Figure 10B, Chi² = 1.33, p=0.51), stress exposure (Figure 2I; Supplementary Figure 10C, Chi² = 0.02, p=0.88), or period of SSRI exposure (Figure 2I; Supplementary Figure 10D, Chi² = 3.54, p=0.17).

The heterogeneity (I^2^) of the overall analysis was 77%. The subgroups based on sex, stress exposure and SSRI exposure timing did not lower the heterogeneity.

### 3.13.

Publication bias Publication bias was assessed using funnel plots. Inspection of the funnel plots supplemented with trim and fill analysis revealed no asymmetry for activity and exploration (Supplementary Figure 11A), stress coping (Supplementary Figure 11C), social behavior (Supplementary Figure 11D), motoric behavior (Supplementary Figure 11G), sensory processing (Supplementary Figure 11H), and reflex and pain sensitivity (Supplementary Figure 11I).

Using trim and fill analysis, we found an indication for funnel plot asymmetry for three behavioral categories. First, for anxiety, studies with moderate and low precision showing increased anxiety as a result of perinatal SSRI exposure were underrepresented, resulting in 20 extra data points and an adjusted estimated effect size SMD 0.26 [0.14, 0.37] (Supplementary Figure 11B). Second, for learning and memory behavior, studies showing worse test scores as a result of perinatal SSRI exposure were underrepresented, resulting in 10 extra data points and an adjusted estimated effect size of SMD −0.21 [-0.40, −0.02] (Supplementary Figure 11E). Finally, for ingestive and reward behavior, studies showing lower scores of ingestive and reward behavior as a result of perinatal SSRI exposure were underrepresented, resulting in eight extra data points and an adjusted estimated effect size of SMD −0.12 [-0.49, 0.25] (Supplementary Figure 11F).

For anxiety and learning and memory, the trim and fill analysis suggested publication bias might be at play and that the effect size we found might have underestimated the true effect. However, publication bias is only one possible explanation for funnel plot asymmetry^144^. Considering strong indications that period of drug exposure mediates the relationship between perinatal SSRI exposure and later-life behavioral outcomes, we further examined this alternative explanation. Separate funnel plots and subsequent trim and fill analysis per exposure period produced no extra data points for anxiety (Supplementary Figure 11B) and few extra data points for learning and memory (Supplementary Figure 11E). This suggests that the funnel plot asymmetry for these categories can largely be explained by subgroup heterogeneity.

### 3.14. Sleep & circadian activity

Seven studies examined the effects of perinatal SSRI exposure on outcome measures related to sleep and circadian activity (Table 4).

**Table 4:** study outcomes for sleep & circadian activity

| Study ID | Measure | Summary of outcome |
| --- | --- | --- |
| Hilakivi et al. 1987<sup>a44</sup> | Sleep-wake behavior measured with a movement sensitive mattress | Less active sleep and more wakefulness during neonatal SSRI treatment |
| Hilakivi et al. 1987<sup>c46</sup> | Sleep-wake behavior measured with a movement sensitive mattress | Less active sleep during neonatal SSRI treatment |
| Hilakivi et al. 1988<sup>a47</sup> | Sleep-wake behavior measured with a movement sensitive mattress | Less active sleep during neonatal SSRI treatment |
| Frank et al. 1997<sup>51</sup> | Sleep architecture using EEG and EMG | More non-REM-REM transitions*. No differences in sleep and wake amount. |
| Popa et al. 2008<sup>74</sup> | Sleep architecture using EEG and EMG | Total REM sleep duration and frequency is higher*. No differences in non-REM sleep. |
| Kiryanova et al. 2013<sup>99</sup> | Running wheel activity during LD, DD (baseline and after short light pulse), and LL (baseline and after long dark pulse) | Baseline: free-running period in DD was shorter*. Otherwise no differences.<br>Light pulse: larger phase advance by light pulse at CT22*, but not at CT16.<br>No difference in phase advance after dark pulse. |
| Kiryanova et al. 2017<sup>a138</sup> | Running wheel activity during LD, after LD advance, during DD (baseline and after short light pulse), and LL | No baseline differences. It took longer to re-entrain to the new LD cycle*. Interaction with maternal stress in the phase shift to light pulses at CT22*, but not at CT16. |

### 3.15. Behavior after challenges

Thirteen studies examined the effects of perinatal SSRI exposure on behavioral responses to pharmacological- and immune challenges in adulthood (Table 5).

**Table 5:**
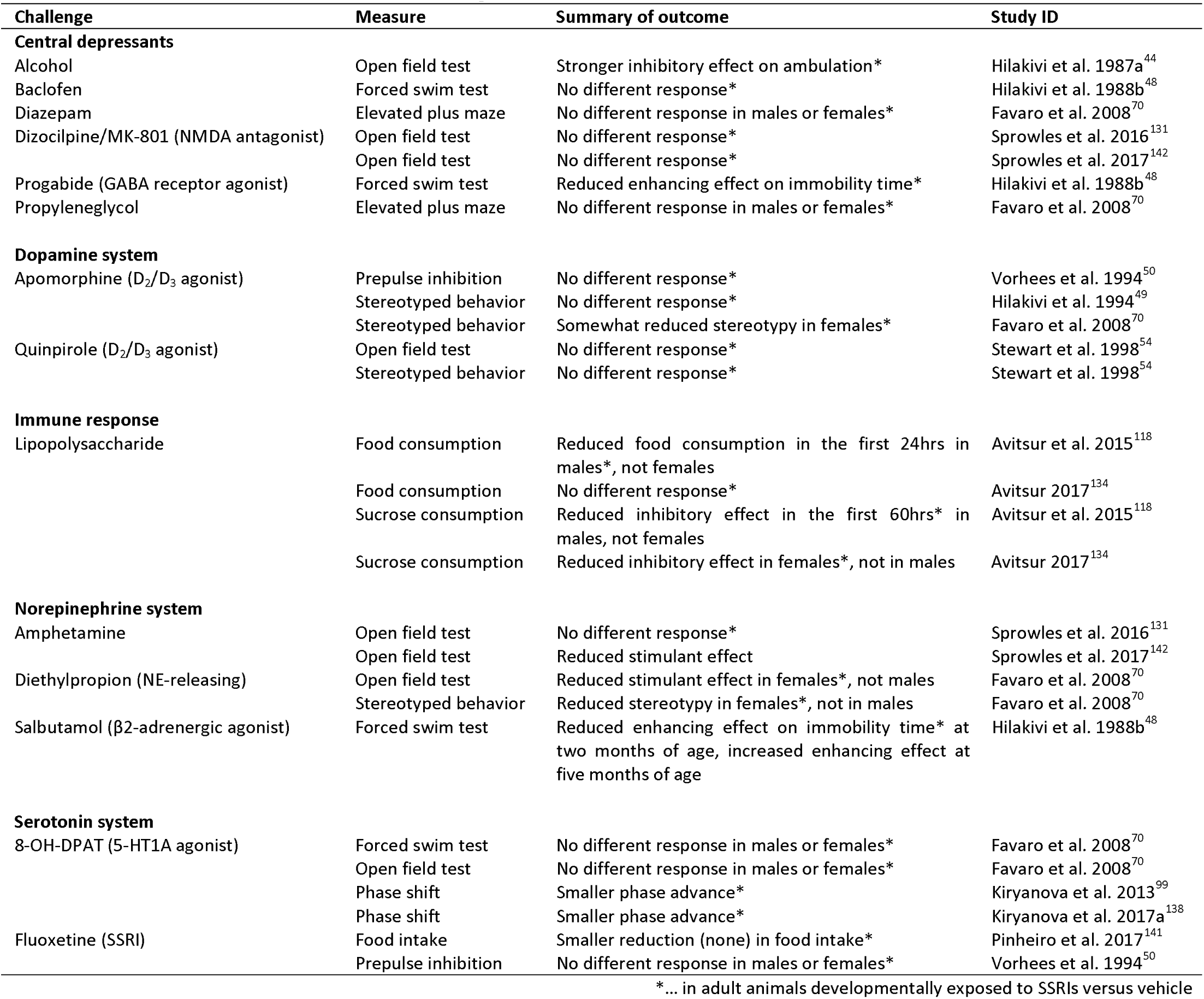
behavioral outcomes after challenges

## 4. Discussion

Our main aim was to systematically review and analyze animal studies to determine whether there is an overall effect of perinatal SSRI exposure on later-life behavior in a spectrum of behavioral domains. We included 99 publications and performed nine separate meta-analyses for different behavioral domains. We found evidence for reduced activity and exploration behavior in SSRI-exposed (N=2646) relative to vehicle-treated (N=1627) animals. In addition, we found evidence for a more passive stress coping style in SSRI-exposed (N=955) compared to vehicle-treated (N=806) animals. Lastly, we found evidence for less efficient sensory processing in SSRI-exposed (N=317) versus vehicle-treated (N=310) animals. All effect sizes were small to medium. We found a tendency for increased anxiety (p=0.06), while no differences were found in social behavior, learning and memory, ingestive- and reward behavior, motoric behavior, and reflex and pain sensitivity as a result of developmental SSRI exposure in animals.

### 4.1. Modulating role of sex, stress exposure, and timing of SSRI exposure

Our secondary aim was to examine the conditions under which a potential effect of developmental SSRI exposure on later-life behavior would manifest itself. We selected three moderators to examine using subgroup analyses: animal sex, presence of perinatal stress exposure (reflecting efforts to mimic aspects of a maternal depressed mood in animal models), and timing of SSRI exposure.

The sex of the animal tested explained part of the heterogeneity in the data for two behavioral categories. The male- and the mixed-sex subgroups showed significantly lower scores for activity and exploration in SSRI-exposed offspring relative to vehicle-exposed offspring, whereas in females there was no significant difference. Interestingly, most other behavioral categories also showed larger effect sizes in males than in females, although these were not statistically significant effects. For learning and memory, we found a significant effect of SSRI exposure in the mixed-sex subgroup, but not in the male or female subgroups. These results may be explained by confounding effects of other moderators such as the timing of SSRI exposure. In general, it is important to realize that subgroup analyses are observational in nature, as they are not based on randomized grouping. To enable more reliable and informative analyses of potential sex effects in the future, researchers should make their data available separately for males and females in a supplementary file.

We found no evidence for a modulatory role of stress exposure on the effects of developmental SSRI exposure on behavior. This could be a reflection of a true absence of an interaction between perinatal stress- and SSRI exposure. It could also be due to the large heterogeneity and wide confidence interval in the stress-exposed group, as a result of the relatively low number of comparisons and the variation in the nature, timing and intensity of the stress protocols used. A selective meta-analysis including only those studies reporting on both stress-unexposed and stress-exposed offspring would yield more insight into the effects of stress exposure, but is beyond the scope of the current review.

The specific period the animal was exposed to an SSRI (prenatal, postnatal, or both) explained the most heterogeneity in the data out of the 3 subgroup analyses we performed. Animals exposed to ^37^ SSRIs postnatally – this roughly corresponds to the third trimester in humans – showed reductions in activity and exploration, social behavior, learning and memory, and sensory processing scores, while ^37^ animals exposed prenatally – roughly corresponding to the first two trimesters in humans – did not.

### 4.2. Potential mechanisms

The effects of developmental SSRI exposure on later-life behavioral outcomes are the result of a combination of direct effects on the developing brain and indirect effects, for example through changes in placental and maternal homeostasis ^14^ and postnatal maternal care ^145^. The serotonin system consists of 15 different receptors that are key players at crucial neurodevelopmental stages, regulating neurogenesis, apoptosis, axon branching and dendritogenesis^11^. Many of the studies included in the synthesis of evidence in the current review, which have been selected on the presence of behavioral outcomes, also include outcomes reflecting brain health from the global to the molecular level: the corticosterone response to stress ^74, 81, 96, 100, 123, 124, 126, 130, 135^, brain structure and connectivity ^71, 77, 84, 93, 101, 110, 122^, neuronal health ^59, 82, 85, 89, 104, 109, 111, 126, 135^, monoamine concentrations in the brain ^43, 44, 46, 59, 105, 116, 117, 133, 135, 140, 141^, protein expression in the brain – mainly related to the serotonergic system and neurogenesis ^62, 71, 78, 88, 91, 97, 127, 129, 141^, gene expression ^76, 94, 96, 112, 113, 120, 121, 123, 137, 141, 143^, and epigenetic modifications ^76, 112, 114, 124^.

Several mechanisms may underlie our current findings. Earlier work in serotonin transporter (SERT) knockout rodents, which lack the SERT and thereby mimic SSRI exposure from conception onwards, showed that 2 main neural networks were changed compared to wildtype rodents: the somatosensory cortex and the corticolimbic circuit^15^. The first network is likely related to the sensory processing deficits we found in SSRI-exposed animals. Axons extending from the thalamus to the cortex transiently express SERT during development, and disruption of serotonin availability cause them to form aberrant trajectories ^146, 147^ and affect the development of the somatosensory cortex^77, 148^. The second network could be responsible for the effects seen on activity and exploration and stress coping behaviors. In addition, changes in neuroendocrine function could play a role in the development of a ^32^ more passive stress coping style in SSRI-exposed animals. It is unclear whether the effects of early SSRI exposure on activity and exploration behavior and stress coping behavior have overlapping brain correlates.

Lastly, we found higher effect sizes in males (relative to females). In general, male offspring seem more vulnerable to various types of stressors during pregnancy than female offspring^149^. Early SSRI-exposure may affect males and females differently because of the sex-specific maturation of the serotonin system^14^. For instance, serotonin levels in early postnatal life in rodents are different in males and females: male pups show a peak of serotonin at PND3, while female pups show more stable serotonin levels with a later peak^150^.

### 4.3. Clinical implications

The neurodevelopmental pattern of the serotonin system is remarkably conserved across species ^29, 32, 33^. Therefore, rodent studies of early SSRI exposure can yield important insights and circumvent some of the difficulties of studying this phenomenon in humans. Preclinical and clinical studies on this topic should ideally continuously inform and supplement each other.

The finding that early SSRI exposure is linked to a passive coping style in adult animals is an interesting manifestation of the “SSRI paradox”. Treatment with antidepressants in adulthood generates a more active coping style in animals^151^ and alleviates symptoms of depression in humans. Conversely, SSRI treatment in the perinatal period leads to a more passive coping style in animals later in life. The most common behavioral test in this category is the forced swim test ^152^. The basic premise of this test is that, confronted with an inescapable situation in a cylinder of water, rodents can either actively try to escape, or go into a state of passive floating. This passive behavioral response may be analogous to maladaptive responses to stress as seen in humans with neuropsychiatric disorders^153^. Similarly, disruptions in sensory processing like those associated with early SSRI exposure in animals are present in ^154^ a spectrum of neuropsychiatric diseases in humans. Our results suggest that the increased risk of symptoms of neuropsychiatric disorders for those prenatally exposed to SSRIs, as indicated in some ^25^ studies, might be mediated by differences in stress coping, sensory processing and perhaps anxiety^25, 155^.

A major challenge in human studies is to properly control for the confounding factor of maternal ^33^ psychiatric condition. Statistical methods aim to approximate this, illustrated by the finding that the association between in utero SSRI exposure and risk of ASD was not significant when controlled for maternal psychiatric diagnosis^156^. However, a clean comparison between children from SSRI- and vehicle-treated mothers without any psychiatric history is not available. Our results suggest that perinatal SSRI exposure exerts effects on neurodevelopmental outcomes at least partially independently from maternal psychiatric condition. As maternal psychiatric disorder might interact with SSRI use to influence offspring outcomes^14, 32^, researchers and clinicians have questioned how clinically relevant rodent studies are. To address this, animal models have been developed aiming to study SSRI exposure in light of maternal (pre)gestational stress^14^. Our current results do not support the notion of an interaction effect of maternal stress exposure and perinatal SSRI exposure on behavioral outcomes in offspring, although the number of studies that examine this is still limited.

The first few postnatal weeks in rodents are instrumental in the maturation of both the serotonin system and cortical circuit wiring, and also show the highest levels of serotonin and its metabolites in the brain^29^. In terms of brain development, this period is approximately equivalent to the third trimester of human gestation^37^. Our finding that SSRI exposure in the first postnatal weeks has the largest effect on later-life behavior in animals therefore implies that SSRI treatment during the last months of pregnancy should have the largest effect on human outcomes. Clinical studies investigating the effect of timing of SSRI exposure are limited and inconsistent. In line with current results, a recent study found that late-pregnancy SSRI exposure was associated with greater depressed and anxious symptoms in children^20^, whereas a meta-analysis found that exposure to SSRIs during the first trimester was most consistently associated with later diagnosis of mental disorders^25^. Perhaps for good reasons, many women discontinue SSRI use over the course of pregnancy, with the least users in the third ^5^ trimester, making this the most challenging trimester to study. Our results suggest, however, that the timing of SSRI exposure should be a key variable of interest in future human studies.

### 4.4. Limitations and strengths

One of the limitations of this study is that the quality of the pooled analyses is only as high as the quality of the individual studies that it consists of, which is hard to determine. Basic characteristics of best practices in experimental studies, such as blinding and randomization, were sparsely reported. This is often the case with animal studies^157, 158^. Especially problematic is the high percentage of studies not reporting all outcome measures that were described in their respective methods section, potentially introducing bias. However, inspection and analysis of funnel plots in search of indications for publication bias was mostly reassuring. Funnel plot asymmetry was largely accounted for by subgroup heterogeneity and therefore likely not a sign of publication bias. Other limitations stem from the features of the animal studies we included, which might not make them optimally suitable for translation to the human situation. For instance, many studies employed bolus daily injections that might lead to transient high serum concentrations of the compounds and their metabolites because of their relatively short half-life in rodents^32^. In humans, SSRI use leads to more stable concentrations over the course of the day. In addition, dosing and route of administration varied widely^33^.

Additional limitations of this study originate from the choices that we had to make during data analysis. Many behavioral tests in the studies that we included have a complex temporal design where, for instance, reflex development or learning is assessed over several days or sexual behavior over several weeks. For lack of an overall score of performance in these tests, we opted to include one time-point in our analyses, thereby reducing these often elegant study designs to a snap shot. Comparison between studies is further complicated by the fact that not all studies report on similar time-points. In addition, besides the subgroup analyses we performed, there are other mediators that may be of interest. These analyses were outside the scope of the current review, but we do think that comparisons between the different SSRIs, the different dosages, animal species, timing of behavioral testing, and the specific test used within each category would be interesting for future studies and meta-analyses. For example, preliminary data exploration along these lines suggests that it is mainly the elevated plus maze that does not show a net effect of perinatal SSRI exposure within the category anxiety. It would be interesting to explore this further.

The strength of this review is that it is the first effort to comprehensively summarize and quantitatively analyze all available evidence on developmental SSRI exposure on behavioral outcomes in animals. The sheer number of animals included in our analyses – hundreds to thousands depending on behavioral category – gives us statistical power that far exceeds the standard in animal studies. Considering the increasing use of SSRIs during pregnancy^1–4^ and the uncertainties about their long-term effects on the developing neurobiology of the child^159^, studies of this phenomenon are necessary. We think this review could be valuable to the field, as we were able to concisely summarize the available animal evidence in order to inform design of future preclinical-as well as clinical studies.

### 4.5. Recommendations and future perspectives

Animal studies will continue to play an important role in this field because of their experimental nature and the ability to mechanistically study the developmental effects of SSRIs. To improve their transparency, quality, and utility, pre-registration of animal experiments (e.g., www.preclinicaltrials.eu) should become common practice.^160^ In addition, reporting of animal studies should be improved by adherence to guidelines such as the ARRIVE guidelines^161, 162^. Animal studies should be expected to adhere to a high standard of reporting for various reasons: substantial public funds are used to support this work, animals are sacrificed, and the research informs clinical study design, decision making, and policy. We would like to emphasize that, although those responsible for making (all) research results available to the scientific and wider community are the researchers themselves, other people and organizations such as funding agencies, universities, collaborating companies, journal editors and peer reviewers should all use their influence to make this the norm.

As to future animal study design, we encourage recent trends and requirements to study both males and females^163^. Females are understudied, and considering that we found indications of sex effects, it is clearly of interest to study both sexes. Additionally, the potential interactions of SSRI use with features of maternal depression remain underinvestigated in animals but are of high translational value. Further mechanistic studies are required to elucidate the neurobiological underpinnings of behavioral symptoms affected by early SSRI exposure. In particular, it remains to be understood whether the effects found on activity and exploration behavior can be traced back to the same neurodevelopmental processes as those found on stress coping behavior. Shifting perspectives slightly, one might wonder why early SSRI exposure does not seem to lead to stronger and more aberrant behavioral alterations than it does, considering the ubiquitous role of serotonin in the brain. Animal studies shed light on individual differences in susceptibility and resilience to the effects of early SSRI exposure, for example using strains of rats differing in their novelty seeking traits^164^.

Implications for future clinical study design appear noteworthy as well: there is a clear need for studies on the effects of early SSRI exposure on mental health and behavior extending into adulthood^159^, especially considering that phenotypic differences may emerge only after adolescence^30^. In addition, while examining the risk for developing mental disorders is important, it could be equally or perhaps more informative to focus on their shared symptoms. Changes in activity and exploration, stress coping, and sensory processing are relevant to people’s quality of life, even if they are not necessarily tied to the diagnosis of a mental disorder. Although subgroup analyses are observational by nature, our results suggest a strong effect of the timing of exposure to SSRIs on their long-term effect, with exposure in the period corresponding to the third trimester in humans conferring the biggest effects. Future studies in human populations should therefore seek to include timing of exposure as a key variable of interest, since this knowledge, if confirmed in humans, bears great interest for clinicians and pregnant women suffering from depression.

## Contributors

JH and JO conceived the study. AR and JR performed the systematic search. AR, LW and JR performed the screening and data extraction process. AR analyzed the data. JL advised on methodology. AR wrote the manuscript, which was revised critically by the other authors LW, JR, JL, JH and JO.

## Acknowledgments

We thank the authors who have provided us with additional information or data related to their studies. We also thank Rob de Vries from SYRCLE for consultation on data analysis, and the members of the Behavioral Neuroscience group at the University of Groningen who consulted on behavioral test categorization.

---

**Supplementary Figure 1:**
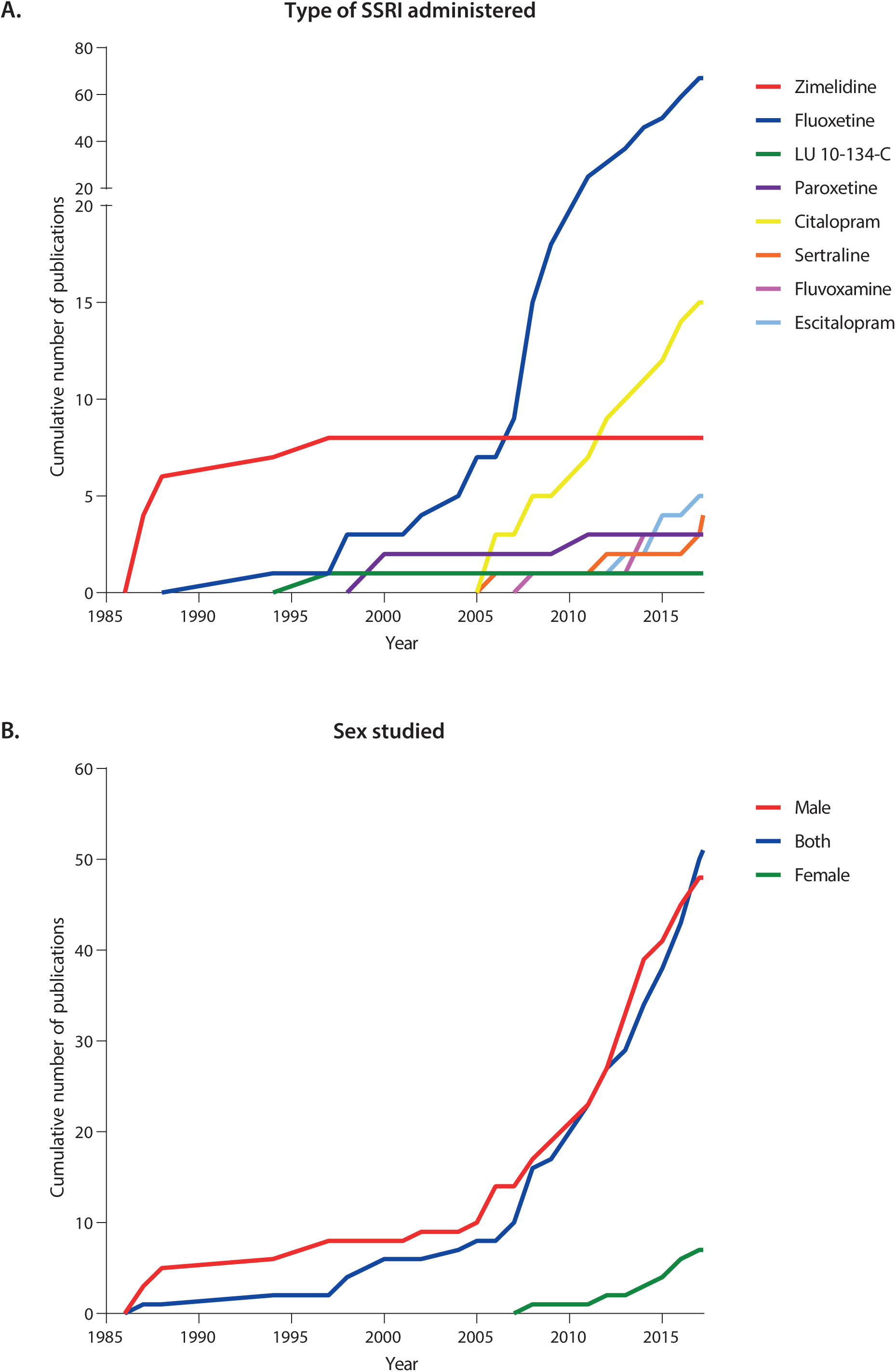
Historical perspective of study characteristics. The cumulative number of publications published each year on behavioral outcomes after perinatal SSRI exposure in animals, with a focus on (A) the type of SSRI administered and (B) the sex studied.

**Supplementary Figure 2:**
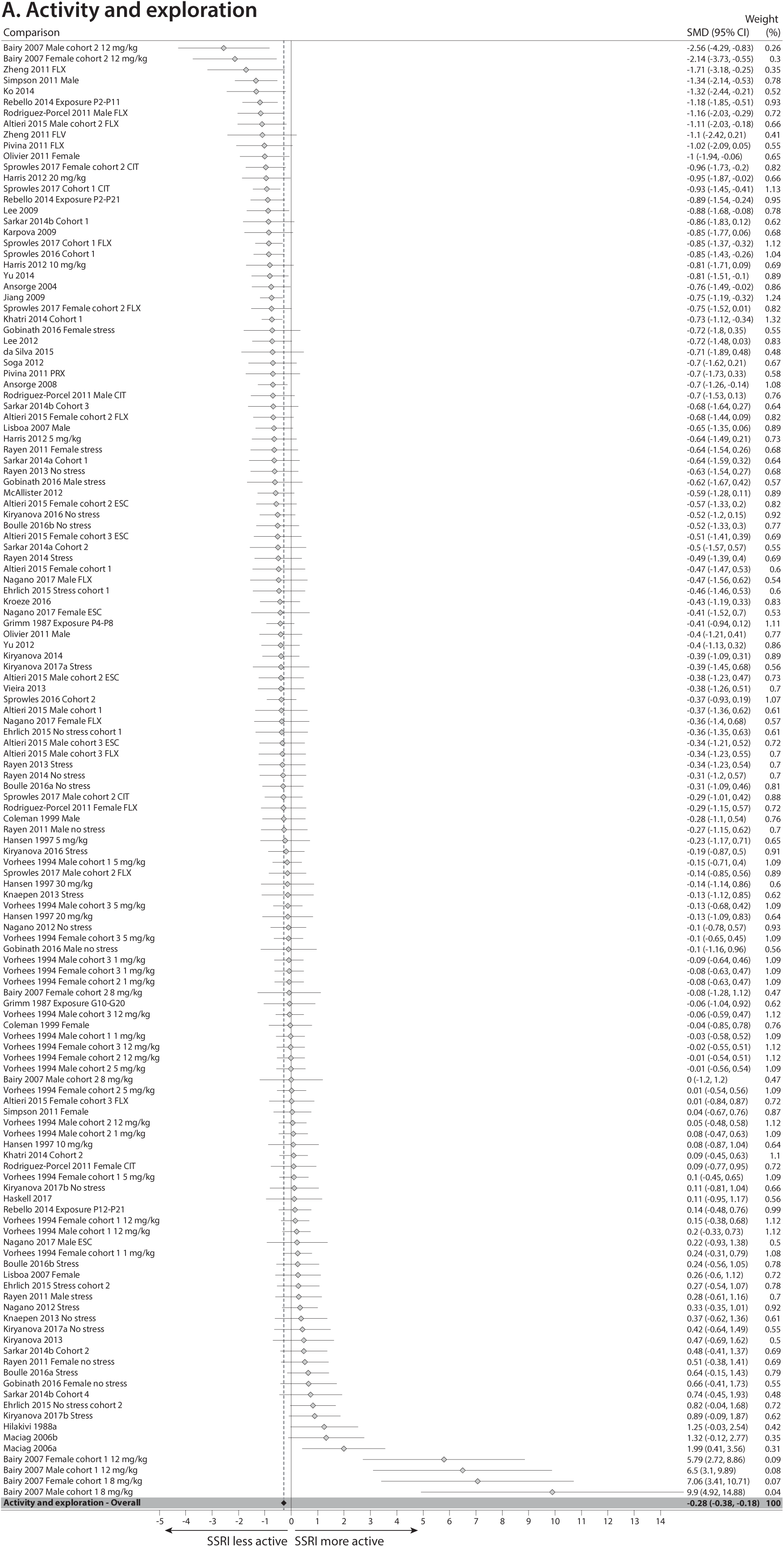
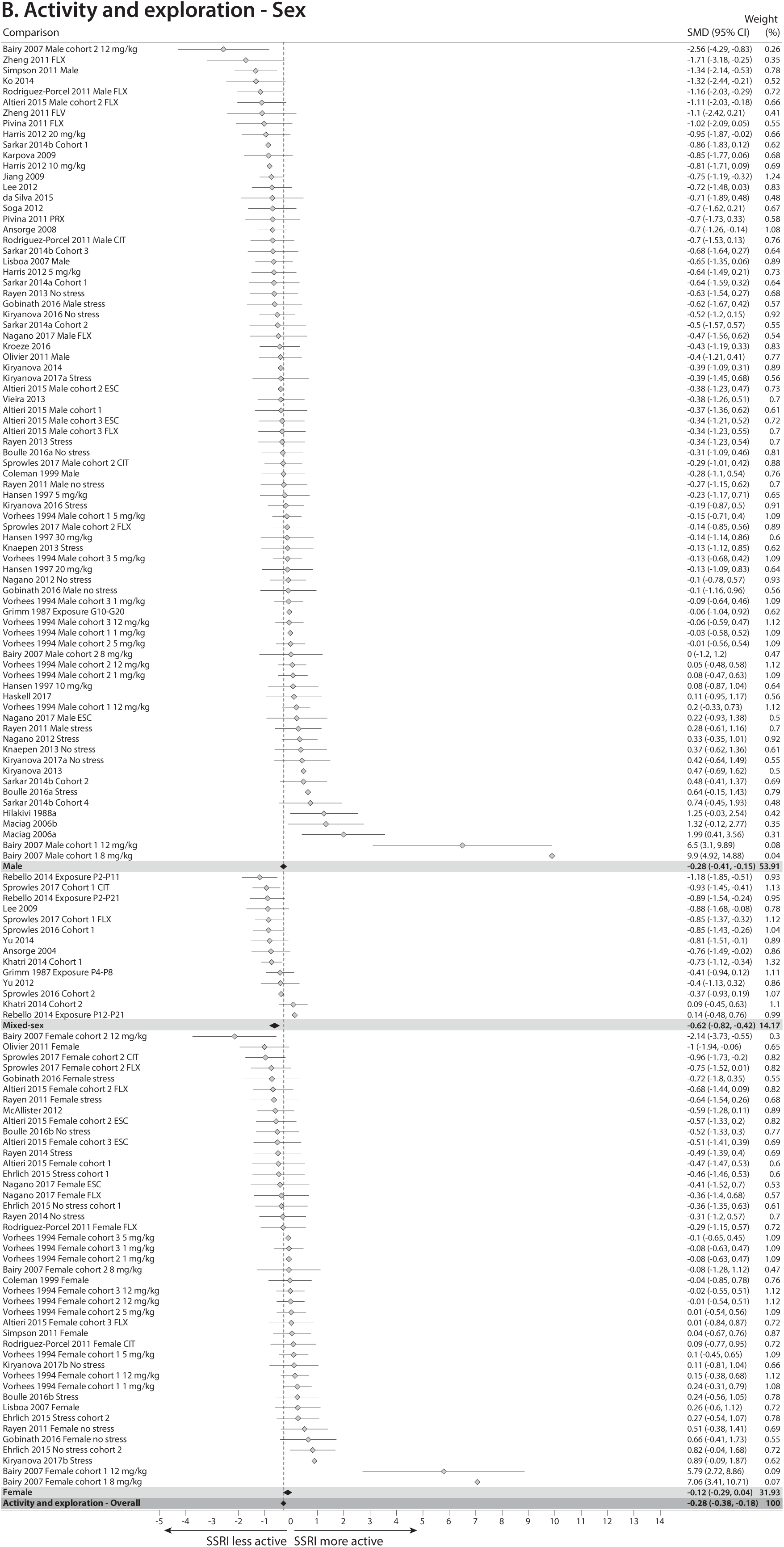
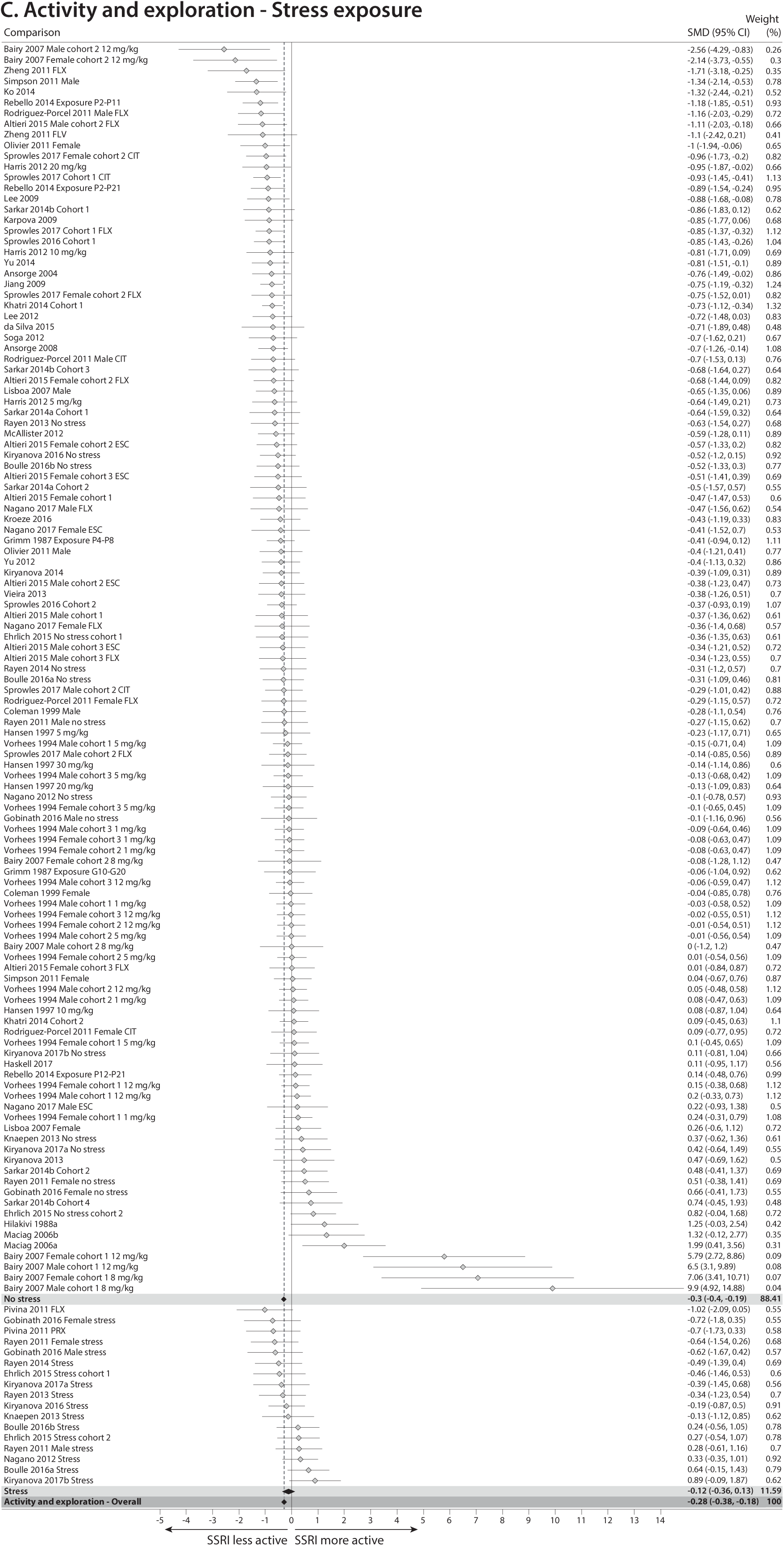
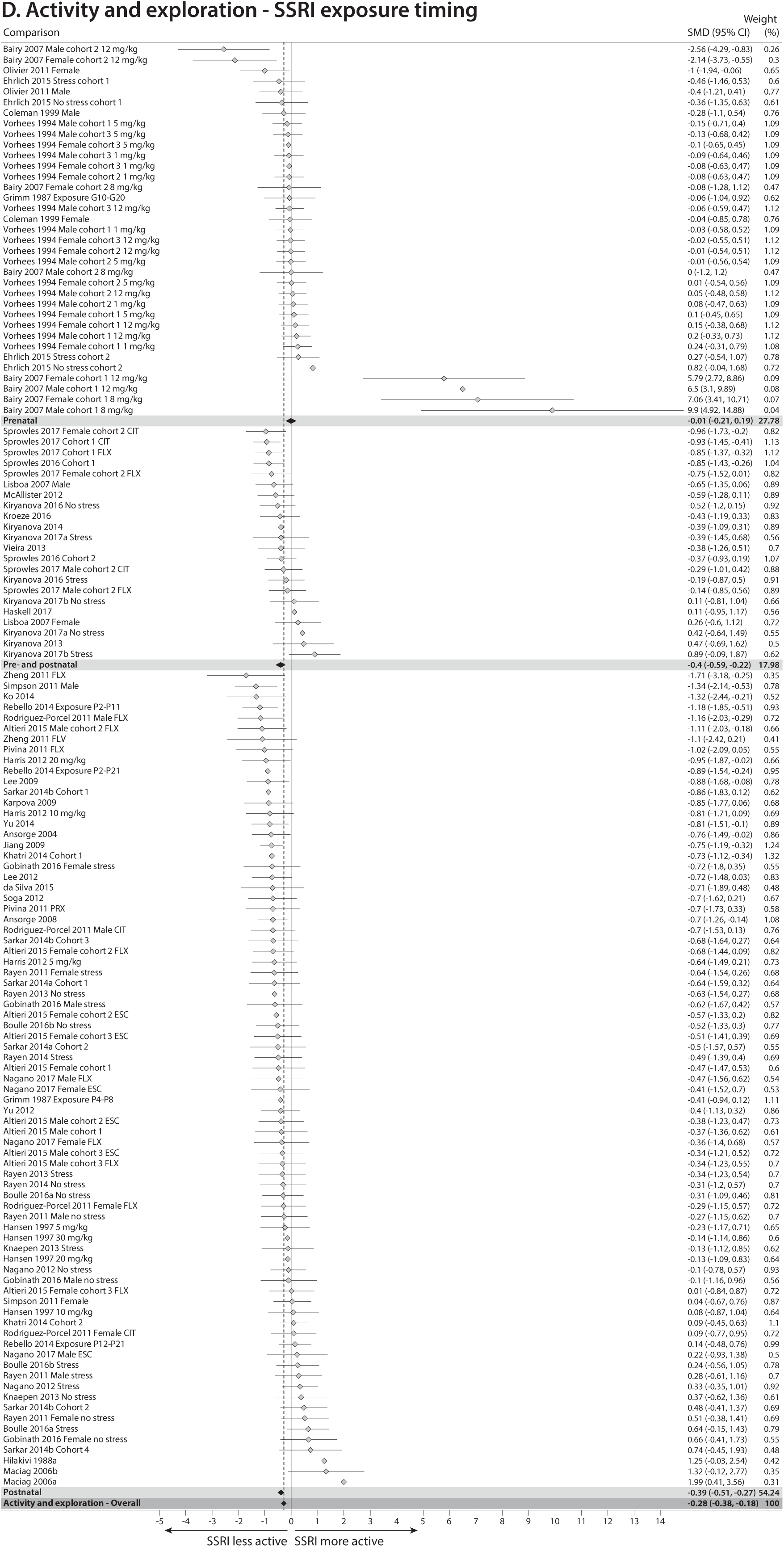
Forest plots of meta-analysis comparing animals perinatally exposed to SSRIs to those exposed to vehicle on the behavioral outcome activity and exploration. (A) Overall analysis. (B) Subgroup analysis based on sex. (C) Subgroup analysis based on presence/absence of stress exposure. (D) Subgroup analysis based on SSRI exposure timing.

**Supplementary Figure 3:**
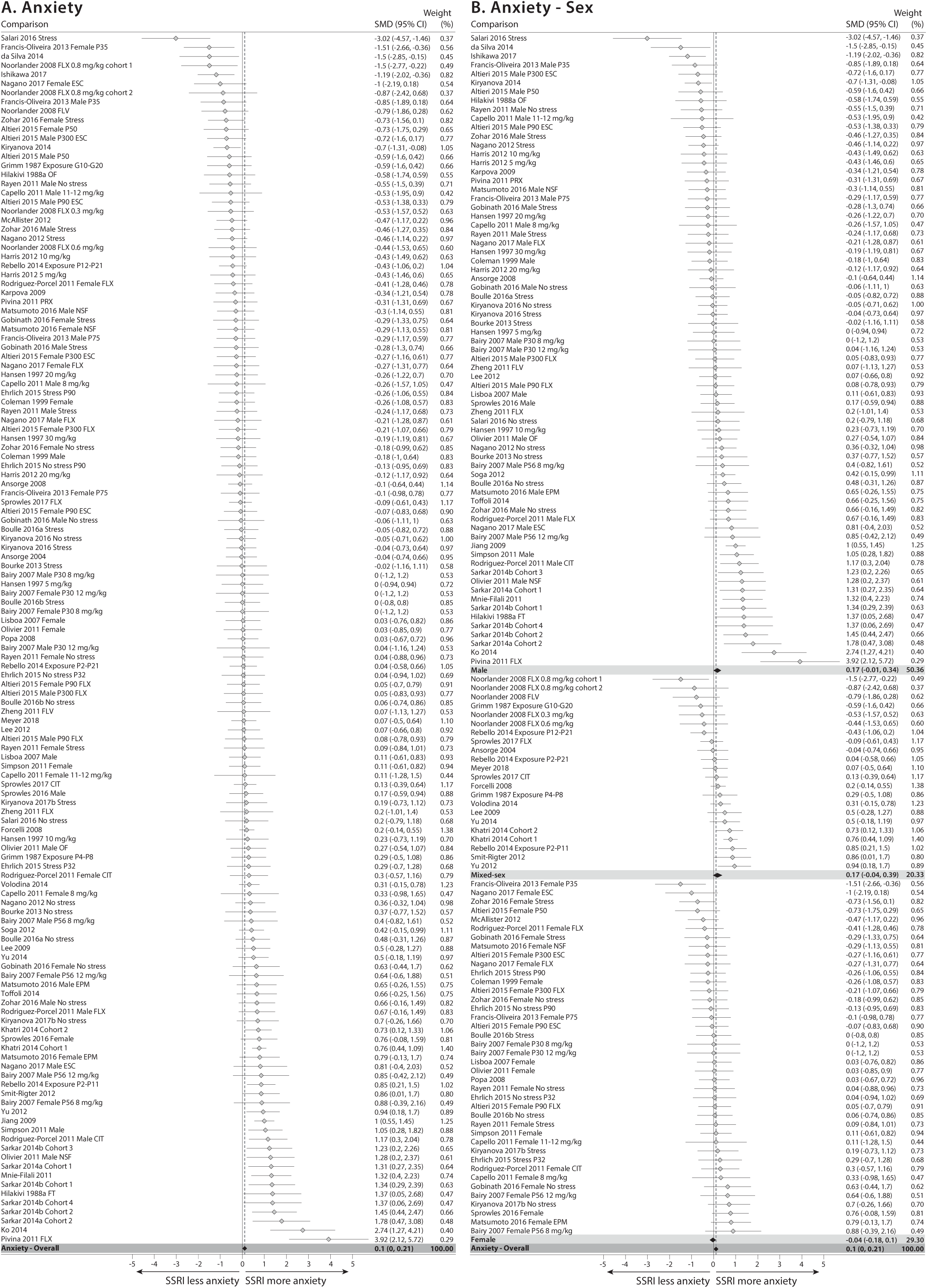
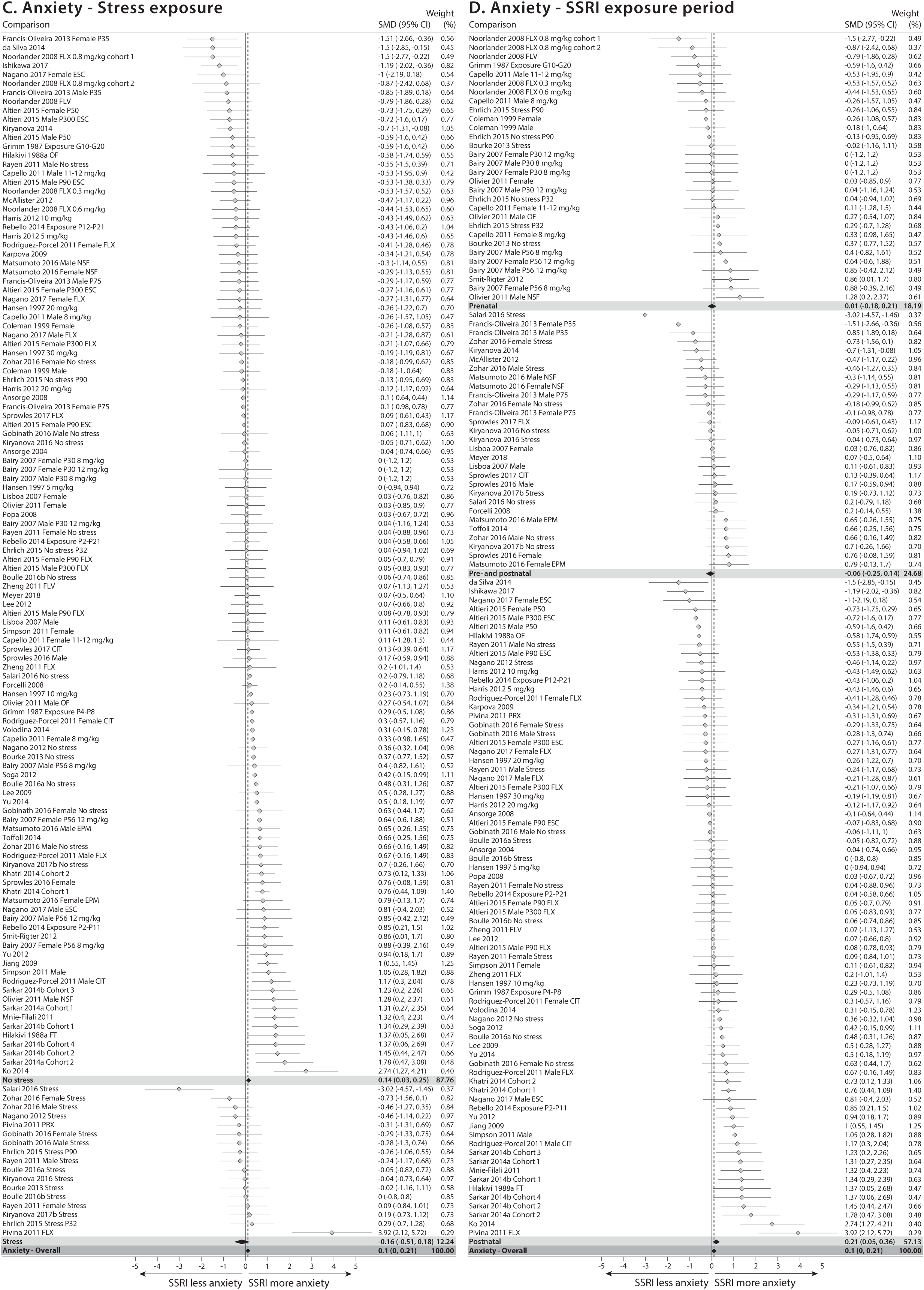
Forest plots of meta-analysis comparing animals perinatally exposed to SSRIs to those exposed to vehicle on the behavioral outcome anxiety. (A) Overall analysis. (B) Subgroup analysis based on sex. (C) Subgroup analysis based on presence/absence of stress exposure. (D) Subgroup analysis based on SSRI exposure timing.

**Supplementary Figure 4:**
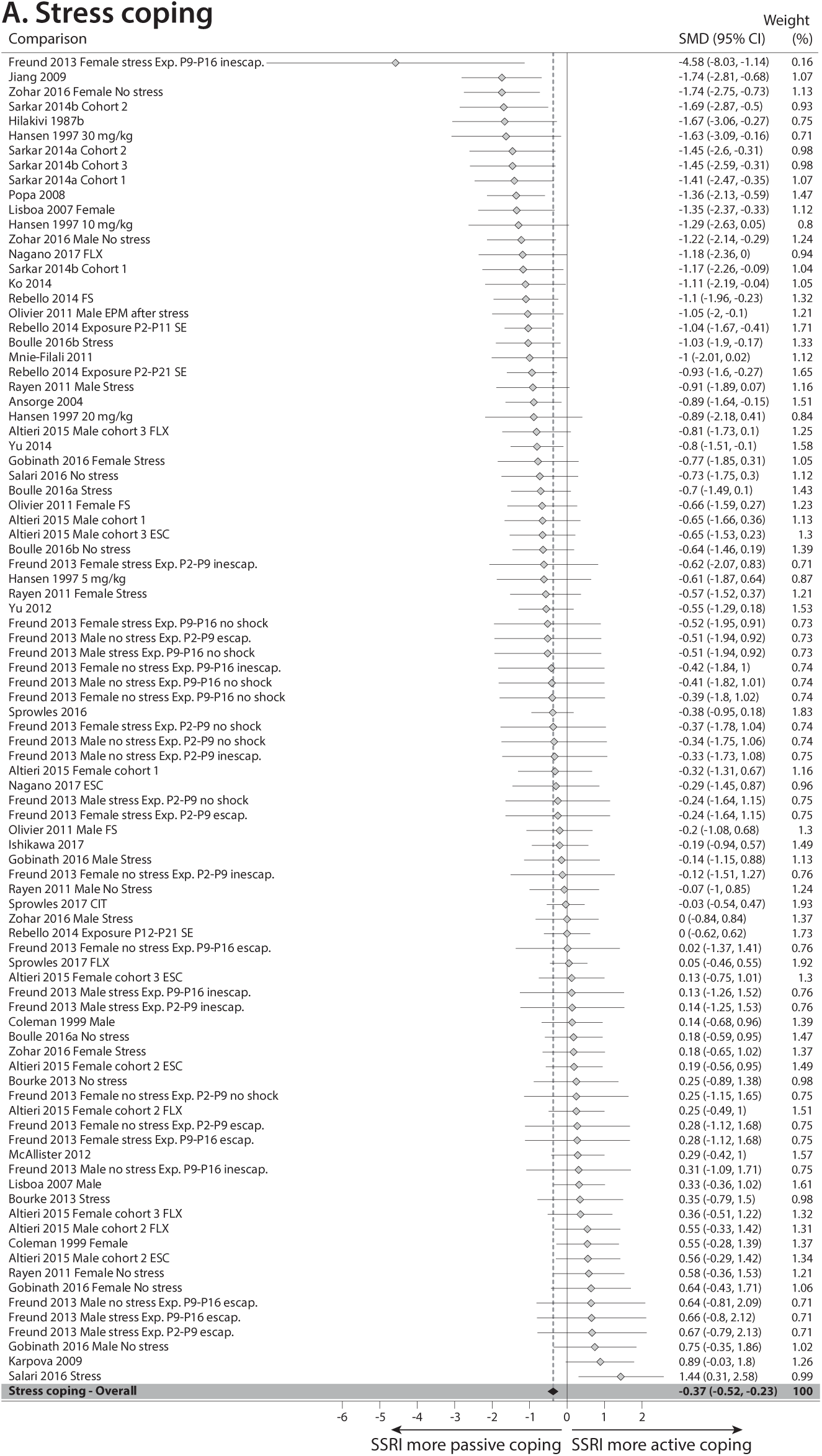
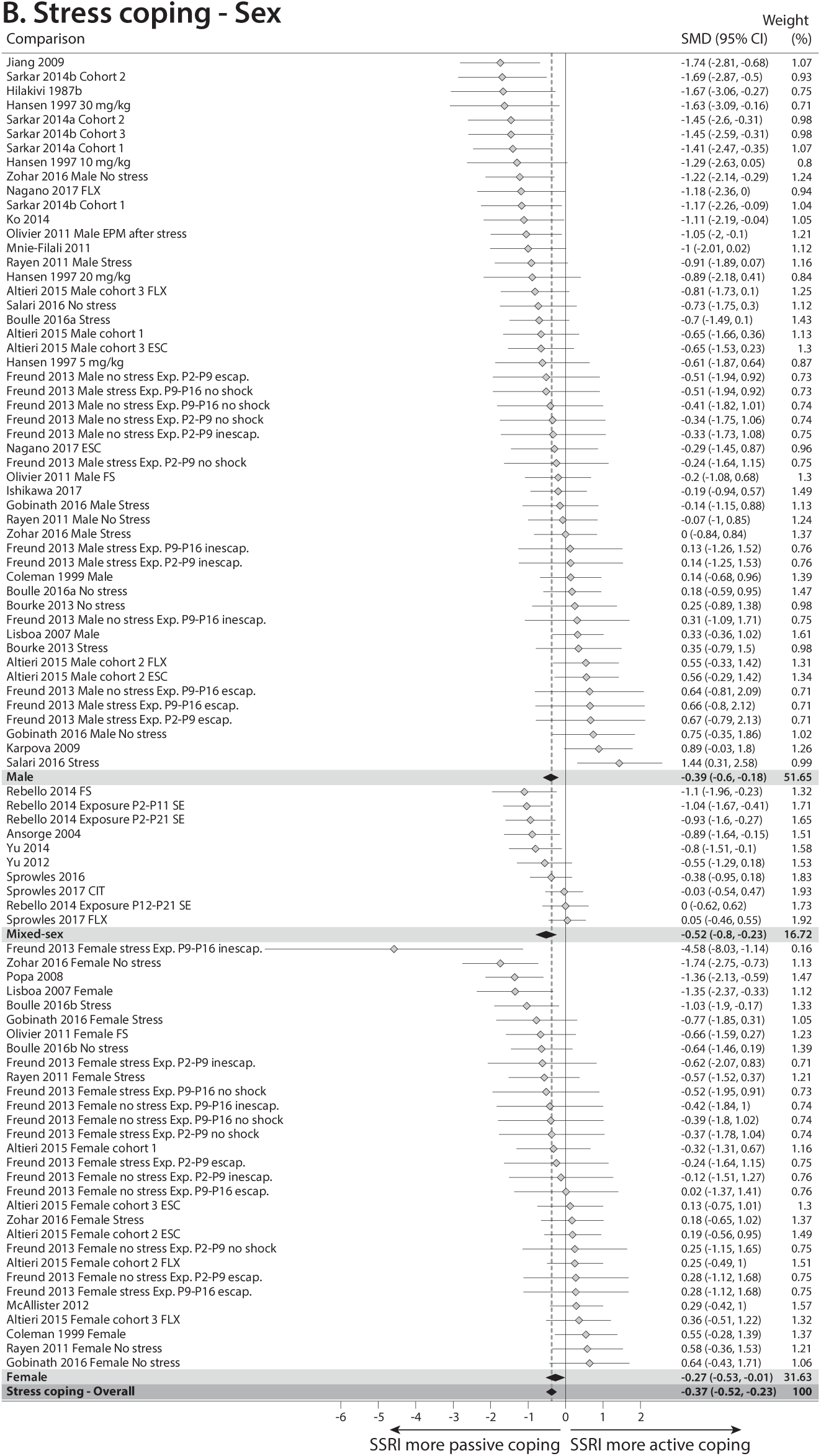
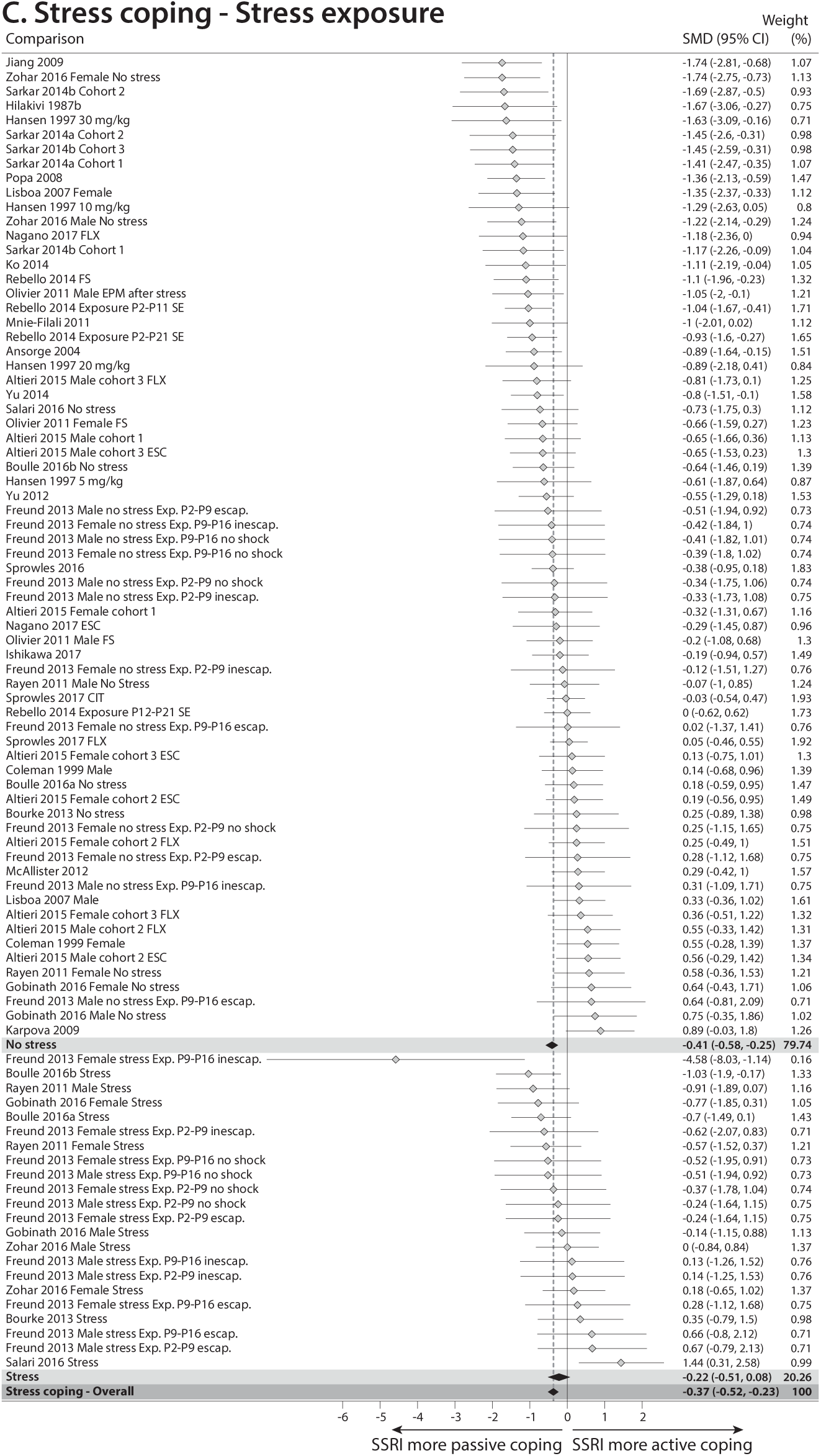
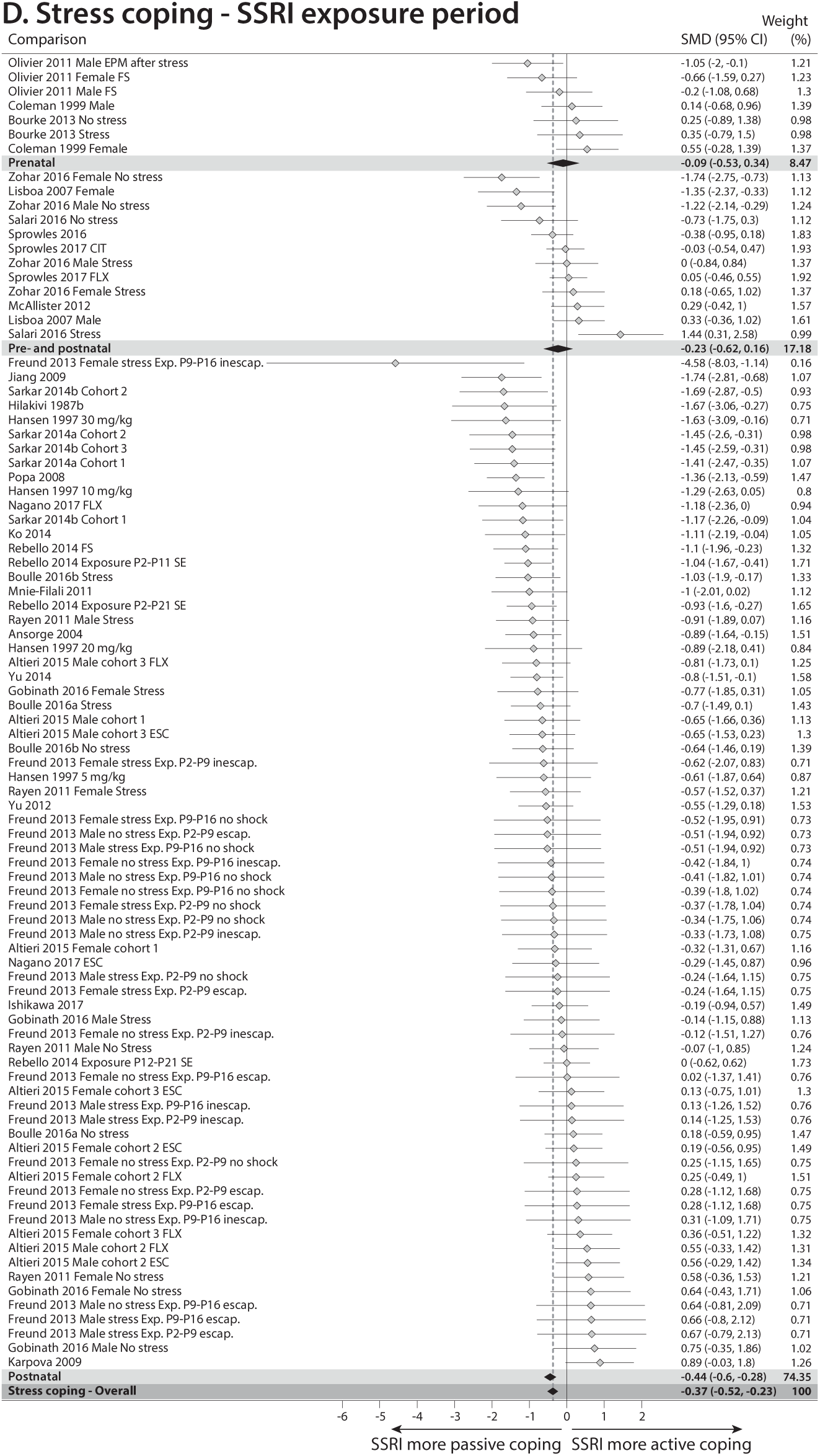
Forest plots of meta-analysis comparing animals perinatally exposed to SSRIs to those exposed to vehicle on the behavioral outcome stress coping. (A) Overall analysis. (B) Subgroup analysis based on sex. (C) Subgroup analysis based on presence/absence of stress exposure. (D) Subgroup analysis based on SSRI exposure timing.

**Supplementary Figure 5:**
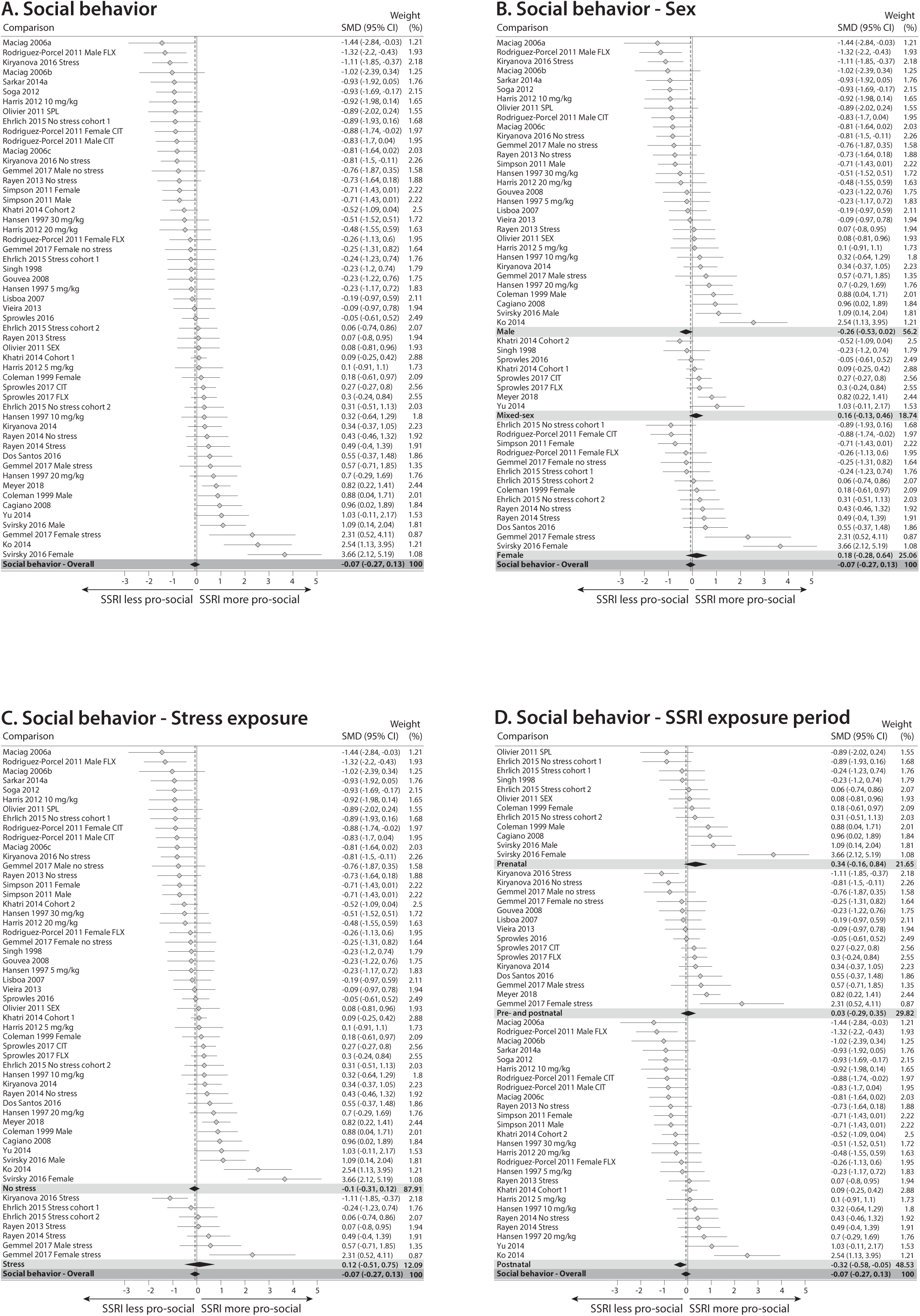
Forest plots of meta-analysis comparing animals perinatally exposed to SSRIs to those exposed to vehicle on the behavioral outcome social behavior. (A) Overall analysis. (B) Subgroup analysis based on sex. (C) Subgroup analysis based on presence/absence of stress exposure. (D) Subgroup analysis based on SSRI exposure timing.

**Supplementary Figure 6:**
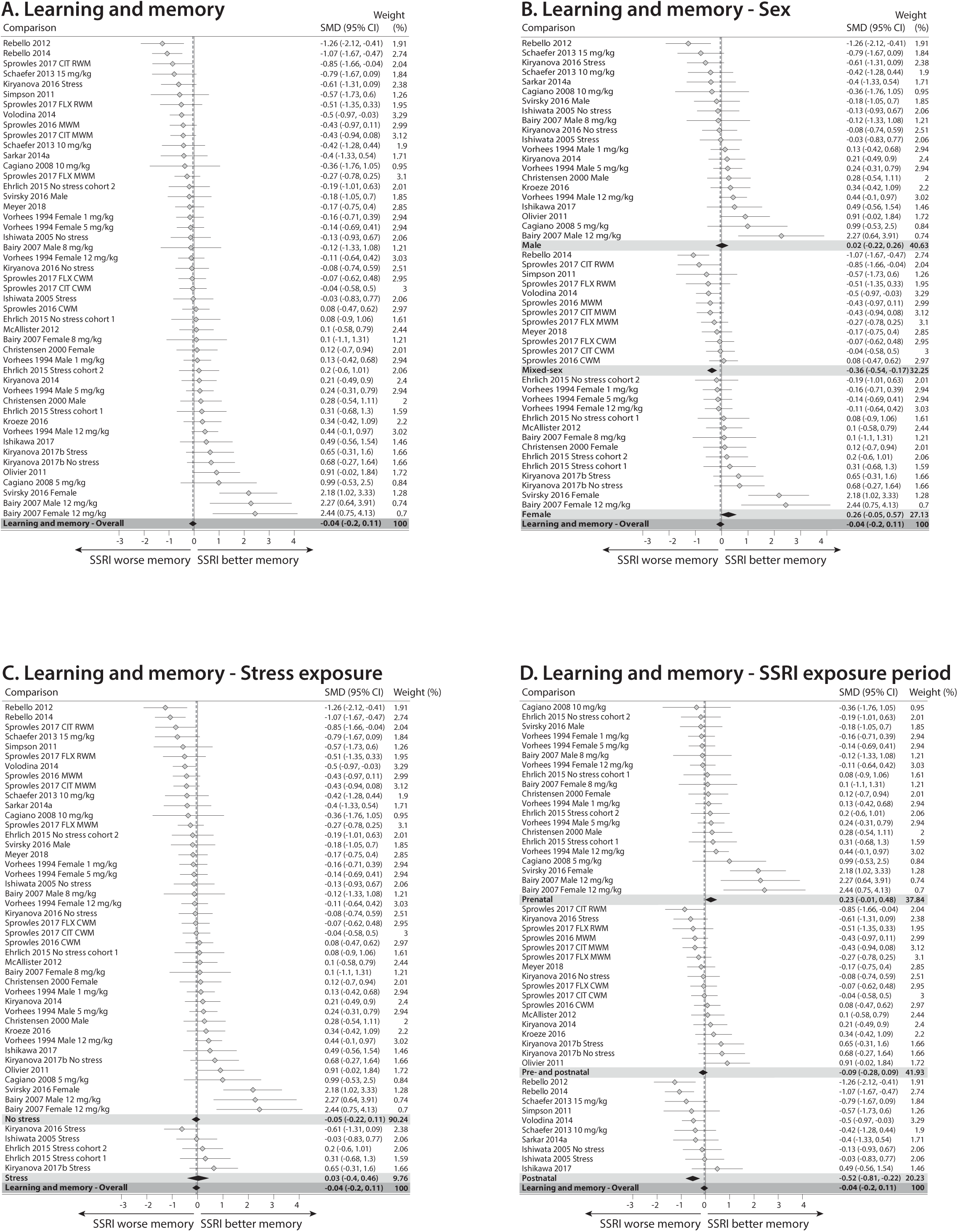
Forest plots of meta-analysis comparing animals perinatally exposed to SSRIs to those exposed to vehicle on the behavioral outcome learning and memory. (A) Overall analysis. (B) Subgroup analysis based on sex. (C) Subgroup analysis based on presence/absence of stress exposure. (D) Subgroup analysis based on SSRI exposure timing.

**Supplementary Figure 7:**
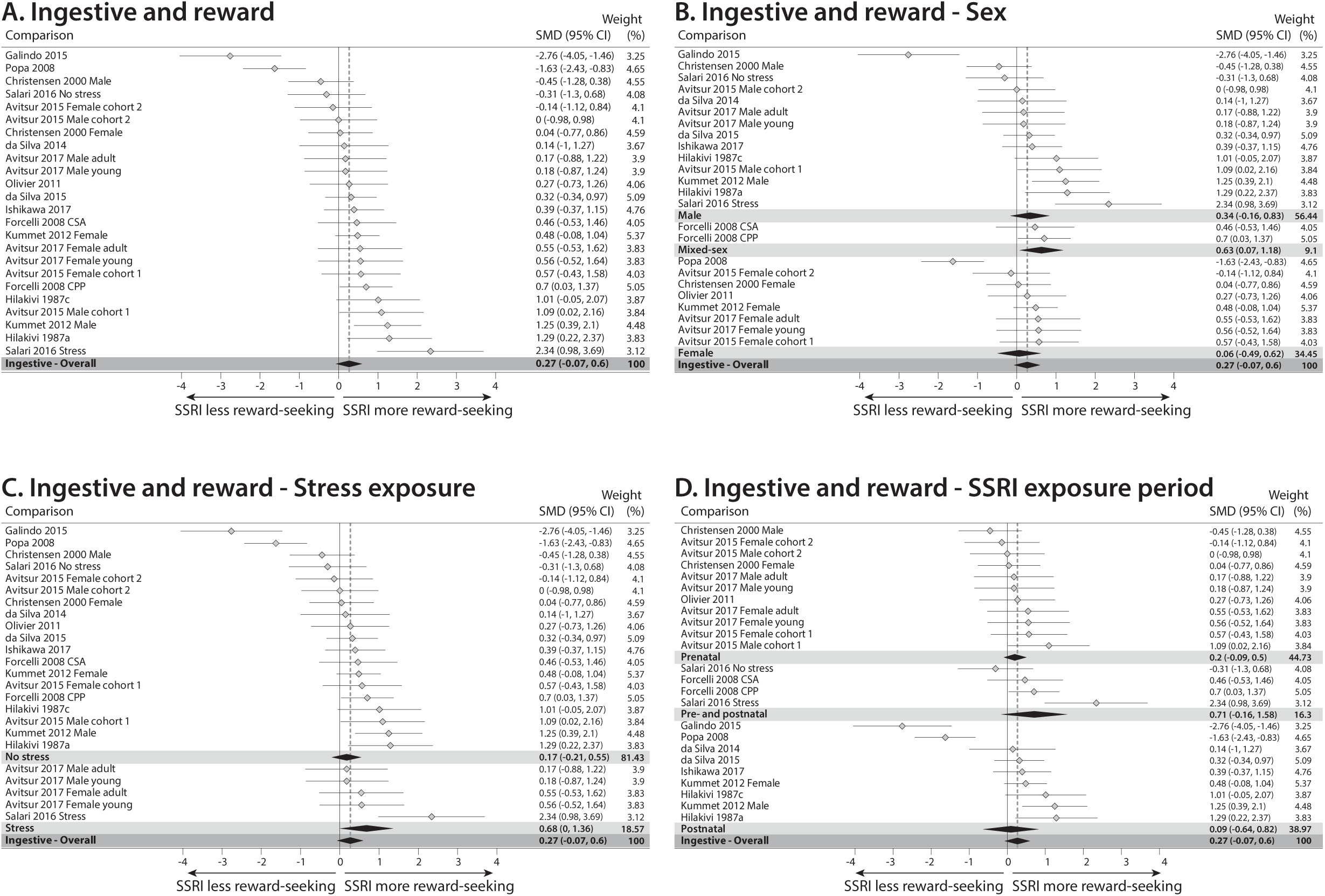
Forest plots of meta-analysis comparing animals perinatally exposed to SSRIs to those exposed to vehicle on the behavioral outcome ingestive- and reward behavior. (A) Overall analysis. (B) Subgroup analysis based on sex. (C) Subgroup analysis based on presence/absence of stress exposure. (D) Subgroup analysis based on SSRI exposure timing.

**Supplementary Figure 8:**
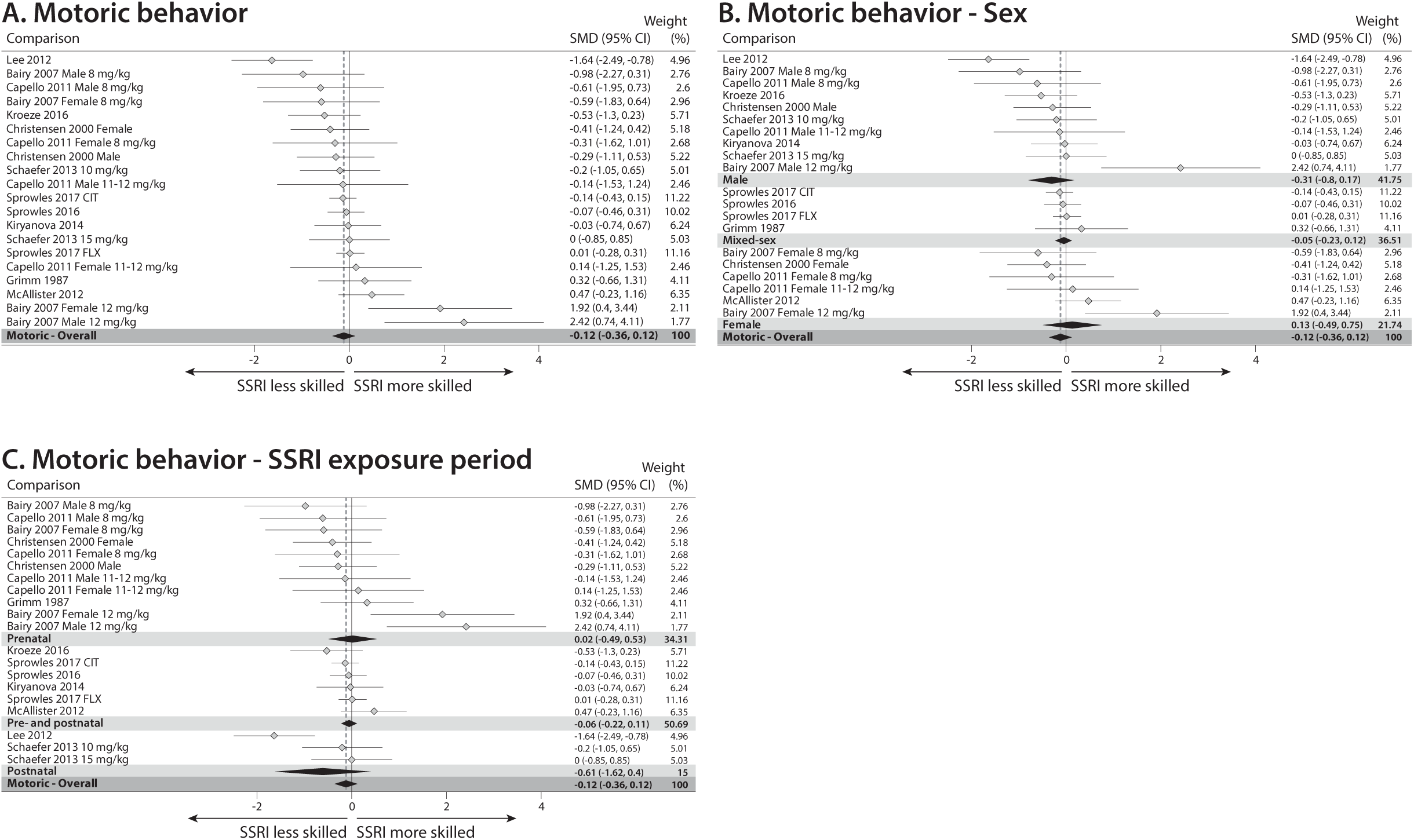
Forest plots of meta-analysis comparing animals perinatally exposed to SSRIs to those exposed to vehicle on the behavioral outcome motoric behavior. (A) Overall analysis. (B) Subgroup analysis based on sex. (C) Subgroup analysis based on SSRI exposure timing.

**Supplementary Figure 9:**
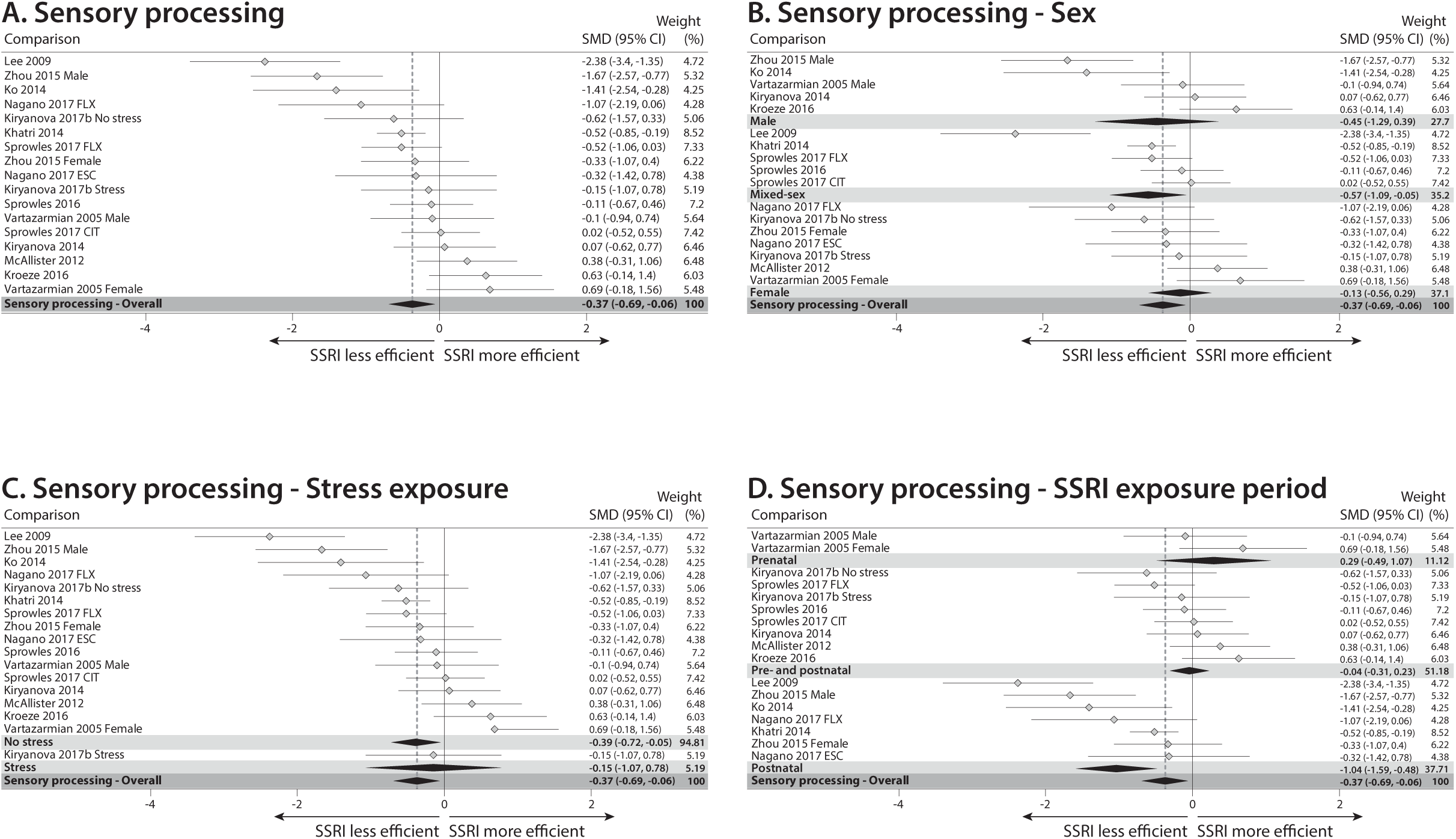
Forest plots of meta-analysis comparing animals perinatally exposed to SSRIs to those exposed to vehicle on the behavioral outcome sensory processing. (A) Overall analysis. (B) Subgroup analysis based on sex. (C) Subgroup analysis based on presence/absence of stress exposure. (D) Subgroup analysis based on SSRI exposure timing.

**Supplementary Figure 10:**
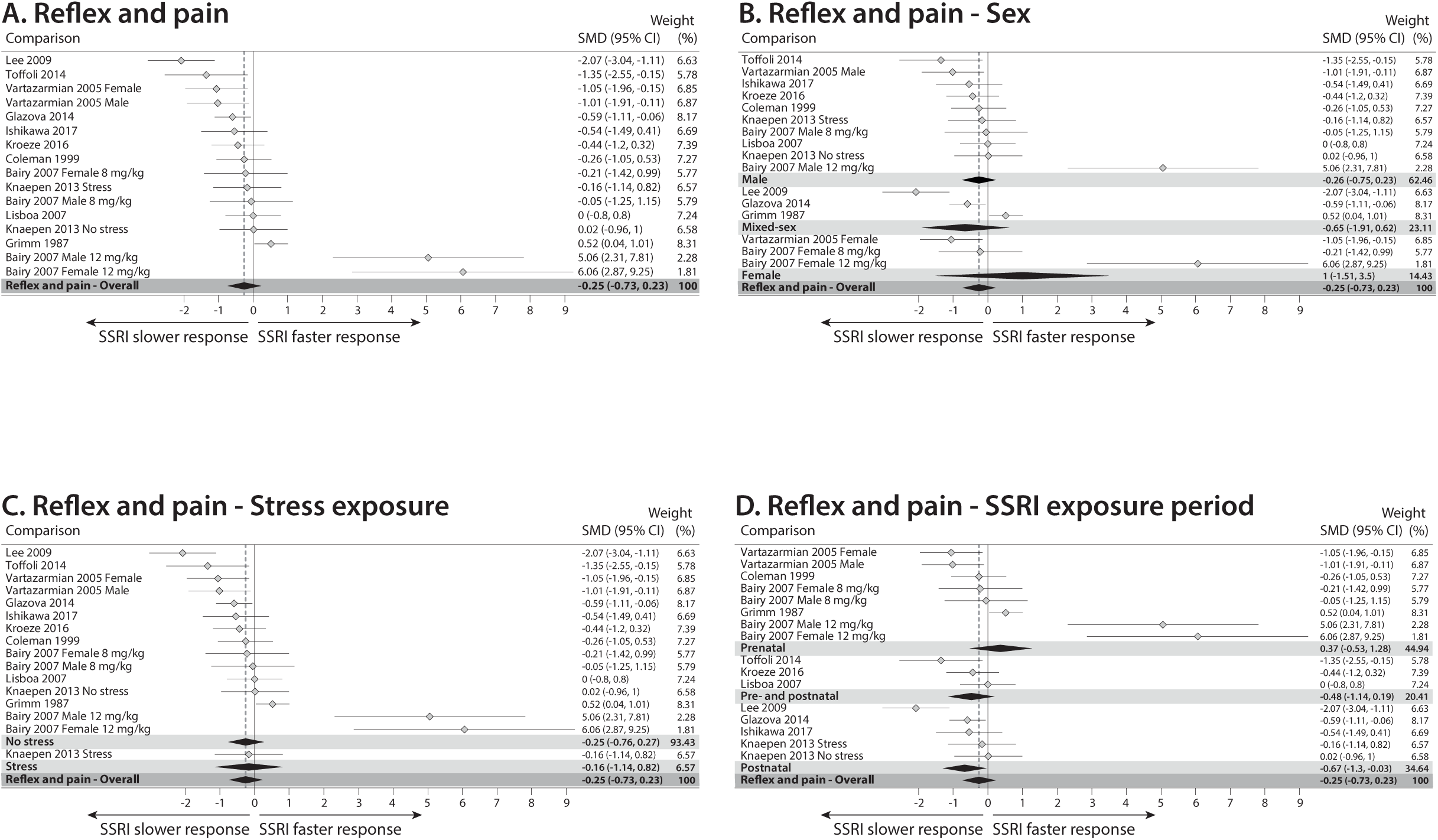
Forest plots of meta-analysis comparing animals perinatally exposed to SSRIs to those exposed to vehicle on the behavioral outcome reflex and pain sensitivity. (A) Overall analysis. (B) Subgroup analysis based on sex. (C) Subgroup analysis based on presence/absence of stress exposure. (D) Subgroup analysis based on SSRI exposure timing.

**Supplementary Figure 11:**
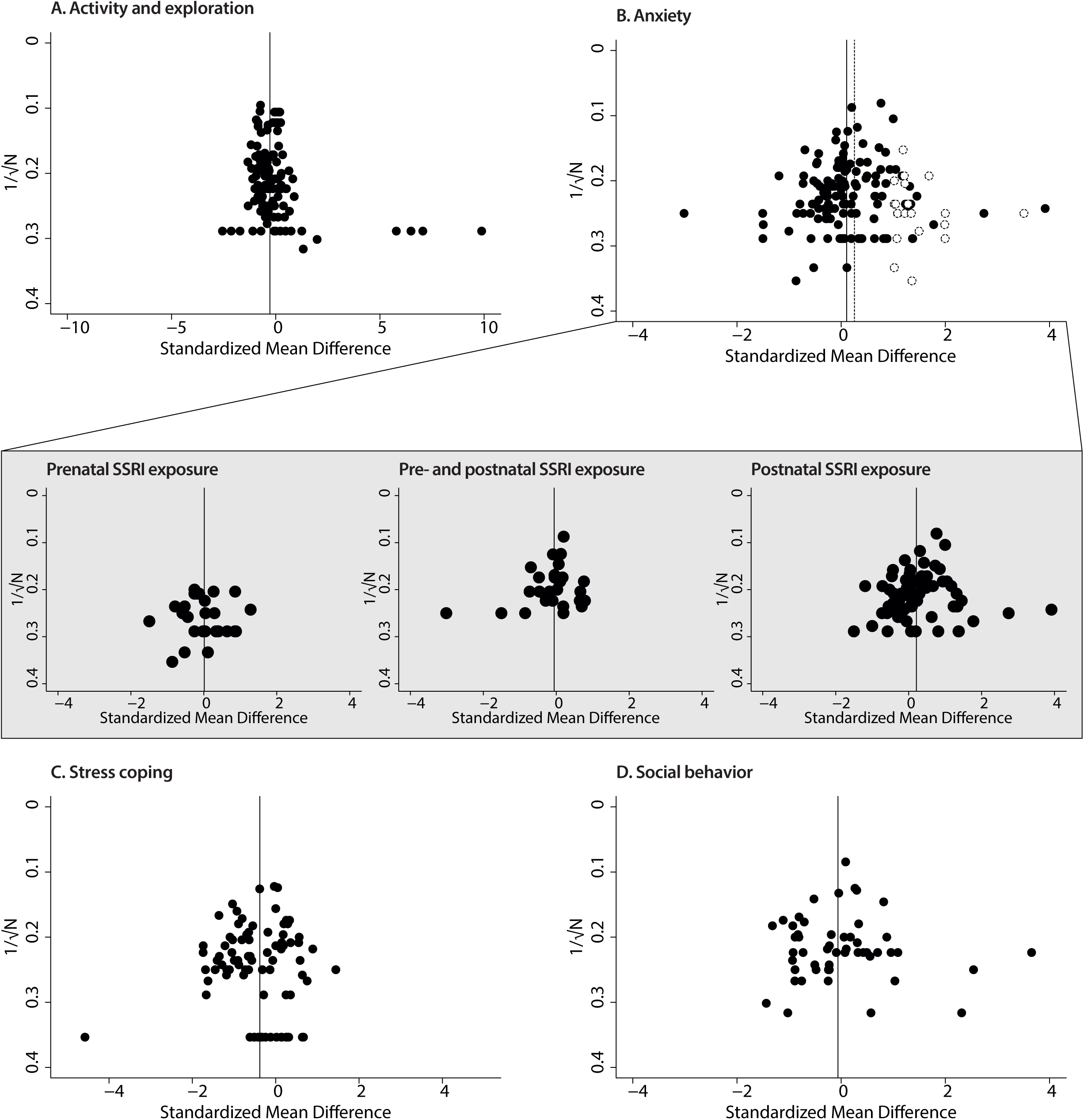
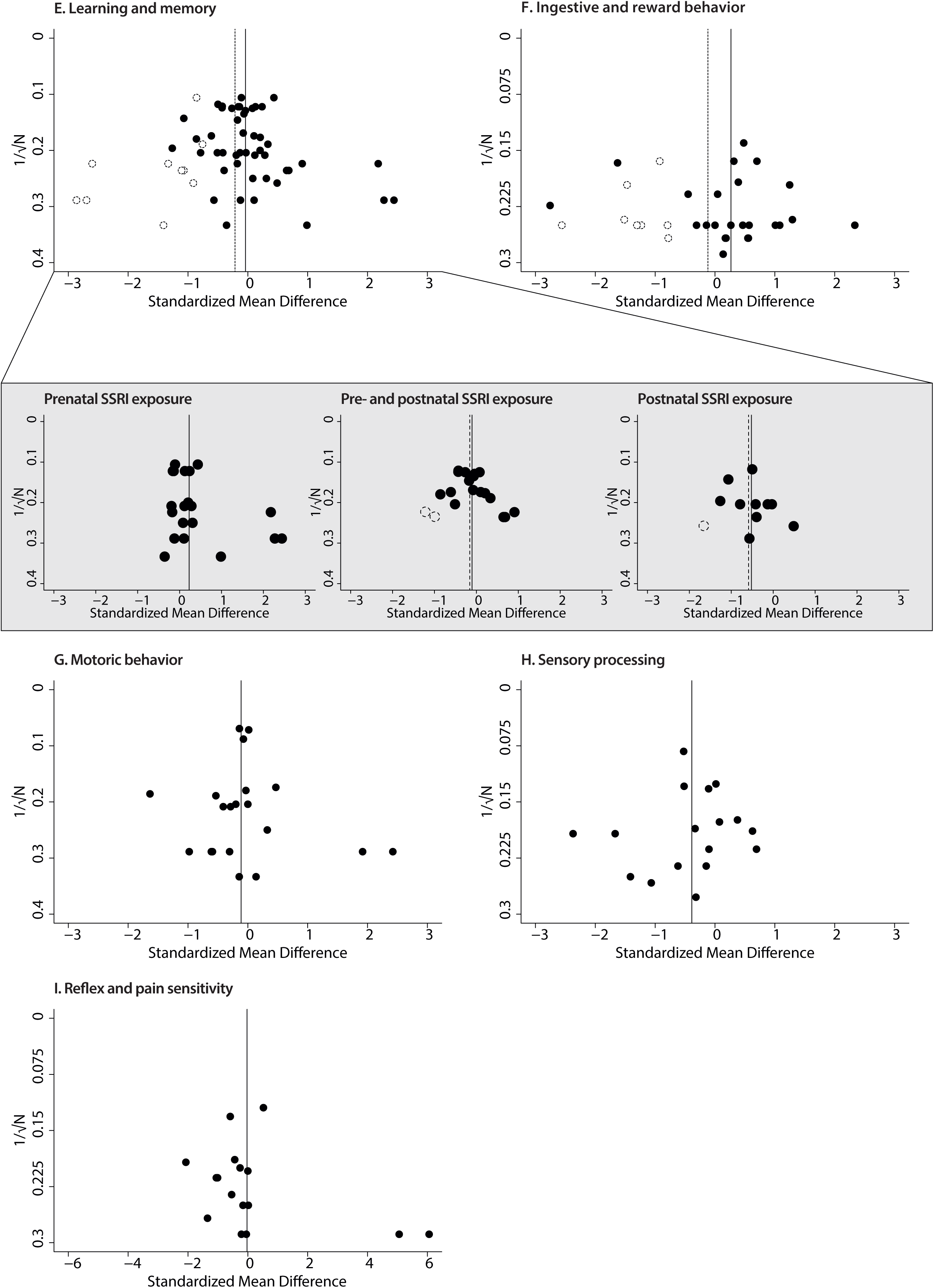
Funnel plots of behavioral outcomes in animals perinatally exposed to SSRIs to those exposed to vehicle on the with imputed extra data points by trim and fill analysis. (A) Activity and exploration. (B) Anxiety. In the gray box the same data separate for each exposure period. (C) Stress coping. (D) Social behavior. (E) Learning and memory. In the gray box the same data separate for each exposure period. (F) Ingestive and reward. (G) Motoric behavior. (H) Sensory processing. (I^2^) Reflex and pain sensitivity.

Supplementary File 1: Systematic search strategy for PubMed, PsycINFO and Web of Science.

Supplementary File 2: Behavioral domains and test prioritization.

## References

1. Bakker, M. K., Kölling, P., van den Berg, P. B., de Walle, H. E. K. & de Jong van den Berg, L. T. W. Increase in use of selective serotonin reuptake inhibitors in pregnancy during the last decade, a population-based cohort study from the Netherlands. Br. J. Clin. Pharmacol. 65, 600–606 (2008).

2. Jimenez-Solem, E. et al. Prevalence of Antidepressant Use during Pregnancy in Denmark, a Nation-Wide Cohort Study. PLoS One 8, e63034 (2013).

3. Cooper, W. O., Willy, M. E., Pont, S. J. & Ray, W. A. Increasing use of antidepressants in pregnancy. Am. J. Obstet. Gynecol. 196, 544.e1-544.e5 (2007).

4. Andrade, S. E. et al. Use of antidepressant medications during pregnancy: a multisite study. Am. J. Obstet. Gynecol. 198, 194.e1–5 (2008).

5. Zoega, H. et al. Use of SSRI and SNRI Antidepressants during Pregnancy: A Population-Based Study from Denmark, Iceland, Norway and Sweden. PLoS One 10, e0144474 (2015).

6. Jordan, S. et al. Selective Serotonin Reuptake Inhibitor (SSRI) Antidepressants in Pregnancy and Congenital Anomalies: Analysis of Linked Databases in Wales, Norway and Funen, Denmark. PLoS One 11, e0165122 (2016).

7. Hayes, R. M. et al. Maternal antidepressant use and adverse outcomes: a cohort study of 228,876 pregnancies. Am. J. Obstet. Gynecol. 207, 49.e1-49.e9 (2012).

8. Huybrechts, K. F. et al. Antidepressant Use Late in Pregnancy and Risk of Persistent Pulmonary Hypertension of the Newborn. JAMA 313, 2142 (2015).

9. Alwan, S., Friedman, J. M. & Chambers, C. Safety of Selective Serotonin Reuptake Inhibitors in Pregnancy: A Review of Current Evidence. CNS Drugs 30, 499–515 (2016).

10. Rampono, J. et al. Placental Transfer of SSRI and SNRI Antidepressants and Effects on the Neonate. Pharmacopsychiatry 42, 95–100 (2009).

11. Gaspar, P., Cases, O. & Maroteaux, L. The developmental role of serotonin: news from mouse molecular genetics. Nat. Rev. Neurosci. 4, 1002–1012 (2003).

12. Muller, C. P. & Jacobs, B. Handbook of the behavioral neurobiology of serotonin. 2009 21, (Academic Press).

13. Teissier, A., Soiza-Reilly, M. & Gaspar, P. Refining the Role of 5-HT in Postnatal Development of Brain Circuits. Front. Cell. Neurosci. 11, 139 (2017).

14. Brummelte, S., Mc Glanaghy, E., Bonnin, A. & Oberlander, T. F. Developmental changes in serotonin signaling: Implications for early brain function, behavior and adaptation. Neuroscience 342, 212–231 (2017).

15. Homberg, J. R., Schubert, D. & Gaspar, P. New perspectives on the neurodevelopmental effects of SSRIs. Trends Pharmacol. Sci. 31, 60–65 (2010).

16. Hanley, G. E., Brain, U. & Oberlander, T. F. Prenatal exposure to serotonin reuptake inhibitor antidepressants and childhood behavior. Pediatr. Res. 78, 174–180 (2015).

17. Hanley, G. E., Brain, U. & Oberlander, T. F. Infant developmental outcomes following prenatal exposure to antidepressants, and maternal depressed mood and positive affect. Early Hum. Dev. 89, 519–524 (2013).

18. Viktorin, A. et al. Association of Antidepressant Medication Use During Pregnancy With Intellectual Disability in Offspring. JAMA Psychiatry 74, 1031 (2017).

19. Hutchison, S. M., Mâsse, L. C., Brain, U. & Oberlander, T. F. A 6-year longitudinal study: Are maternal depressive symptoms and Selective Serotonin Reuptake Inhibitor (SSRI) antidepressant treatment during pregnancy associated with everyday measures of executive function in young children? Early Hum. Dev. 128, 21–26 (2019).

20. Lupattelli, A. et al. Effect of Time-Dependent Selective Serotonin Reuptake Inhibitor Antidepressants During Pregnancy on Behavioral, Emotional, and Social Development in Preschool-Aged Children. J. Am. Acad. Child Adolesc. Psychiatry 57, 200–208 (2018).

21. Oberlander, T. F., Gingrich, J. A. & Ansorge, M. S. Sustained neurobehavioral effects of exposure to SSRI antidepressants during development: molecular to clinical evidence. Clin. Pharmacol. Ther. 86, 672–677 (2009).

22. Johnson, K. C., Smith, A. K., Stowe, Z. N., Newport, D. J. & Brennan, P. A. Preschool outcomes following prenatal serotonin reuptake inhibitor exposure: differences in language and behavior, but not cognitive function. J. Clin. Psychiatry 77, e176–82 (2016).

23. Brown, A. S. et al. Association of Selective Serotonin Reuptake Inhibitor Exposure During Pregnancy With Speech, Scholastic, and Motor Disorders in Offspring. JAMA Psychiatry 73, 1163 (2016).

24. Smearman, E. L. et al. School-age social behavior and pragmatic language ability in children with prenatal serotonin reuptake inhibitor exposure. Dev. Psychopathol. 1–10 (2019). doi:10.1017/S0954579418001372

25. Halvorsen, A., Hesel, B., Østergaard, S. D. & Danielsen, A. A. In utero exposure to selective serotonin reuptake inhibitors and development of mental disorders: a systematic review and meta-analysis. Acta Psychiatr. Scand. 139, 493–507 (2019).

26. Gentile, S. Untreated depression during pregnancy: Short- and long-term effects in offspring. A systematic review. Neuroscience 342, 154–166 (2017).

27. Semple, B. D., Blomgren, K., Gimlin, K., Ferriero, D. M. & Noble-Haeusslein, L. J. Brain development in rodents and humans: Identifying benchmarks of maturation and vulnerability to injury across species. Prog. Neurobiol. 106–107, 1–16 (2013).

28. Zucker, I. Risk mitigation for children exposed to drugs during gestation: A critical role for animal preclinical behavioral testing. Neurosci. Biobehav. Rev. 77, 107–121 (2017).

29. Gingrich, J. A. et al. New Insights into How Serotonin Selective Reuptake Inhibitors Shape the Developing Brain. Birth Defects Res. 109, 924–932 (2017).

30. Glover, M. E. & Clinton, S. M. Of rodents and humans: A comparative review of the neurobehavioral effects of early life SSRI exposure in preclinical and clinical research. Int. J. Dev. Neurosci. 51, 50–72 (2016).

31. Grieb, Z. A. & Ragan, C. M. The effects of perinatal SSRI exposure on anxious behavior and neurobiology in rodent and human offspring. Eur. Neuropsychopharmacol. 1–16 (2019). doi:10.1016/j.euroneuro.2019.07.239

32. Bourke, C. H., Stowe, Z. N. & Owens, M. J. Prenatal Antidepressant Exposure: Clinical and Preclinical Findings. Pharmacol. Rev. 66, 435–465 (2014).

33. Millard, S. J., Weston-Green, K. & Newell, K. A. The effects of maternal antidepressant use on offspring behaviour and brain development: Implications for risk of neurodevelopmental disorders. Neurosci. Biobehav. Rev. 80, 743–765 (2017).

34. Ornoy, A. Neurobehavioral risks of SSRIs in pregnancy: Comparing human and animal data. Reprod. Toxicol. 72, 191–200 (2017).

35. Moher, D., Liberati, A., Tetzlaff, J. & Altman, D. G. Preferred Reporting Items for Systematic Reviews and Meta-Analyses: The PRISMA Statement. PLoS Med. 6, e1000097 (2009).

36. Hooijmans, C. R., Tillema, A., Leenaars, M. & Ritskes-Hoitinga, M. Enhancing search efficiency by means of a search filter for finding all studies on animal experimentation in PubMed. Lab. Anim. 44, 170–175 (2010).

37. Workman, A. D., Charvet, C. J., Clancy, B., Darlington, R. B. & Finlay, B. L. Modeling Transformations of Neurodevelopmental Sequences across Mammalian Species. J. Neurosci. 33, 7368–7383 (2013).

38. Schneider, C. A., Rasband, W. S. & Eliceiri, K. W. NIH Image to ImageJ: 25 years of image analysis. Nat. Methods 9, 671–5 (2012).

39. Hooijmans, C. R. et al. SYRCLE’s risk of bias tool for animal studies. BMC Med. Res. Methodol. 14, 43 (2014).

40. Viechtbauer, W. Conducting Meta-Analyses in R with the metafor Package. J. Stat. Softw. 36, 1–48 (2010).

41. Zwetsloot, P.-P. et al. Standardized mean differences cause funnel plot distortion in publication bias assessments. Elife 6, 1–20 (2017).

42. Duval, S. & Tweedie, R. Trim and fill: A simple funnel-plot-based method of testing and adjusting for publication bias in meta-analysis. Biometrics 56, 455–63 (2000).

43. Grimm, V. E. & Frieder, B. Prenatal and early postnatal exposure to zimelidine: behavioral, neurochemical and histological findings in rats. Int. J. Neurosci. 33, 225–235 (1987).

44. Hilakivi, L. A., Stenberg, D., Sinclair, J. D. & Kiianmaa, K. Neonatal desipramine or zimeldine treatment causes long-lasting changes in brain monoaminergic systems and alcohol related behavior in rats. Psychopharmacology (Berl). 91, 403–409 (1987).

45. Hilakivi, L. A. & Hilakivi, I. Increased adult behavioral ‘despair’ in rats neonatally exposed to desipramine or zimeldine: An animal model of depression? Pharmacol. Biochem. Behav. 28, 367–369 (1987).

46. Hilakivi, L. A., Hilakivi, I. & Kiianmaa, K. Neonatal antidepressant administration suppresses concurrent active (REM) sleep and increases adult alcohol consumption in rats. Alcohol Alcohol Suppl. 1, 339–43 (1987).

47. Hilakivi, L. A., Taira, T. & Hilakivi, I. Early postnatal deprivation of active sleep with desipramine or zimeldine impairs later behavioural reactivity to auditory stimuli in rats. Acta Physiol. Scand. 132, 191–8 (1988).

48. Hilakivi, L. A., Taira, T., Hilakivi, I. & Loikas, P. Neonatal treatment with monoamine uptake inhibitors alters later response in behavioural ‘despair’ test to beta and GABA-B receptor agonists. Pharmacol. Toxicol. 63, 57–61 (1988).

49. Hilakivi, I. Early postnatal antidepressant treatment and juvenile stereotyped behaviour in rats. Eur. J. Pharmacol. 271, 223–226 (1994).

50. Vorhees, C. V et al. A developmental neurotoxicity evaluation of the effects of prenatal exposure to fluoxetine in rats. Fundam. Appl. Toxicol. 23, 194–205 (1994).

51. Frank, M. G. & Heller, H. C. Neonatal treatments with the serotonin uptake inhibitors clomipramine and zimelidine, but not the noradrenaline uptake inhibitor desipramine, disrupt sleep patterns in adult rats. Brain Res. 768, 287–293 (1997).

52. Hansen, H. H., Sanchez, C. & Meier, E. Neonatal administration of the selective serotonin reuptake inhibitor Lu 10-134-C increases forced swimming-induced immobility in adult rats: a putative animal model of depression? J. Pharmacol. Exp. Ther. 283, 1333–1341 (1997).

53. Singh, Y., Jaiswal, A. K., Singh, M. & Bhattacharya, S. K. Effect of prenatal diazepam, phenobarbital, haloperidol and fluoxetine exposure on foot shock induced aggression in rats. Indian J. Exp. Biol. 36, 1023–1024 (1998).

54. Stewart, C. W. et al. Gestational exposure to cocaine or pharmacologically related compounds: effects on behavior and striatal dopamine receptors. Life Sci. 63, 2015–2022 (1998).

55. Coleman, F. H., Christensen, H. D., Gonzalez, C. L. & Rayburn, W. F. Behavioral changes in developing mice after prenatal exposure to paroxetine (Paxil). Am. J. Obstet. Gynecol. 181, 1166–1171 (1999).

56. Christensen, H. D., Rayburn, W. F. & Gonzalez, C. L. Chronic prenatal exposure to paroxetine (Paxil) and cognitive development of mice offspring. Neurotoxicol. Teratol. 22, 733–739 (2000).

57. Mendes-da-Silva, C. et al. Neonatal treatment with fluoxetine reduces depressive behavior induced by forced swim in adult rats. Arq. Neuropsiquiatr. 60, 928–31 (2002).

58. Ansorge, M. S., Zhou, M., Lira, A., Hen, R. & Gingrich, J. A. Early-Life Blockade of the 5-HT Transporter Alters Emotional Behavior in Adult Mice. Science (80-.). 306, 879–881 (2004).

59. Ishiwata, H., Shiga, T. & Okado, N. Selective serotonin reuptake inhibitor treatment of early postnatal mice reverses their prenatal stress-induced brain dysfunction. Neuroscience 133, 893–901 (2005).

60. Vartazarmian, R., Malik, S., Baker, G. B. & Boksa, P. Long-term effects of fluoxetine or vehicle administration during pregnancy on behavioral outcomes in guinea pig offspring. Psychopharmacology (Berl). 178, 328–338 (2005).

61. Deiro, T. C. B. J. et al. Sertraline delays the somatic growth and reflex ontogeny in neonate rats. Physiol. Behav. 87, 338–344 (2006).

62. Maciag, D. et al. Neonatal Antidepressant Exposure has Lasting Effects on Behavior and Serotonin Circuitry. Neuropsychopharmacology 31, 47–57 (2006).

63. Maciag, D., Williams, L., Coppinger, D. & Paul, I. A. Neonatal citalopram exposure produces lasting changes in behavior which are reversed by adult imipramine treatment. Eur. J. Pharmacol. 532, 265–269 (2006).

64. Maciag, D., Coppinger, D. & Paul, I. A. Evidence that the deficit in sexual behavior in adult rats neonatally exposed to citalopram is a consequence of 5-HT1 receptor stimulation during development. BRAIN Res. 1125, 171–175 (2006).

65. Bairy, K. L., Madhyastha, S., Ashok, K. P., Bairy, I. & Malini, S. Developmental and behavioral consequences of prenatal fluoxetine. Pharmacology 79, 1–11 (2007).

66. Lisboa, S. F. S., Oliveira, P. E., Costa, L. C., Venancio, E. J. & Moreira, E. G. Behavioral evaluation of male and female mice pups exposed to fluoxetine during pregnancy and lactation. Pharmacology 80, 49–56 (2007).

67. Ansorge, M. S., Morelli, E. & Gingrich, J. A. Inhibition of serotonin but not norepinephrine transport during development produces delayed, persistent perturbations of emotional behaviors in mice. J. Neurosci. 28, 199–207 (2008).

68. Cagiano, R. Neurofunctional effects in rats prenatally exposed to fluoxetine. Eur. Rev. Med. Pharmacol. Sci. 12, 137–148 (2008).

69. Deiró, T. C. B. D. J. et al. Neonatal exposure to citalopram, a serotonin selective reuptake inhibitor, programs a delay in the reflex ontogeny in rats. Arq. Neuropsiquiatr. 66, 736–40 (2008).

70. Favaro, P. das N., Costa, L. C. & Moreira, E. G. Maternal fluoxetine treatment decreases behavioral response to dopaminergic drugs in female pups. Neurotoxicol. Teratol. 30, 487–494 (2008).

71. Forcelli, P. A. & Heinrichs, S. C. Teratogenic effects of maternal antidepressant exposure on neural substrates of drug-seeking behavior in offspring. Addict. Biol. 13, 52–62 (2008).

72. Gouvêa, T. S., Morimoto, H. K., de Faria, M. J. S. S., Moreira, E. G. & Gerardin, D. C. C. Maternal exposure to the antidepressant fluoxetine impairs sexual motivation in adult male mice. Pharmacol. Biochem. Behav. 90, 416–9 (2008).

73. Noorlander, C. W. et al. Modulation of serotonin transporter function during fetal development causes dilated heart cardiomyopathy and lifelong behavioral abnormalities. PLoS One 3, e2782 (2008).

74. Popa, D., Léna, C., Alexandre, C. & Adrien, J. Lasting syndrome of depression produced by reduction in serotonin uptake during postnatal development: evidence from sleep, stress, and behavior. J. Neurosci. 28, 3546–54 (2008).

75. Jiang, X.-Z. et al. [Neonatal fluoxetine exposure induced depression-like behaviors in adult Kunming mice and the antidepressant-like effect of agmatine]. Yao Xue Xue Bao 44, 716–21 (2009).

76. Karpova, N. N., Lindholm, J., Pruunsild, P., Timmusk, T. & Castren, E. Long-lasting behavioural and molecular alterations induced by early postnatal fluoxetine exposure are restored by chronic fluoxetine treatment in adult mice. Eur. Neuropsychopharmacol. 19, 97–108 (2009).

77. Lee, L.-J. Neonatal fluoxetine exposure affects the neuronal structure in the somatosensory cortex and somatosensory-related behaviors in adolescent rats. Neurotox. Res. 15, 212–23 (2009).

78. Capello, C. F. et al. Serotonin transporter occupancy in rats exposed to serotonin reuptake inhibitors in utero or via breast milk. J. Pharmacol. Exp. Ther. 339, 275–85 (2011).

79. Mnie-Filali, O. et al. Pharmacological blockade of 5-HT7 receptors as a putative fast acting antidepressant strategy. Neuropsychopharmacology 36, 1275–1288 (2011).

80. Olivier, J. D. A. A. et al. Fluoxetine administration to pregnant rats increases anxiety-related behavior in the offspring. Psychopharmacology (Berl). 217, 419–432 (2011).

81. Pivina, S. G., Fedotova, Y. O., Akulova, V. K. & Ordyan, N. E. Effects of selective inhibitors of serotonin reuptake on the anxiety behavior and activity of the pituitary-adrenal system in prenatally stressed male rats. Neurochem. J. 5, 47–51 (2011).

82. Rayen, I., van den Hove, D. L., Prickaerts, J., Steinbusch, H. W. & Pawluski, J. L. Fluoxetine during Development Reverses the Effects of Prenatal Stress on Depressive-Like Behavior and Hippocampal Neurogenesis in Adolescence. PLoS One 6, e24003 (2011).

83. Rodriguez-Porcel, F. et al. Neonatal exposure of rats to antidepressants affects behavioral reactions to novelty and social interactions in a manner analogous to autistic spectrum disorders. Anat. Rec. (Hoboken). 294, 1726–35 (2011).

84. Simpson, K. L. et al. Perinatal antidepressant exposure alters cortical network function in rodents. Proc. Natl. Acad. Sci. 108, 18465–18470 (2011).

85. Zheng, J. et al. Neonatal exposure to fluoxetine and fluvoxamine alteres spine density in mouse hippocampal CA1 pyramidal neurons. Int. J. Clin. Exp. Pathol. 4, 162–8 (2011).

86. Harris, S. S., Maciag, D., Simpson, K. L., Lin, R. C. S. S. & Paul, I. A. Dose-dependent effects of neonatal SSRI exposure on adult behavior in the rat. Brain Res. 1429, 52–60 (2012).

87. Swilley-Harris, S. Dose-dependent Effects of Neonatal SSRI Exposure on Behavior and Brain Serotonin Turnover In the Adult Rat. (The University of Mississippi Medical Center, 2010).

88. Kummet, G. J. et al. Neonatal SSRI Exposure Programs a Hypermetabolic State in Adult Mice. J. Nutr. Metab. 2012, 1–8 (2012).

89. Lee, L.-J. & Lee, L. J.-H. Neonatal fluoxetine exposure alters motor performances of adolescent rats. Dev. Neurobiol. 72, 1122–32 (2012).

90. McAllister, B. B., Kiryanova, V. & Dyck, R. H. Behavioural outcomes of perinatal maternal fluoxetine treatment. Neuroscience 226, 356–66 (2012).

91. Nagano, M., Liu, M., Inagaki, H., Kawada, T. & Suzuki, H. Early intervention with fluoxetine reverses abnormalities in the serotonergic system and behavior of rats exposed prenatally to dexamethasone. Neuropharmacology 63, 292–300 (2012).

92. Rebello, T. J. Serotonin Modulates the Maturation of the Medial Prefrontal Cortex and Hippocampus: Relevance to the Etiology of Emotional and Cognitive Behaviors. (Columbia University, 2012).

93. Smit-Rigter, L. A. et al. Prenatal fluoxetine exposure induces life-long serotonin 5-HT₃ receptor-dependent cortical abnormalities and anxiety-like behaviour. Neuropharmacology 62, 865–70 (2012).

94. Soga, T., Wong, D. W., Putteeraj, M., Song, K. P. & Parhar, I. S. Early-life citalopram-induced impairments in sexual behavior and the role of androgen receptor. Neuroscience 225, 172–184 (2012).

95. Yu, Q. Developmental monoamine signaling impacts adult affective and aggressive behaviors. (Columbia University, 2012).

96. Bourke, C. H., Stowe, Z. N., Neigh, G. N., Olson, D. E. & Owens, M. J. Prenatal exposure to escitalopram and/or stress in rats produces limited effects on endocrine, behavioral, or gene expression measures in adult male rats. Neurotoxicol. Teratol. 39, 100–109 (2013).

97. Francis-Oliveira, J. et al. Fluoxetine exposure during pregnancy and lactation: Effects on acute stress response and behavior in the novelty-suppressed feeding are age and gender-dependent in rats. Behav. Brain Res. 252, 195–203 (2013).

98. Freund, N., Thompson, B. S., DeNormandie, J., Vaccarro, K. & Andersen, S. L. Windows of vulnerability: Maternal separation, age, and fluoxetine on adolescent depressive-like behavior in rats. Neuroscience 249, 88–97 (2013).

99. Kiryanova, V., Smith, V. M., Dyck, R. H. & Antle, M. C. The effects of perinatal fluoxetine treatment on the circadian system of the adult mouse. Psychopharmacology (Berl). 225, 743–51 (2013).

100. Knaepen, L. et al. Developmental Fluoxetine Exposure Normalizes the Long-Term Effects of Maternal Stress on Post-Operative Pain in Sprague-Dawley Rat Offspring. PLoS One 8, e57608 (2013).

101. Rayen, I., Steinbusch, H. W. M., Charlier, T. D. & Pawluski, J. L. Developmental fluoxetine exposure and prenatal stress alter sexual differentiation of the brain and reproductive behavior in male rat offspring. Psychoneuroendocrinology 38, 1618–29 (2013).

102. Schaefer, T. L. et al. Cognitive impairments from developmental exposure to serotonergic drugs: citalopram and MDMA. Int. J. Neuropsychopharmacol. 16, 1383–94 (2013).

103. Vieira, M. L. et al. Could maternal exposure to the antidepressants fluoxetine and St. John’s Wort induce long-term reproductive effects on male rats? Reprod. Toxicol. 35, 102–7 (2013).

104. da Silva, A. I. et al. Fluoxetine treatment of rat neonates significantly reduces oxidative stress in the hippocampus and in behavioral indicators of anxiety later in postnatal life. Can. J. Physiol. Pharmacol. 92, 330–7 (2014).

105. Glazova, N. Y. et al. Effects of neonatal fluvoxamine administration on the physical development and activity of the serotoninergic system in white rats. Acta Naturae 6, 98–105 (2014).

106. Khatri, N., Simpson, K. L., Lin, R. C. S. & Paul, I. A. Lasting neurobehavioral abnormalities in rats after neonatal activation of serotonin 1A and 1B receptors: possible mechanisms for serotonin dysfunction in autistic spectrum disorders. Psychopharmacology (Berl). 231, 1191–200 (2014).

107. Khatri, N. Role of Early Life Stimulation of Serotonin 1A and 1B (5-HT1A and 5-HT1B) Receptors in the Lasting Neurobehavioral Effects of neonatal SSRI Exposure. (The University of Mississippi Medical Center, 2013).

108. Kiryanova, V. & Dyck, R. H. Increased Aggression, Improved Spatial Memory, and Reduced Anxiety-Like Behaviour in Adult Male Mice Exposed to Fluoxetine Early in Life. Dev. Neurosci. 36, 396–408 (2014).

109. Ko, M.-C., Lee, L. J.-H., Li, Y. & Lee, L.-J. Long-term consequences of neonatal fluoxetine exposure in adult rats. Dev. Neurobiol. 74, 1038–1051 (2014).

110. Rayen, I., Steinbusch, H. W. M., Charlier, T. D. & Pawluski, J. L. Developmental fluoxetine exposure facilitates sexual behavior in female offspring. Psychopharmacology (Berl). 231, 123–133 (2014).

111. Rebello, T. J. et al. Postnatal Day 2 to 11 Constitutes a 5-HT-Sensitive Period Impacting Adult mPFC Function. J. Neurosci. 34, 12379–12393 (2014).

112. Sarkar, A. et al. Hippocampal HDAC4 Contributes to Postnatal Fluoxetine-Evoked Depression-Like Behavior. Neuropsychopharmacology 39, 2221–2232 (2014).

113. Sarkar, A., Chachra, P. & Vaidya, V. A. Postnatal fluoxetine-evoked anxiety is prevented by concomitant 5-HT2A/C receptor blockade and mimicked by postnatal 5-HT2A/C receptor stimulation. Biol. Psychiatry 76, 858–68 (2014).

114. Toffoli, L. V et al. Maternal exposure to fluoxetine during gestation and lactation affects the DNA methylation programming of rat’s offspring: modulation by folic acid supplementation. Behav. Brain Res. 265, 142–147 (2014).

115. Volodina, M. A. et al. [Effects of neonatal fluvoxamine administration to white rats and their correction by semax treatment]. Izv. Akad. Nauk. Seriia Biol. 391–7 (2014).

116. Yu, Q. et al. Dopamine and serotonin signaling during two sensitive developmental periods differentially impact adult aggressive and affective behaviors in mice. Mol. Psychiatry 19, 688–698 (2014).

117. Altieri, S. C. et al. Perinatal vs genetic programming of serotonin states associated with anxiety. Neuropsychopharmacology 40, 1456–70 (2015).

118. Avitsur, R. et al. Prenatal fluoxetine exposure affects cytokine and behavioral response to an immune challenge. J. Neuroimmunol. 284, 49–56 (2015).

119. da Silva, A. I. et al. Fluoxetine induces lean phenotype in rat by increasing the brown/white adipose tissue ratio and UCP1 expression. J. Bioenerg. Biomembr. 47, 309–318 (2015).

120. Ehrlich, D. E. et al. Prenatal stress, regardless of concurrent escitalopram treatment, alters behavior and amygdala gene expression of adolescent female rats. Neuropharmacology 97, 251–258 (2015).

121. Galindo, L. C. M. et al. Neonatal serotonin reuptake inhibition reduces hypercaloric diet effects on fat mass and hypothalamic gene expression in adult rats. Int. J. Dev. Neurosci. 46, 76–81 (2015).

122. Zhou, X. et al. Behavioral training reverses global cortical network dysfunction induced by perinatal antidepressant exposure. Proc. Natl. Acad. Sci. 112, 2233–2238 (2015).

123. Boulle, F. et al. Prenatal stress and early-life exposure to fluoxetine have enduring effects on anxiety and hippocampal BDNF gene expression in adult male offspring. Dev. Psychobiol. 58, 427–438 (2016).

124. Boulle, F. et al. Developmental fluoxetine exposure increases behavioral despair and alters epigenetic regulation of the hippocampal BDNF gene in adult female offspring. Horm. Behav. 80, 47–57 (2016).

125. Dos Santos, A. H., et al. In utero and lactational exposure to fluoxetine delays puberty onset in female rats offspring. Reprod. Toxicol. 62, 1–8 (2016).

126. Gobinath, A. R., Workman, J. L., Chow, C., Lieblich, S. E. & Galea, L. A. M. M. Maternal postpartum corticosterone and fluoxetine differentially affect adult male and female offspring on anxiety-like behavior, stress reactivity, and hippocampal neurogenesis. Neuropharmacology 101, 165–178 (2016).

127. Kiryanova, V., Meunier, S. J., Vecchiarelli, H. A., Hill, M. N. & Dyck, R. H. Effects of maternal stress and perinatal fluoxetine exposure on behavioral outcomes of adult male offspring. Neuroscience 320, 281–96 (2016).

128. Kroeze, Y. et al. Perinatal reduction of functional serotonin transporters results in developmental delay. Neuropharmacology 109, 96–111 (2016).

129. Matsumoto, A. K. et al. Co-exposure to fish oil or folic acid does not reverse effects in the progeny induced by maternal exposure to fluoxetine. Neurotoxicol. Teratol. 56, 1–8 (2016).

130. Salari, A.-A., Fatehi-Gharehlar, L., Motayagheni, N. & Homberg, J. R. Fluoxetine normalizes the effects of prenatal maternal stress on depression- and anxiety-like behaviors in mouse dams and male offspring. Behav. Brain Res. 311, 354–367 (2016).

131. Sprowles, J. L. N. et al. Perinatal exposure to the selective serotonin reuptake inhibitor citalopram alters spatial learning and memory, anxiety, depression, and startle in Sprague-Dawley rats. Int. J. Dev. Neurosci. 54, 39–52 (2016).

132. Svirsky, N., Levy, S. & Avitsur, R. Prenatal exposure to selective serotonin reuptake inhibitors (SSRI) increases aggression and modulates maternal behavior in offspring mice. Dev. Psychobiol. 58, 71–82 (2016).

133. Zohar, I., Shoham, S. & Weinstock, M. Perinatal citalopram does not prevent the effect of prenatal stress on anxiety, depressive-like behaviour and serotonergic transmission in adult rat offspring. Eur. J. Neurosci. 43, 590–600 (2016).

134. Avitsur, R. Increased symptoms of illness following prenatal stress: Can it be prevented by fluoxetine? Behav. Brain Res. 317, 62–70 (2017).

135. Gemmel, M. et al. Perinatal fluoxetine effects on social play, the HPA system, and hippocampal plasticity in pre-adolescent male and female rats: Interactions with pre-gestational maternal stress. Psychoneuroendocrinology 84, 159–171 (2017).

136. Haskell, S. E. et al. Cardiac Outcomes After Perinatal Sertraline Exposure in Mice. J. Cardiovasc. Pharmacol. 70, 119–127 (2017).

137. Ishikawa, C. & Shiga, T. The postnatal 5-HT1A receptor regulates adult anxiety and depression differently via multiple molecules. Prog. Neuro-Psychopharmacology Biol. Psychiatry 78, 66–74 (2017).

138. Kiryanova, V., Smith, V. M., Dyck, R. H. & Antle, M. C. Circadian behavior of adult mice exposed to stress and fluoxetine during development. Psychopharmacology (Berl). 234, 793–804 (2017).

139. Kiryanova, V., Meunier, S. J. & Dyck, R. H. Behavioural outcomes of adult female offspring following maternal stress and perinatal fluoxetine exposure. Behav. Brain Res. 331, 84–91 (2017).

140. Nagano, R., Nagano, M., Nakai, A., Takeshita, T. & Suzuki, H. Differential effects of neonatal SSRI treatments on hypoxia-induced behavioral changes in male and female offspring. Neuroscience 360, 95–105 (2017).

141. Pinheiro, I. L. et al. Neonatal fluoxetine exposure modulates serotonergic neurotransmission and disturb inhibitory action of serotonin on food intake. Behav. Brain Res. 357–358, 65–70 (2019).

142. Sprowles, J. L. N. et al. Differential effects of perinatal exposure to antidepressants on learning and memory, acoustic startle, anxiety, and open-field activity in Sprague-Dawley rats. Int. J. Dev. Neurosci. 61, 92–111 (2017).

143. Meyer, L. R. et al. Perinatal SSRI exposure permanently alters cerebral serotonin receptor mRNA in mice but does not impact adult behaviors. J. Matern. Neonatal Med. 31, 1393–1401 (2018).

144. Sterne, J. A. C. et al. Recommendations for examining and interpreting funnel plot asymmetry in meta-analyses of randomised controlled trials. BMJ 343, d4002–d4002 (2011).

145. Pawluski, J. L., Li, M. & Lonstein, J. S. Serotonin and motherhood: From molecules to mood. Front. Neuroendocrinol. 53, 100742 (2019).

146. Bonnin, A., Torii, M., Wang, L., Rakic, P. & Levitt, P. Serotonin modulates the response of embryonic thalamocortical axons to netrin-1. Nat. Neurosci. 10, 588–597 (2007).

147. Bonnin, A. et al. A transient placental source of serotonin for the fetal forebrain. Nature 472, 347–350 (2011).

148. Xu, Y., Sari, Y. & Zhou, F. C. Selective serotonin reuptake inhibitor disrupts organization of thalamocortical somatosensory barrels during development. Dev. Brain Res. 150, 151–161 (2004).

149. Hodes, G. E. & Epperson, C. N. Sex Differences in Vulnerability and Resilience to Stress Across the Life Span. Biol. Psychiatry 86, 421–432 (2019).

150. Connell, S., Karikari, C. & Hohmann, C. F. Sex-specific development of cortical monoamine levels in mouse. Dev. Brain Res. 151, 187–191 (2004).

151. Slattery, D. A. & Cryan, J. F. Using the rat forced swim test to assess antidepressant-like activity in rodents. Nat. Protoc. 7, 1009–1014 (2012).

152. Porsolt, R. D., Le Pichon, M. & Jalfre, M. Depression: a new animal model sensitive to antidepressant treatments. Nature 266, 730–2 (1977).

153. Commons, K. G., Cholanians, A. B., Babb, J. A. & Ehlinger, D. G. The Rodent Forced Swim Test Measures Stress-Coping Strategy, Not Depression-like Behavior. ACS Chem. Neurosci. 8, 955–960 (2017).

154. Hornix, B. E., Havekes, R. & Kas, M. J. H. Multisensory cortical processing and dysfunction across the neuropsychiatric spectrum. Neurosci. Biobehav. Rev. 97, 138–151 (2019).

155. Malm, H. et al. Gestational Exposure to Selective Serotonin Reuptake Inhibitors and Offspring Psychiatric Disorders: A National Register-Based Study. J. Am. Acad. Child Adolesc. Psychiatry 55, 359–366 (2016).

156. Sorensen, M. J. et al. Antidepressant exposure in pregnancy and risk of autism spectrum disorders. Clin. Epidemiol. 5, 449 (2013).

157. Avey, M. T., et al. The Devil Is in the Details: Incomplete Reporting in Preclinical Animal Research. PLoS One 11, e0166733 (2016).

158. Kilkenny, C., et al. Survey of the Quality of Experimental Design, Statistical Analysis and Reporting of Research Using Animals. PLoS One 4, (2009).

159. Rotem-Kohavi, N. & Oberlander, T. F. Variations in Neurodevelopmental Outcomes in Children with Prenatal SSRI Antidepressant Exposure. Birth Defects Res. 109, 909–923 (2017).

160. Jansen of Lorkeers, S. J., Doevendans, P. A. & Chamuleau, S. A. J. All preclinical trials should be registered in advance in an online registry. Eur. J. Clin. Invest. 44, 891–892 (2014).

161. Kilkenny, C., Browne, W. J., Cuthill, I. C., Emerson, M. & Altman, D. G. Improving Bioscience Research Reporting: The ARRIVE Guidelines for Reporting Animal Research. PLoS Biol. 8, e1000412 (2010).

162. Muhlhausler, B. S., Bloomfield, F. H. & Gillman, M. W. Whole Animal Experiments Should Be More Like Human Randomized Controlled Trials. PLoS Biol. 11, e1001481 (2013).

163. Clayton, J. A. & Collins, F. S. Policy: NIH to balance sex in cell and animal studies. Nature 509, 282–283 (2014).

164. Glover, M. E. et al. Early-life exposure to the SSRI paroxetine exacerbates depression-like behavior in anxiety/depression-prone rats. Neuroscience 284, 775–797 (2015).

