## Supplementary File 1 for "Perinatal selective serotonin reuptake inhibitor exposure and behavioral outcomes: a systematic review and meta-analyses of animal studies"

*SYRCLE animal filter used for PubMed and adapted for PsycINFO and Web of Science

**PubMed:**

(perinatal[tiab] OR peri-natal[tiab] OR prenatal[tiab] OR pre-natal[tiab] OR postnatal[tiab] OR post-natal[tiab] OR antepartum[tiab] OR ante-partum[tiab] OR ante partum[tiab] OR antepartal[tiab] OR prepartum[tiab] OR pre-partum[tiab] OR prepartal[tiab] OR pre-partal[tiab] OR neonatal[tiab] OR neo-natal[tiab] OR neonate[tiab] OR neonates[tiab] OR prenatal exposure delayed effects[Mesh] OR prenatal injuries[Mesh] OR adolescent[tiab] OR adolescents[tiab] OR adolescence[tiab] OR juvenile[tiab] OR newborn[tiab] OR newborns[tiab] OR infant[tiab] OR infants[tiab] OR infantile[tiab] OR pregnancy[tiab] OR pregnancy[Mesh:NoExp] OR pregnant[tiab] OR pregnancies[tiab] OR pregnancy complications[Mesh:NoExp] OR maternal-fetal exchange[Mesh] OR maternal-fetal[tiab] OR maternal-foetal[tiab] OR transplacental[tiab] OR in utero[tiab] OR gestation[tiab] OR gestational[tiab] OR early-life[tiab] OR development[tiab] OR developmental[tiab] OR developing[tiab] OR fetus[tiab] OR fetuses[tiab ] OR fetus[Mesh:NoExp] OR foetus[tiab] OR foetuses[tiab] OR fetal[tiab] OR embryo[tiab] OR embryos[tiab] OR embryonic[tiab] OR embryology[tiab] OR embryology[MesH:NoExp] OR embryology[subheading] OR embryogenesis[tiab] OR embryo-genesis[tiab] OR embryonic and fetal development[Mesh:NoExp] OR fetal development[Mesh:NoExp] OR embryonic development[Mesh:NoExp] OR growth and development[Mesh:NoExp] OR age factors[Mesh:NoExp])

AND

(SSRI[tiab] OR SSRIs[tiab] OR Selective Serotonin Uptake Inhibitor[tiab] OR Selective Serotonin Uptake Inhibitors[tiab] OR Selective Serotonin Reuptake Inhibitor[tiab] OR Selective Serotonin Reuptake Inhibitors[tiab] OR Serotonin Uptake Inhibitor[tiab] OR Serotonin Uptake Inhibitors[tiab] OR Serotonin Uptake Inhibitors[Pharmacological Action] OR Serotonin Uptake Inhibitors[Mesh] OR Uptake Inhibitor, Serotonin[tiab] OR Uptake Inhibitors, Serotonin[tiab] OR Serotonin Reuptake Inhibitor[tiab] OR Serotonin Reuptake Inhibitors[tiab] OR Reuptake Inhibitor, Serotonin[tiab] OR Reuptake Inhibitors, Serotonin[tiab] OR 5-Hydroxytryptamine Uptake Inhibitor[tiab] OR 5-Hydroxytryptamine Uptake Inhibitors[tiab] OR 5 Hydroxytryptamine Uptake Inhibitor[tiab] OR 5 Hydroxytryptamine Uptake Inhibitors[tiab] OR 5-HT Uptake Inhibitor[tiab] OR 5-HT Uptake Inhibitors[tiab] OR Uptake Inhibitor, 5-HT[tiab] OR Uptake Inhibitors, 5-HT[tiab] OR 5 HT Uptake Inhibitor[tiab] OR 5 HT Uptake Inhibitors[tiab] OR vilazodone[tiab] OR dapoxetine[tiab] OR Citalopram[tiab] OR femoxetine[tiab] OR Fluoxetine[tiab] OR Fluvoxamine[tiab] OR indalpine[tiab] OR Paroxetine[tiab] OR Sertraline[tiab] OR Zimeldine[tiab] OR vilazodone hydrochloride[Mesh] OR dapoxetine[supplementary concept] OR Citalopram[Mesh] OR femoxetine[Supplementary Concept] OR Fluoxetine[Mesh] OR Fluvoxamine[Mesh] OR indalpine[Supplementary Concept] OR Paroxetine[Mesh] OR Sertraline[Mesh] OR Zimeldine[Mesh] OR Priligy[tiab] OR Cytalopram[tiab] OR Escitalopram[tiab] OR Lexapro[tiab] OR femoxitine[tiab] OR Fluoxetin[tiab] OR Sarafem[tiab] OR Prozac[tiab] OR Fluvoxamin[tiab] OR Fluvoxamina[tiab] OR Luvox[tiab] OR Fevarin[tiab] OR Floxyfral[tiab] OR Dumirox[tiab] OR Faverin[tiab] OR Upstene[tiab] OR Seroxat[tiab] OR Paxil[tiab] OR Aropax[tiab] OR Zoloft[tiab] OR Lustral[tiab] OR Aremis[tiab] OR Besitran[tiab] OR Zimelidine[tiab] OR Zimelidin[tiab] OR Zelmid[tiab])

AND

("animal experimentation"[MeSH Terms] OR "models, animal"[MeSH Terms] OR "invertebrates"[MeSH Terms] OR "Animals"[Mesh:noexp] OR "animal population groups"[MeSH Terms] OR "chordata"[MeSH Terms:noexp] OR "chordata, nonvertebrate"[MeSH Terms] OR "vertebrates"[MeSH Terms:noexp] OR "amphibians"[MeSH Terms] OR "birds"[MeSH Terms] OR "fishes"[MeSH Terms] OR "reptiles"[MeSH Terms] OR "mammals"[MeSH Terms:noexp] OR "primates"[MeSH Terms:noexp] OR "artiodactyla"[MeSH Terms] OR "carnivora"[MeSH Terms] OR "cetacea"[MeSH Terms] OR "chiroptera"[MeSH Terms] OR "elephants"[MeSH Terms] OR "hyraxes"[MeSH Terms] OR "insectivora"[MeSH Terms] OR "lagomorpha"[MeSH Terms] OR "marsupialia"[MeSH Terms] OR "monotremata"[MeSH Terms] OR "perissodactyla"[MeSH Terms] OR "rodentia"[MeSH Terms] OR "scandentia"[MeSH Terms] OR "sirenia"[MeSH Terms] OR "xenarthra"[MeSH Terms] OR "haplorhini"[MeSH Terms:noexp] OR "strepsirhini"[MeSH Terms] OR "platyrrhini"[MeSH Terms] OR "tarsii"[MeSH Terms] OR "catarrhini"[MeSH Terms:noexp] OR "cercopithecidae"[MeSH Terms] OR "hylobatidae"[MeSH Terms] OR "hominidae"[MeSH Terms:noexp] OR "gorilla gorilla"[MeSH Terms] OR "pan paniscus"[MeSH Terms] OR "pan troglodytes"[MeSH Terms] OR "pongo pygmaeus"[MeSH Terms]) OR ((animals[tiab] OR animal[tiab] OR mice[Tiab] OR mus[Tiab] OR mouse[Tiab] OR murine[Tiab] OR woodmouse[tiab] OR rats[Tiab] OR rat[Tiab] OR murinae[Tiab] OR muridae[Tiab] OR cottonrat[tiab] OR cottonrats[tiab] OR hamster[tiab] OR hamsters[tiab] OR cricetinae[tiab] OR rodentia[Tiab] OR rodent[Tiab] OR rodents[Tiab] OR pigs[Tiab] OR pig[Tiab] OR swine[tiab] OR swines[tiab] OR piglets[tiab] OR piglet[tiab] OR boar[tiab] OR boars[tiab] OR "sus scrofa"[tiab] OR ferrets[tiab] OR ferret[tiab] OR polecat[tiab] OR polecats[tiab] OR "mustela putorius"[tiab] OR "guinea pigs"[Tiab] OR "guinea pig"[Tiab] OR cavia[Tiab] OR callithrix[Tiab] OR marmoset[Tiab] OR marmosets[Tiab] OR cebuella[Tiab] OR hapale[Tiab] OR octodon[Tiab] OR chinchilla[Tiab] OR chinchillas[Tiab] OR gerbillinae[Tiab] OR gerbil[Tiab] OR gerbils[Tiab] OR jird[Tiab] OR jirds[Tiab] OR merione[Tiab] OR meriones[Tiab] OR rabbits[Tiab] OR rabbit[Tiab] OR hares[Tiab] OR hare[Tiab] OR diptera[Tiab] OR flies[Tiab] OR fly[Tiab] OR dipteral[Tiab] OR drosphila[Tiab] OR drosophilidae[Tiab] OR cats[Tiab] OR cat[Tiab] OR carus[Tiab] OR felis[Tiab] OR nematoda[Tiab] OR nematode[Tiab] OR nematoda[Tiab] OR nematode[Tiab] OR nematodes[Tiab] OR sipunculida[Tiab] OR dogs[Tiab] OR dog[Tiab] OR canine[Tiab] OR canines[Tiab] OR canis[Tiab] OR sheep[Tiab] OR sheeps[Tiab] OR mouflon[Tiab] OR mouflons[Tiab] OR ovis[Tiab] OR goats[Tiab] OR goat[Tiab] OR capra[Tiab] OR capras[Tiab] OR rupicapra[Tiab] OR chamois[Tiab] OR haplorhini[Tiab] OR monkey[Tiab] OR monkeys[Tiab] OR anthropoidea[Tiab] OR anthropoids[Tiab] OR saguinus[Tiab] OR tamarin[Tiab] OR tamarins[Tiab] OR leontopithecus[Tiab] OR hominidae[Tiab] OR ape[Tiab] OR apes[Tiab] OR pan[Tiab] OR paniscus[Tiab] OR "pan paniscus"[Tiab] OR bonobo[Tiab] OR bonobos[Tiab] OR troglodytes[Tiab] OR "pan troglodytes"[Tiab] OR gibbon[Tiab] OR gibbons[Tiab] OR siamang[Tiab] OR siamangs[Tiab] OR nomascus[Tiab] OR symphalangus[Tiab] OR chimpanzee[Tiab] OR chimpanzees[Tiab] OR prosimians[Tiab] OR "bush baby"[Tiab] OR prosimian[Tiab] OR bush babies[Tiab] OR galagos[Tiab] OR galago[Tiab] OR pongidae[Tiab] OR gorilla[Tiab] OR gorillas[Tiab] OR pongo[Tiab] OR pygmaeus[Tiab] OR "pongo pygmaeus"[Tiab] OR orangutans[Tiab] OR pygmaeus[Tiab] OR lemur[Tiab] OR lemurs[Tiab] OR lemuridae[Tiab] OR horse[Tiab] OR horses[Tiab] OR pongo[Tiab] OR equus[Tiab] OR cow[Tiab] OR calf[Tiab] OR bull[Tiab] OR chicken[Tiab] OR chickens[Tiab] OR gallus[Tiab] OR quail[Tiab] OR bird[Tiab] OR birds[Tiab] OR quails[Tiab] OR poultry[Tiab] OR poultries[Tiab] OR fowl[Tiab] OR fowls[Tiab] OR reptile[Tiab] OR reptilia[Tiab] OR reptiles[Tiab] OR snakes[Tiab] OR snake[Tiab] OR lizard[Tiab] OR lizards[Tiab] OR alligator[Tiab] OR alligators[Tiab] OR crocodile[Tiab] OR crocodiles[Tiab] OR turtle[Tiab] OR turtles[Tiab] OR amphibian[Tiab] OR amphibians[Tiab] OR amphibia[Tiab] OR frog[Tiab] OR frogs[Tiab] OR bombina[Tiab] OR salientia[Tiab] OR toad[Tiab] OR toads[Tiab] OR "epidalea calamita"[Tiab] OR salamander[Tiab] OR salamanders[Tiab] OR eel[Tiab] OR eels[Tiab] OR fish[Tiab] OR fishes[Tiab] OR pisces[Tiab] OR catfish[Tiab] OR catfishes[Tiab] OR siluriformes[Tiab] OR arius[Tiab] OR heteropneustes[Tiab] OR sheatfish[Tiab] OR perch[Tiab] OR perches[Tiab] OR percidae[Tiab] OR perca[Tiab] OR trout[Tiab] OR trouts[Tiab] OR char[Tiab] OR chars[Tiab] OR salvelinus[Tiab] OR "fathead minnow"[Tiab] OR minnow[Tiab] OR cyprinidae[Tiab] OR carps[Tiab] OR carp[Tiab] OR zebrafish[Tiab] OR zebrafishes[Tiab] OR goldfish[Tiab] OR goldfishes[Tiab] OR guppy[Tiab] OR guppies[Tiab] OR chub[Tiab] OR chubs[Tiab] OR tinca[Tiab] OR barbels[Tiab] OR barbus[Tiab] OR pimephales[Tiab] OR promelas[Tiab] OR "poecilia reticulata"[Tiab] OR mullet[Tiab] OR mullets[Tiab] OR seahorse[Tiab] OR seahorses[Tiab] OR mugil curema[Tiab] OR atlantic cod[Tiab] OR shark[Tiab] OR sharks[Tiab] OR catshark[Tiab] OR anguilla[Tiab] OR salmonid[Tiab] OR salmonids[Tiab] OR whitefish[Tiab] OR whitefishes[Tiab] OR salmon[Tiab] OR salmons[Tiab] OR sole[Tiab] OR solea[Tiab] OR "sea lamprey"[Tiab] OR lamprey[Tiab] OR lampreys[Tiab] OR pumpkinseed[Tiab] OR sunfish[Tiab] OR sunfishes[Tiab] OR tilapia[Tiab] OR tilapias[Tiab] OR turbot[Tiab] OR turbots[Tiab] OR flatfish[Tiab] OR flatfishes[Tiab] OR sciuridae[Tiab] OR squirrel[Tiab] OR squirrels[Tiab] OR chipmunk[Tiab] OR chipmunks[Tiab] OR suslik[Tiab] OR susliks[Tiab] OR vole[Tiab] OR voles[Tiab] OR lemming[Tiab] OR lemmings[Tiab] OR muskrat[Tiab] OR muskrats[Tiab] OR lemmus[Tiab] OR otter[Tiab] OR otters[Tiab] OR marten[Tiab] OR martens[Tiab] OR martes[Tiab] OR weasel[Tiab] OR badger[Tiab] OR badgers[Tiab] OR ermine[Tiab] OR mink[Tiab] OR minks[Tiab] OR sable[Tiab] OR sables[Tiab] OR gulo[Tiab] OR gulos[Tiab] OR wolverine[Tiab] OR wolverines[Tiab] OR minks[Tiab] OR mustela[Tiab] OR llama[Tiab] OR llamas[Tiab] OR alpaca[Tiab] OR alpacas[Tiab] OR camelid[Tiab] OR camelids[Tiab] OR guanaco[Tiab] OR guanacos[Tiab] OR chiroptera[Tiab] OR chiropteras[Tiab] OR bat[Tiab] OR bats[Tiab] OR fox[Tiab] OR foxes[Tiab] OR iguana[Tiab] OR iguanas[Tiab] OR xenopus laevis[Tiab] OR parakeet[Tiab] OR parakeets[Tiab] OR parrot[Tiab] OR parrots[Tiab] OR donkey[Tiab] OR donkeys[Tiab] OR mule[Tiab] OR mules[Tiab] OR zebra[Tiab] OR zebras[Tiab] OR shrew[Tiab] OR shrews[Tiab] OR bison[Tiab] OR bisons[Tiab] OR buffalo[Tiab] OR buffaloes[Tiab] OR deer[Tiab] OR deers[Tiab] OR bear[Tiab] OR bears[Tiab] OR panda[Tiab] OR pandas[Tiab] OR "wild hog"[Tiab] OR "wild boar"[Tiab] OR fitchew[Tiab] OR fitch[Tiab] OR beaver[Tiab] OR beavers[Tiab] OR jerboa[Tiab] OR jerboas[Tiab] OR capybara[Tiab] OR capybaras[Tiab]) NOT medline[subset])

**PsychINFO:**

pregnancy/ or antepartum period/ or neonatal period/ or neonatal development/ or perinatal period/ or postnatal period/ or prenatal exposure/ or prenatal development/ or prenatal developmental stages/ or embryo/ or fetus/ or neonatal development/ or animal development/ or development/ or adolescent development/ or infant development/ or (pregnan* or perinatal or peri-natal or prenatal or pre-natal or postnatal or post-natal or antepartum or ante-partum or (ante partum) or antepartal or prepartum or pre-partum or prepartal or pre-partal or neonatal or neo-natal or neonate* or adolescent* or adolescence or juvenile or newborn* or infant* or maternal-fetal or maternal-foetal or transplacental or (in utero) or gestation* or early-life or development* or developing or fetus* or foetus* or fetal or embryo* or embryo-genesis).ti,ab,id.

AND

exp serotonin reuptake inhibitors/ or sertraline/ or (SSRI* or (Selective adj3 Serotonin adj3 Uptake adj3 Inhibitor) or (Selective adj3 Serotonin adj3 Uptake adj3 Inhibitors) OR (Selective adj3 Serotonin adj3 Reuptake adj3 Inhibitor) or (Selective adj3 Serotonin adj3 Reuptake adj3 Inhibitors) or (Serotonin adj3 Uptake adj3 Inhibitor) or (Serotonin adj3 Uptake adj3 Inhibitors) or (Serotonin adj3 Reuptake adj3 Inhibitor) or (Serotonin adj3 Reuptake adj3 Inhibitors) or ( 5-Hydroxytryptamine adj3 Uptake adj3 Inhibitor) or ( 5-Hydroxytryptamine adj3 Uptake adj3 Inhibitors) or (5 adj3 Hydroxytryptamine adj3 Uptake adj3 Inhibitor) or (5 adj3 Hydroxytryptamine adj3 Uptake adj3 Inhibitors) or (5-HT adj3 Uptake adj3 Inhibitor) or (5-HT adj3 Uptake adj3 Inhibitors) or (5 adj3 HT adj3 Uptake adj3 Inhibitor) or (5 adj3 HT adj3 Uptake adj3 Inhibitors) or vilazodone or dapoxetine or Citalopram or femoxetine or Fluoxetine or Fluvoxamine or indalpine or Paroxetine or Sertraline or Zimeldine or dapoxetine or Citalopram or femoxetine or Fluoxetine or Fluvoxamine or indalpine or Paroxetine or Sertraline or Zimeldine or Priligy or Cytalopram or Escitalopram or Lexapro or femoxitine or Fluoxetin or Sarafem or Prozac or Fluvoxamin or Fluvoxamina or Luvox or Fevarin or Floxyfral or Dumirox or Faverin or Upstene or Seroxat or Paxil or Aropax or Zoloft or Lustral or Aremis or Besitran or Zimelidine or Zimelidin or Zelmid).ti,ab,id.

AND

animal research/ or animal models/ or animals/ or female animals/ or male animals/ or exp invertebrates/ or exp vertebrates/ or (animal OR animals OR pisces OR fish OR fishes OR catfish OR catfishes OR sheatfish OR silurus OR arius OR heteropneustes OR clarias OR gariepinus OR fathead minnow OR fathead minnows OR pimephales OR promelas OR cichlidae OR trout OR trouts OR char OR chars OR salvelinus OR salmo OR oncorhynchus OR guppy OR guppies OR millionfish OR poecilia OR goldfish OR goldfishes OR carassius OR auratus OR mullet OR mullets OR mugil OR curema OR shark OR sharks OR cod OR cods OR gadus OR morhua OR carp OR carps OR cyprinus OR carpio OR killifish OR eel OR eels OR anguilla OR zander OR sander OR lucioperca OR stizostedion OR turbot OR turbots OR psetta OR flatfish OR flatfishes OR plaice OR pleuronectes OR platessa OR tilapia OR tilapias OR oreochromis OR sarotherodon OR common sole OR dover sole OR solea OR zebrafish OR zebrafishes OR danio OR rerio OR seabass OR dicentrarchus OR labrax OR morone OR lamprey OR lampreys OR petromyzon OR pumpkinseed OR pumpkinseeds OR lepomis OR gibbosus OR herring OR clupea OR harengus OR amphibia OR amphibian OR amphibians OR anura OR salientia OR frog OR frogs OR rana OR toad OR toads OR bufo OR xenopus OR laevis OR bombina OR epidalea OR calamita OR salamander OR salamanders OR newt OR newts OR triturus OR reptilia OR reptile OR reptiles OR bearded dragon OR pogona OR vitticeps OR iguana OR iguanas OR lizard OR lizards OR anguis fragilis OR turtle OR turtles OR snakes OR snake OR aves OR bird OR birds OR quail OR quails OR coturnix OR bobwhite OR colinus OR virginianus OR poultry OR poultries OR fowl OR fowls OR chicken OR chickens OR gallus OR zebra finch OR taeniopygia OR guttata OR canary OR canaries OR serinus OR canaria OR parakeet OR parakeets OR grasskeet OR parrot OR parrots OR psittacine OR psittacines OR shelduck OR tadorna OR goose OR geese OR branta OR leucopsis OR woodlark OR lullula OR flycatcher OR ficedula OR hypoleuca OR dove OR doves OR geopelia OR cuneata OR duck OR ducks OR greylag OR graylag OR anser OR harrier OR circus pygargus OR red knot OR great knot OR calidris OR canutus OR godwit OR limosa OR lapponica OR meleagris OR gallopavo OR jackdaw OR corvus OR monedula OR ruff OR philomachus OR pugnax OR lapwing OR peewit OR plover OR vanellus OR swan OR cygnus OR columbianus OR bewickii OR gull OR chroicocephalus OR ridibundus OR albifrons OR great tit OR parus OR aythya OR fuligula OR streptopelia OR risoria OR spoonbill OR platalea OR leucorodia OR blackbird OR turdus OR merula OR blue tit OR cyanistes OR pigeon OR pigeons OR columba OR pintail OR anas OR starling OR sturnus OR owl OR athene noctua OR pochard OR ferina OR cockatiel OR nymphicus OR hollandicus OR skylark OR alauda OR tern OR sterna OR teal OR crecca OR oystercatcher OR haematopus OR ostralegus OR shrew OR shrews OR sorex OR araneus OR crocidura OR russula OR european mole OR talpa OR chiroptera OR bat OR bats OR eptesicus OR serotinus OR myotis OR dasycneme OR daubentonii OR pipistrelle OR pipistrellus OR cat OR cats OR felis OR catus OR feline OR dog OR dogs OR canis OR canine OR canines OR otter OR otters OR lutra OR badger OR badgers OR meles OR fitchew OR fitch OR foumart or foulmart OR ferrets OR ferret OR polecat OR polecats OR mustela OR putorius OR weasel OR weasels OR fox OR foxes OR vulpes OR common seal OR phoca OR vitulina OR grey seal OR halichoerus OR horse OR horses OR equus OR equine OR equidae OR donkey OR donkeys OR mule OR mules OR pig OR pigs OR swine OR swines OR hog OR hogs OR boar OR boars OR porcine OR piglet OR piglets OR sus OR scrofa OR llama OR llamas OR lama OR glama OR deer OR deers OR cervus OR elaphus OR cow OR cows OR bos taurus OR bos indicus OR bovine OR bull OR bulls OR cattle OR bison OR bisons OR sheep OR sheeps OR ovis aries OR ovine OR lamb OR lambs OR mouflon OR mouflons OR goat OR goats OR capra OR caprine OR chamois OR rupicapra OR leporidae OR lagomorpha OR lagomorph OR rabbit OR rabbits OR oryctolagus OR cuniculus OR laprine OR hares OR lepus OR rodentia OR rodent OR rodents OR murinae OR mouse OR mice OR mus OR musculus OR murine OR woodmouse OR apodemus OR rat OR rats OR rattus OR norvegicus OR guinea pig OR guinea pigs OR cavia OR porcellus OR hamster OR hamsters OR mesocricetus OR cricetulus OR cricetus OR gerbil OR gerbils OR jird OR jirds OR meriones OR unguiculatus OR jerboa OR jerboas OR jaculus OR chinchilla OR chinchillas OR beaver OR beavers OR castor fiber OR castor canadensis OR sciuridae OR squirrel OR squirrels OR sciurus OR chipmunk OR chipmunks OR marmot OR marmots OR marmota OR suslik OR susliks OR spermophilus OR cynomys OR cottonrat OR cottonrats OR sigmodon OR vole OR voles OR microtus OR myodes OR glareolus OR primate OR primates OR prosimian OR prosimians OR lemur OR lemurs OR lemuridae OR loris OR bush baby OR bush babies OR bushbaby OR bushbabies OR galago OR galagos OR anthropoidea OR anthropoids OR simian OR simians OR monkey OR monkeys OR marmoset OR marmosets OR callithrix OR cebuella OR tamarin OR tamarins OR saguinus OR leontopithecus OR squirrel monkey OR squirrel monkeys OR saimiri OR night monkey OR night monkeys OR owl monkey OR owl monkeys OR douroucoulis OR aotus OR spider monkey OR spider monkeys OR ateles OR baboon OR baboons OR papio OR rhesus monkey OR macaque OR macaca OR mulatta OR cynomolgus OR fascicularis OR green monkey OR green monkeys OR chlorocebus OR vervet OR vervets OR pygerythrus OR hominoidea OR ape OR apes OR hylobatidae OR gibbon OR gibbons OR siamang OR siamangs OR nomascus OR symphalangus OR hominidae OR orangutan OR orangutans OR pongo OR chimpanzee OR chimpanzees OR pan troglodytes OR bonobo OR bonobos OR pan paniscus OR gorilla OR gorillas OR troglodytes).ti,ab,id.

**Web of Science**

TS=(pregnan* or perinatal or peri-natal or prenatal or pre-natal or postnatal or post-natal or antepartum or ante-partum or “ante partum” or antepartal or prepartum or pre-partum or prepartal or pre-partal or neonatal or neonate* or adolescent* or adolescence or juvenile or newborn* or infant* or maternal-fetal or maternal-foetal or transplacental or “in utero” or gestation* or early-life or development* or developing or fetus* or foetus* or fetal or embryo* or embryo-genesis)

AND

TS=(SSRI or SSRIs or (Selective NEAR/3 Serotonin NEAR/3 Uptake NEAR/3 Inhibitor) or (Selective NEAR/3 Serotonin NEAR/3 Uptake NEAR/3 Inhibitors) OR (Selective NEAR/3 Serotonin NEAR/3 Reuptake NEAR/3 Inhibitor) or (Selective NEAR/3 Serotonin NEAR/3 Reuptake NEAR/3 Inhibitors) or (Serotonin NEAR/3 Uptake NEAR/3 Inhibitor) or (Serotonin NEAR/3 Uptake NEAR/3 Inhibitors) or (Serotonin NEAR/3 Reuptake NEAR/3 Inhibitor) or (Serotonin NEAR/3 Reuptake NEAR/3 Inhibitors) or ( 5-Hydroxytryptamine NEAR/3 Uptake NEAR/3 Inhibitor) or ( 5-Hydroxytryptamine NEAR/3 Uptake NEAR/3 Inhibitors) or (5 NEAR/3 Hydroxytryptamine NEAR/3 Uptake NEAR/3 Inhibitor) or (5 NEAR/3 Hydroxytryptamine NEAR/3 Uptake NEAR/3 Inhibitors) or (5-HT NEAR/3 Uptake NEAR/3 Inhibitor) or (5-HT NEAR/3 Uptake NEAR/3 Inhibitors) or (5 NEAR/3 HT NEAR/3 Uptake NEAR/3 Inhibitor) or (5 NEAR/3 HT NEAR/3 Uptake NEAR/3 Inhibitors) or vilazodone or dapoxetine or Citalopram or femoxetine or Fluoxetine or Fluvoxamine or indalpine or Paroxetine or Sertraline or Zimeldine or dapoxetine or Citalopram or femoxetine or Fluoxetine or Fluvoxamine or indalpine or Paroxetine or Sertraline or Zimeldine or Priligy or Cytalopram or Escitalopram or Lexapro or femoxitine or Fluoxetin or Sarafem or Prozac or Fluvoxamin or Fluvoxamina or Luvox or Fevarin or Floxyfral or Dumirox or Faverin or Upstene or Seroxat or Paxil or Aropax or Zoloft or Lustral or Aremis or Besitran or Zimelidine or Zimelidin or Zelmid)

AND

TS=(animal OR animals OR pisces OR fish OR fishes OR catfish OR catfishes OR sheatfish OR silurus OR arius OR heteropneustes OR clarias OR gariepinus OR “fathead minnow” OR “fathead minnows” OR pimephales OR promelas OR cichlidae OR trout OR trouts OR char OR chars OR salvelinus OR salmo OR oncorhynchus OR guppy OR guppies OR millionfish OR poecilia OR goldfish OR goldfishes OR carassius OR auratus OR mullet OR mullets OR mugil OR curema OR shark OR sharks OR cod OR cods OR gadus OR morhua OR carp OR carps OR cyprinus OR carpio OR killifish OR eel OR eels OR anguilla OR zander OR sander OR lucioperca OR stizostedion OR turbot OR turbots OR psetta OR flatfish OR flatfishes OR plaice OR pleuronectes OR platessa OR tilapia OR tilapias OR oreochromis OR sarotherodon OR “common sole” OR “dover sole” OR solea OR zebrafish OR zebrafishes OR danio OR rerio OR seabass OR dicentrarchus OR labrax OR morone OR lamprey OR lampreys OR petromyzon OR pumpkinseed OR pumpkinseeds OR lepomis OR gibbosus OR herring OR clupea OR harengus OR amphibia OR amphibian OR amphibians OR anura OR salientia OR frog OR frogs OR rana OR toad OR toads OR bufo OR xenopus OR laevis OR bombina OR epidalea OR calamita OR salamander OR salamanders OR newt OR newts OR triturus OR reptilia OR reptile OR reptiles OR “bearded dragon” OR pogona OR vitticeps OR iguana OR iguanas OR lizard OR lizards OR “anguis fragilis” OR turtle OR turtles OR snakes OR snake OR aves OR bird OR birds OR quail OR quails OR coturnix OR bobwhite OR colinus OR virginianus OR poultry OR poultries OR fowl OR fowls OR chicken OR chickens OR gallus OR “zebra finch” OR taeniopygia OR guttata OR canary OR canaries OR serinus OR canaria OR parakeet OR parakeets OR grasskeet OR parrot OR parrots OR psittacine OR psittacines OR shelduck OR tadorna OR goose OR geese OR branta OR leucopsis OR woodlark OR lullula OR flycatcher OR ficedula OR hypoleuca OR dove OR doves OR geopelia OR cuneata OR duck OR ducks OR greylag OR graylag OR anser OR harrier OR “circus pygargus” OR “red knot” OR “great knot” OR calidris OR canutus OR godwit OR limosa OR lapponica OR meleagris OR gallopavo OR jackdaw OR corvus OR monedula OR ruff OR philomachus OR pugnax OR lapwing OR peewit OR plover OR vanellus OR swan OR cygnus OR columbianus OR bewickii OR gull OR chroicocephalus OR ridibundus OR albifrons OR “great tit” OR parus OR aythya OR fuligula OR streptopelia OR risoria OR spoonbill OR platalea OR leucorodia OR blackbird OR turdus OR merula OR “blue tit” OR cyanistes OR pigeon OR pigeons OR columba OR pintail OR anas OR starling OR sturnus OR owl OR “athene noctua” OR pochard OR ferina OR cockatiel OR nymphicus OR hollandicus OR skylark OR alauda OR tern OR sterna OR teal OR crecca OR oystercatcher OR haematopus OR ostralegus OR shrew OR shrews OR sorex OR araneus OR crocidura OR russula OR “european mole” OR talpa OR chiroptera OR bat OR bats OR eptesicus OR serotinus OR myotis OR dasycneme OR daubentonii OR pipistrelle OR pipistrellus OR cat OR cats OR felis OR catus OR feline OR dog OR dogs OR canis OR canine OR canines OR otter OR otters OR lutra OR badger OR badgers OR meles OR fitchew OR fitch OR foumart or foulmart OR ferrets OR ferret OR polecat OR polecats OR mustela OR putorius OR weasel OR weasels OR fox OR foxes OR vulpes OR “common seal” OR phoca OR vitulina OR “grey seal” OR halichoerus OR horse OR horses OR equus OR equine OR equidae OR donkey OR donkeys OR mule OR mules OR pig OR pigs OR swine OR swines OR hog OR hogs OR boar OR boars OR porcine OR piglet OR piglets OR sus OR scrofa OR llama OR llamas OR lama OR glama OR deer OR deers OR cervus OR elaphus OR cow OR cows OR “bos Taurus” OR “bos indicus” OR bovine OR bull OR bulls OR cattle OR bison OR bisons OR sheep OR sheeps OR “ovis aries” OR ovine OR lamb OR lambs OR mouflon OR mouflons OR goat OR goats OR capra OR caprine OR chamois OR rupicapra OR leporidae OR lagomorpha OR lagomorph OR rabbit OR rabbits OR oryctolagus OR cuniculus OR laprine OR hares OR lepus OR rodentia OR rodent OR rodents OR murinae OR mouse OR mice OR mus OR musculus OR murine OR woodmouse OR apodemus OR rat OR rats OR rattus OR norvegicus OR “guinea pig” OR “guinea pigs” OR cavia OR porcellus OR hamster OR hamsters OR mesocricetus OR cricetulus OR cricetus OR gerbil OR gerbils OR jird OR jirds OR meriones OR unguiculatus OR jerboa OR jerboas OR jaculus OR chinchilla OR chinchillas OR beaver OR beavers OR “castor fiber” OR “castor Canadensis” OR sciuridae OR squirrel OR squirrels OR sciurus OR chipmunk OR chipmunks OR marmot OR marmots OR marmota OR suslik OR susliks OR spermophilus OR cynomys OR cottonrat OR cottonrats OR sigmodon OR vole OR voles OR microtus OR myodes OR glareolus OR primate OR primates OR prosimian OR prosimians OR lemur OR lemurs OR lemuridae OR loris OR “bush baby” OR “bush babies” OR bushbaby OR bushbabies OR galago OR galagos OR anthropoidea OR anthropoids OR simian OR simians OR monkey OR monkeys OR marmoset OR marmosets OR callithrix OR cebuella OR tamarin OR tamarins OR saguinus OR leontopithecus OR “squirrel monkey” OR “squirrel monkeys” OR saimiri OR “night monkey” OR “night monkeys” OR “owl monkey” OR “owl monkeys” OR douroucoulis OR aotus OR “spider monkey” OR “spider monkeys” OR ateles OR baboon OR baboons OR papio OR “rhesus monkey” OR macaque OR macaca OR mulatta OR cynomolgus OR fascicularis OR “green monkey” OR “green monkeys” OR chlorocebus OR vervet OR vervets OR pygerythrus OR hominoidea OR ape OR apes OR hylobatidae OR gibbon OR gibbons OR siamang OR siamangs OR nomascus OR symphalangus OR hominidae OR orangutan OR orangutans OR pongo OR chimpanzee OR chimpanzees OR “pan troglodytes” OR bonobo OR bonobos OR “pan paniscus” OR gorilla OR gorillas OR troglodytes)
