## Supplementary File 2 for "Perinatal selective serotonin reuptake inhibitor exposure and behavioral outcomes: a systematic review and meta-analyses of animal studies"

| **Category** | **Behavioral test** | **Primary outcome** | **Alternative outcome #1** | **Alternative outcome #2** | **Alternative outcome #3** | **Alternative outcome #4** | **Alternative outcome #5** |
| --- | --- | --- | --- | --- | --- | --- | --- |
| **Activity & exploration** | Open field test | Distance moved | Ambulation (crossings) | Duration ambulation |  |  |  |
|  | Home cage activity | Total activity (beam breaks, seconds or distance) |  |  |  |  |  |
|  | Novel object exploration | Exploration time object | Entries to target zone |  |  |  |  |
|  | Voluntary running wheel | Running wheel time/counts |  |  |  |  |  |
|  | 8-arm maze | No. of novel arm choices |  |  |  |  |  |
|  | Elevated plus maze | Distance moved |  |  |  |  |  |
|  | Object directed behavior | Distance moved |  |  |  |  |  |
| **Anxiety-like behavior** | Open field test | Time in center | Thigmotaxis time | Distance in center | Entries in center | Latency to enter field | Defecation score |
|  | Elevated plus maze | Time on open arms | Time on closed arms | Latency to open arms |  |  |  |
|  | Novelty suppressed feeding | Latency to feed |  |  |  |  |  |
|  | Elevated zero maze | Time in open quadrants |  |  |  |  |  |
|  | Light-dark test | Time in light chamber |  |  |  |  |  |
|  | Marble burying test | No. marbles buried |  |  |  |  |  |
|  | Response to (long) tone | Immobility during tone | Distance moved during tone |  |  |  |  |
|  | Defensive withdrawal | Latency to exit |  |  |  |  |  |
| **Stress coping** | Forced swim test | Immobility duration test 2 | Immobility duration test 1 | Immobility counts | Immobility latency | Duration struggling |  |
|  | Shock avoidance | Escape latency |  |  |  |  |  |
|  | Tail suspension test | Immobility duration |  |  |  |  |  |
|  | Learned helplessness | Escape latency |  |  |  |  |  |
|  | EPM after stress | Time on open arms |  |  |  |  |  |
|  | Open field after stress | Time in center |  |  |  |  |  |
|  | Marble burying after stress | No. marbles buried |  |  |  |  |  |
|  | Acoustic startle after stress | % habituation |  |  |  |  |  |
| **Social behavior** | Copulatory behavior (male) | Number of ejaculations | Number of intromissions | Number of mounts |  |  |  |
|  | Aggression | Time in aggressive behavior | Attack latency | % animals aggressive |  |  |  |
|  | Resident-intruder test | Duration attack | Percentage of time aggression |  |  |  |  |
|  | Social play test | Time total play behavior | Frequency total play behavior | Duration no social behavior | Duration pinning | Duration pouncing |  |
|  | Social interaction test | Time total social behavior | Frequency total social behavior |  |  |  |  |
|  | Object-conspecific preference | Preference object:animal |  |  |  |  |  |
|  | Social preference test | Preference stranger:familiar | Preference stranger:empty | Duration social interaction |  |  |  |
|  | Copulatory behavior (female) | Frequency proceptive behaviors | Lordosis quotient |  |  |  |  |
|  | Pup retrieval | Time to retrieve first pup |  |  |  |  |  |
|  | Sexual incentive motivation | Preference sexual:social |  |  |  |  |  |
|  | Ultrasonic vocalizations | Number of vocalizations | Presence of vocalizations |  |  |  |  |
|  | Maternal behavior | Time to retrieve pups |  |  |  |  |  |
|  | Nest quality | Nest quality score |  |  |  |  |  |
| **Learning & memory** | Morris water maze | Time in target quadrant | Crosses target quadrant | Av. distance from target | Time to escape |  |  |
|  | Passive avoidance test | Time to enter dark comptmnt |  |  |  |  |  |
|  | Cincinnati water maze | Latency to escape | Number of errors |  |  |  |  |
|  | Cued fear conditioning | Freezing during tone |  |  |  |  |  |
|  | Novel object recognition | Exploration novel:familiar | Duration investigation novel object |  |  |  |  |
|  | Contextual fear conditioning | Freezing |  |  |  |  |  |
|  | Scent recognition | Contact novel scent |  |  |  |  |  |
|  | Complex maze | No. correct trials |  |  |  |  |  |
|  | Radial water maze | No. errors |  |  |  |  |  |
|  | Barnes maze | Time to escape |  |  |  |  |  |
| **Ingestive & reward** | Food intake | Food intake (g or calories) |  |  |  |  |  |
|  | Sucrose preference test | % sucrose preference |  |  |  |  |  |
|  | Alcohol consumption | % of total fluid intake |  |  |  |  |  |
|  | Sucrose consumption | Sucrose consumption (g) |  |  |  |  |  |
|  | Cocaine reward sensitivity | Time in cocaine-paired compartment vs baseline | Number of injections |  |  |  |  |
|  | Tube runway | Latency to reach food |  |  |  |  |  |
| **Motoric** | Swimming | Latency to reach platform | Swimming score |  |  |  |  |
|  | Horizontal ladder test | Time to cross |  |  |  |  |  |
|  | Rotarod test | Latency to fall |  |  |  |  |  |
|  | Running | Time to approach food/water |  |  |  |  |  |
|  | Bar holding | Latency to fall |  |  |  |  |  |
|  | Beam traversing | Time to traverse |  |  |  |  |  |
|  | Grooming behavior | Grooming duration |  |  |  |  |  |
|  | Walking (shaking/instability) | Stability score |  |  |  |  |  |
| **Sensory processing** | Prepulse inhibition/acoustic startle | % PPI (83%) | % habituation |  |  |  |  |
|  | Auditory test | Discrimination threshold |  |  |  |  |  |
|  | Gap crossing test | Max. crossable distance |  |  |  |  |  |
|  | Olfactory investigation | Contact new:old scent |  |  |  |  |  |
| **Reflex & pain** | Negative geotaxis | Time to turn | Turning score | PND reflex appearance |  |  |  |
|  | Righting reflex | Time to turn | Turning score | PND reflex appearance |  |  |  |
|  | Thermal sensitivity | Time to response |  |  |  |  |  |
|  | Vibrissa-placing | PND reflex appearance |  |  |  |  |  |
|  | Acoustic startle reflex | PND reflex appearance |  |  |  |  |  |
|  | Cliff avoidance | PND reflex appearance |  |  |  |  |  |
|  | Free-fall righting | PND reflex appearance |  |  |  |  |  |
|  | Palmar grasp | PND reflex appearance |  |  |  |  |  |
|  | Angle of fall | Maximum angle |  |  |  |  |  |
|  | Gait reflex | Time to crawl out of circle |  |  |  |  |  |
|  | Mechanical sensitivity | Paw withdraw threshold |  |  |  |  |  |
| **Sleep & circadian** | Sleep-wake behavior | % wakefulness | % REM sleep |  |  |  |  |
|  | Phase shifts to pulse | Phase shift |  |  |  |  |  |
|  | Phase angle LD | Phase angle |  |  |  |  |  |
|  | Entrainment to cycle advance | Days to entrain |  |  |  |  |  |
|  | Hours of activity LL DD LD | Alpha |  |  |  |  |  |
|  | Free-running period | Tau |  |  |  |  |  |
